# Multiomics unveils extracellular vesicle-driven mechanisms of endothelial communication in human carotid atherosclerosis

**DOI:** 10.1101/2024.06.21.599781

**Authors:** Sneha Raju, Mandy E Turner, Christian Cao, Majed Abdul-Samad, Neil Punwasi, Mark C Blaser, Rachel ME Cahalane, Steven R Botts, Kamalben Prajapati, Sarvatit Patel, Ruilin Wu, Dakota Gustafson, Natalie J Galant, Lindsey Fiddes, Melody Chemaly, Ulf Hedin, Ljubica Matic, Michael Seidman, Vallijah Subasri, Sasha A Singh, Elena Aikawa, Jason E Fish, Kathryn L Howe

**Affiliations:** Toronto General Hospital Research Institute, University Health Network, Toronto, Canada; Institute of Medical Science, University of Toronto, Toronto, Canada; Division of Vascular Surgery, Toronto General Hospital, Toronto, Canada; Faculty of Medicine, University of Toronto, Toronto ON, Canada; Center for Interdisciplinary Cardiovascular Sciences, Cardiovascular Division, Department of Medicine, Brigham and Women’s Hospital, Harvard Medical School, Boston, MA, USA; Department of Laboratory Medicine and Pathobiology, University of Toronto, Toronto, ON, Canada; Peter Munk Cardiac Centre, Toronto General Hospital, Toronto, Canada; Mechanobiology and Medical Device Research group (MMDRG), Biomedical Engineering, College of Science and Engineering, University of Galway, Galway, Ireland; Princess Margaret Cancer Center, Toronto, ON, Canada; Vascular Surgery Division, Department of Molecular Medicine and Surgery, Karolinska University Hospital and Karolinska Institut, Stockholm, Sweden; Center for Excellence in Vascular Biology, Brigham and Women’s Hospital, Harvard Medical School, Boston, MA, USA

**Keywords:** carotid atherosclerosis, extracellular vesicles, microRNAs, proteomics, endothelial cells

## Abstract

**Background:** Carotid atherosclerosis is orchestrated by cell-cell communication that drives progression along a clinical continuum (asymptomatic to symptomatic). Extracellular vesicles (EVs) are cell-derived nanoparticles representing a new paradigm in cellular communication. Little is known about their biological cargo, cellular origin/destination, and functional roles in human atherosclerotic plaque.

**Methods:** EVs were enriched via size exclusion chromatography from human carotid endarterectomy samples dissected into paired plaque and marginal zones (symptomatic n=16, asymptomatic n=13). EV cargos were assessed via whole transcriptome miRNA sequencing and mass spectrometry-based proteomics. EV multi-omics were integrated with bulk and single cell RNA-sequencing (scRNA-seq) datasets to predict EV cellular origin and ligand-receptor interactions, and multi-modal biological network integration of EV-cargo was completed. EV functional impact was assessed with endothelial angiogenesis assays.

**Results:** Carotid plaques contained more EVs than adjacent marginal zones, with differential enrichment for EV-miRNAs and EV-proteins in key atherogenic pathways. EV cellular origin analysis suggested that tissue EV signatures originated from endothelial cells (EC), smooth muscle cells (SMC), and immune cells. Integrated tissue vesiculomics and scRNA-seq indicated complex EV-vascular cell communication that changed with disease progression and plaque vulnerability (i.e., symptomatic disease). Plaques from symptomatic patients, but not asymptomatic patients, were characterized by increased involvement of endothelial pathways and more complex ligand-receptor interactions, relative to their marginal zones. Plaque-EVs were predicted to mediate communication with ECs. Pathway enrichment analysis delineated an endothelial signature with roles in angiogenesis and neovascularization – well-known indices of plaque instability. This was validated functionally, wherein human carotid symptomatic plaque EVs induced sprouting angiogenesis in comparison to their matched marginal zones.

**Conclusion:** Our findings indicate that EVs may drive dynamic changes in plaques through EV- vascular cell communication and effector functions that typify vulnerability to rupture, precipitating symptomatic disease. The discovery of endothelial-directed angiogenic processes mediated by EVs creates new therapeutic avenues for atherosclerosis.

## Introduction

Atherosclerosis is a multi-cellular biological process of lipid accumulation, unresolved inflammation, and plaque development in the arterial wall.^1,2^ While some plaques enlarge and impede blood flow, plaques can also become unstable and rupture, thereby releasing atheroembolic material that occludes an end-artery leading to a transient ischemic attack (TIA, i.e. neurologic recovery) or stroke.^1,3^ Atherosclerosis exists along a disease continuum wherein much of the disease process is silent and presents as asymptomatic disease; low-density lipoprotein accumulation and endothelial dysfunction initiates the disease process, followed by recruitment of adaptive and innate immune cells and migration of smooth muscle cells (SMCs) leading to plaque progression and development of the necrotic core, followed by eventual fibrous cap rupture and thrombosis causing clinical events such as myocardial infarction or stroke.^2^ Altered cellular communication is a critical linchpin of atherogenesis.^4–7^ Recent work has identified the importance of endothelial cells (ECs) in directing differential, polarized messaging with cells found in circulation and those in the vessel wall.^8^ Given that several types of plaque cells exist in transitory states (ECs, SMCs, macrophages), are highly dynamic, and can alter their phenotype in the active plaque microenvironment, an exciting possibility is that cellular states dictate and result from changes in cell-cell communication^9–12^

Much of the early and late stages of atherosclerotic disease is silent or asymptomatic, creating challenges in patient stratification. Unlike management of symptomatic carotid atherosclerotic disease, which is well established and includes surgical removal of the plaque (carotid endarterectomy) or endovascular carotid stenting with the goal of secondary stroke prevention ^13–16^, management of patients with asymptomatic carotid disease is more nuanced, as interventions have an upfront risk of stroke and we have not yet solved how to accurately predict plaque rupture.^17,18^ Current non-operative management with lipid reduction, anti- hypertensive therapy, and lifestyle modifications (i.e. smoking cessation) reduces stroke risk, but some patients progress despite meeting these targets.^17,19^ Prior studies have employed whole-tissue proteomics, transcriptomics, and metabolomics to identify major shifts in plaque biology with disease progression.^20–23^ However, despite ongoing research, there are still no universally reliable means to identify the patient at risk and novel therapeutic targets need to be unveiled through a better understanding of plaque biology as it evolves along the disease continuum.

Given the multi-cellular environment of the atherosclerotic plaque, cell-cell communication is a major factor driving the biology of the disease. Extracellular vesicles (EVs) are lipid bilayered membrane-enclosed cell-derived nanoparticles that carry bioactive molecules such as microRNAs (miRNAs) and proteins from a donor to a recipient cell, where they participate in EV- ligand based interactions or are endocytosed and release contents that can modulate recipient cell phenotype.^24^ Given the clinical potential of this field, the development of Minimal Information for the Study of Extracellular Vesicles (MISEV) guidelines has been crucial for improving clarity, consistency in reporting, and interpretation of EV-based studies.^25,26^ Previous studies in atherosclerotic plaques have examined larger EV particles (i.e., microparticles; primarily >200 nm) and provided early insights into their importance in atherosclerotic biology.^27,28^ However the evolution of EV isolation and technical progress in EV cargo analysis mandates a fresh examination of their biological roles, particularly with respect to small EVs (<200 nm) given their abundant numbers and potential for therapeutic modulation.^29,30^ Recently, a proteomics study of human carotid plaque-derived small and large EVs identified their role in advanced atherosclerotic calcification, and found that EV-proteomics can identify additional phenotypes than those at the whole plaque level, warranting independent investigation.^31^ Our understanding of carotid tissue vulnerability as modulated by EV cellular source, EV-cell communication, and the impact of EVs on downstream cellular processes is limited. There is a need to understand the holistic biological niche of stable and vulnerable carotid plaques in these contexts, with the goal of identifying novel targets for preventing rupture through plaque stabilization.

The intraplaque environment is dynamic, changes over time, and there is a significant role for cell-cell communication. EVs contain cargo that serve as important mediators of cell-cell communication with functional effects that alter cell biology in recipient cells.^32–34^ What is not known however, are the alterations to cell-cell communication that render an atherosclerotic plaque vulnerable to rupture. Patients with symptomatic carotid disease have demonstrated clinical consequences of carotid plaque vulnerability where atheroembolic events have caused transient ischemic events or stroke.^35^ Here, we performed a novel analysis to uncover the transcriptomic and proteomic EV profile in carotid plaque and adjacent marginal zones, in both asymptomatic and symptomatic disease states. By integrating our data with scRNA-seq datasets, we identified the predicted EV-cell communication pathways that may be enriched in asymptomatic and symptomatic (vulnerable) carotid plaques. Furthermore, we found a strong endothelial signature and demonstrated a functional role for plaque EVs in regulating angiogenesis and neovascularization – well-known precursors of plaque instability.^36–38^ Importantly, symptomatic plaques were studied within 30 days of the last neurologic event, with pathological evaluation of a subset that clearly demonstrated intraplaque hemorrhage, consistent with the EV angiogenesis experiments pursued. Together, our data suggest that EVs may contribute to dynamic changes seen in plaques through EV-cell communication and effector functions, identifying a mechanism for the neovascularization and angiogenesis that contribute to plaque destabilization. The discovery of an endothelial-centric process that could be mediated by EVs points towards novel therapeutics targeting this highly accessible and strategic cellular layer.

## Methods

Detailed methods are provided in the Supplemental Material. Human carotid endarterectomy specimens were obtained from patients undergoing surgery for asymptomatic and symptomatic disease. Samples were handled under the institutional research ethics board protocols set by University Health Network, Toronto Canada (18-5282). Patients gave written informed consent and procedures were followed in accordance with institutional guidelines. Tissues were separated by plaque and marginal (non-atheromatous) zones and EV enrichment performed according to MISEV2023 guidelines. Transcriptomics (next-generation miRNA sequencing) and proteomics (mass spectrometry-based) data are deposited in GEO and ProteomeXChange Consortium via the PRIDE repository (publicly available at the time of publication), respectively. Cell-cell communication was evaluated by integrating data with the TISSUES database^39^ and a publicly available scRNA-sequencing dataset of carotid endarterectomy specimens containing plaque and marginal zones (GSE159677).^20^ Multi-omic network analysis was performed using BIONIC.^40^ Functional validation of predicted roles for EVs on angiogenesis was performed in Human Umbilical Vein Endothelial Cell (HUVEC) spheroid-based sprouting assays.

### Statistical Analysis

The Fisher’s exact test tested the prevalence of gene ontology (GO) terms in whole tissue proteomics. For non-omics data with sample sizes <10, a two-sided Mann-Whitney or Wilcoxon (for paired samples) test was completed and for n≥10, data distribution was assessed and differences between two groups were tested using a two-tailed unpaired Student’s *t*-test. For comparisons between three or more groups, a one-way ANOVA was used with Šidák multiple comparisons test. Differences were significant for values of p<0.05 and are indicated in the figures. Proteomics (tissue and EV) were quantified using Discover (Version 2.5) and peptides were filtered based on a 1.0% FDR. A minimum of at least 2 unique peptides for each protein was required for the protein to be included in the analyses. Differential enrichment analysis was performed using Qlucore Omics Explorer statistical software (Version 3.9; Qlucore, Sweden) and differentially enriched proteins were calculated using a two-group comparison with a Benjamini-Hochberg false discovery rate (FDR)-corrected p-value (i.e., q-value) <0.05.

Transcriptomic data (miRNA) was quantified in Partek Flow (v10.0.23.0326) including trimming of adaptors, sequence alignment to genome (h38 assembly, Homo Sapiens, BioProject PRJNA31257, Bowtie), and quantification (miRBase mature miRNAs, Version 22). Differential expression of miRNA in experimental conditions was determined via DESeq2 with adjustment for multiple comparisons using Benjamini and Hochberg’s FDR step-up (adjusted p-value<0.05). Predicted messenger RNA (mRNA) targets of differentially expressed EV-miRNA were identified using MicroRNA Enrichment Turned Network (MIENTURNET version 2019-11-25) using the miRTarBase database. Pathway analysis of differentially enriched EV-miRNAs and EV-proteins were completed using Enrichr using the Gene Ontology (Biological Process and Cellular Component) and Kyoto Encyclopedia of Genes and Genomes (KEGG) databases with multiple testing correction using the Benjamini-Hochberg procedure (adjusted p<0.05). Protein-protein interaction analysis was completed in STRING v12.0. The TISSUES and Tabula Sapiens databases were accessed through STRING v12.0 and Enrich, respectively. EV cellular identity enrichement analysis was completed using the AddModule score function in Seurat (Version 5.0.1) to assess the enrichment of EV protein signatures among different cell types. The ‘CellChat’ R package (version 1.6.0) was used to infer cell interaction networks by filtering the CellChat cell-cell communication ligand database for genes associated with EV protein gene IDs. Comparison of fold enrichments between cohorts regarding endothelial-related GO biological processes were performed using ANOVA and Tukey’s honest significant difference.

## Results

### Atherosclerotic tissue proteome and extracellular vesicle (EV) concentration identify a potential role for EVs in modulating the disease continuum

As recently shown by Alsaigh et al., single cell RNA sequencing (scRNA-seq) of matched human carotid marginal zones and plaque zones harbour different cellular subpopulations, revealing distinct microenvironments, and cellular communication strategies that may provide insights into disease progression.^20^ To begin understanding the continuum of disease in carotid atherosclerosis, we delineated EV-based biology in plaque versus marginal zones from all patients (asymptomatic and symptomatic). Human carotid endarterectomy specimens for whole tissue proteomics (n=5) were dissected into plaque zones (PZ) and marginal zones (MZ) based on gross morphology by a single trained surgeon (Dr. S. Raju), with another 59 paired samples similarly dissected for EV enrichment and characterization (Figure 1A). Tissue histopathology with hematoxylin and eosin and Movat pentachrome stains delineated plaque and marginal zones to be different with distinguishing features of advanced atherosclerotic plaque morphology (e.g. necrotic core, intraplaque hemorrhage) visualized exclusively in plaque regions (Figure S1A-B).

**Figure 1.**
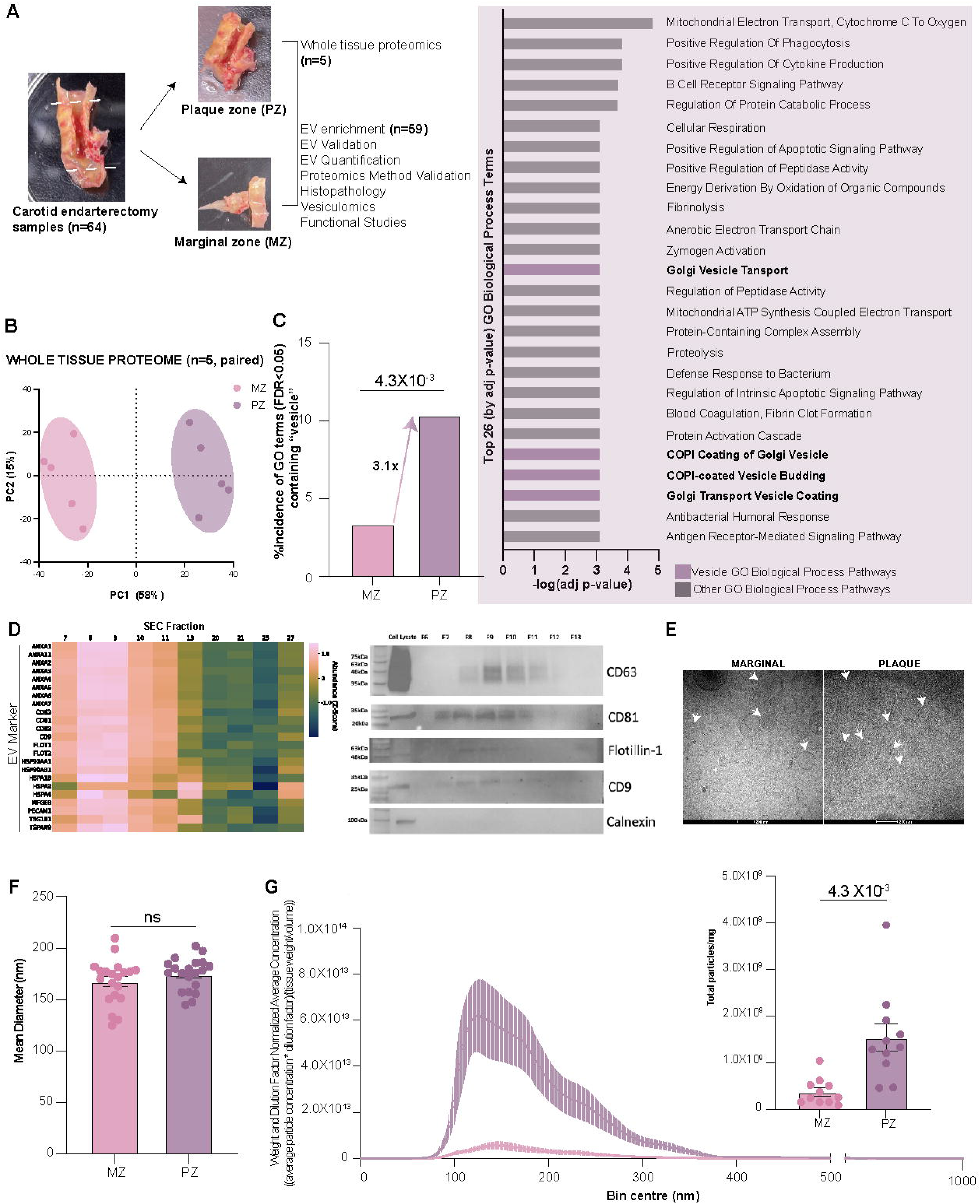
Whole tissue proteomics identifies a potential role for EVs in modulating plaque biology corroborated with increased enrichment of tissue resident EVs from human atherosclerotic carotid plaques. A. Experimental overview. Carotid endarterectomy specimens (*n=*64) were collected from the operating room and dissected into plaque and marginal zones for downstream analysis. B. Principal component analysis of carotid tissue proteomes between the PZ and MZ (donor matched *n*=5; donor uncorrected in Figure S2C). C. Percent incidence of Gene Ontology (GO; Biological Process and Cellular Component) terms (false discovery rate <0.05) containing “vesicle” derived from differentially enriched proteins (q- value<0.05) from plaque and marginal zones (left panel; *n*=5). Right panel lists top 26 (by adjusted p-value) GO Biological Process terms where vesicle-associated terms from plaque regions are highlighted (marginal zone in Figure S2D). D. Heatmap representing per-fraction proteomics identifies prototypical EV markers enriched in fractions 7-10 (left panel). Protein abundance was Z-score normalized by protein (Representative sample, *n*=4 in Figure S5A). Western blot analysis depicting EV markers (CD63, CD81, CD9, and Flot1) in EV-enriched fractions (7-10) isolated from human atherosclerotic plaque tissue (right panel, *n*=1). E. Cryo- EM of EVs isolated from plaque and adjacent marginal zones. Arrows indicate select EV structures as bilayered nanoparticles with dense cores (Representative Sample, *n*=4). Scale bar = 200nm. F. Quantification of EV mean diameter by nanoparticle tracking analysis (NTA; *n*=20). G. Nanoparticle concentration (normalized to tissue weight, NTA dilution factor, and total EV- enriched volume; *n=*11) binned by particle size from plaque (PZ, purple) and marginal zones (MZ, pink). Quantification of nanoparticle concentration across all sizes by NTA (*n*=11, right panel). Bar graphs show mean ± SEM. Statistical significance assessed by Fisher’s exact test (C) and a two-tailed paired t-test (11-20 pairs; F,G).

Tissue proteomes (Figure S2A) from the plaque zones and marginal zones clustered separately by principal component analysis (Figure 1B donor corrected; Figure S2C donor uncorrected), with 514 differentially enriched proteins identified by zone (Figure S2B). Pathway analysis of differentially enriched proteins in the marginal and plaque zone (padj<0.05) using Gene Ontology (GO) and Kyoto Encyclopedia of Genes and Genomes (KEGG) revealed atherogenic signatures, while GO terms containing ‘vesicle’ were increased in plaque zones specifically (Figure 1C, S2D-F, S3A-B). We next pursued EV enrichment in plaque and marginal zones separately. EVs were enriched using size exclusion chromatography that included an additional density ultracentrifugation step to remove extracellular matrix co-isolation^31^ and was validated in accordance with the MISEV2023 criteria (Figure 1D-G, S4A-F, S5A, S6). Briefly, EVs were enriched in fractions 7-10 as evidenced by single vesicle analysis using nanoparticle tracking analysis (NTA) and as a protein peak using the bicinchoninic acid assay (Figure S4E). The secondary protein peak (Figure S4E) represents smaller proteins that elute in the later fractions.^41^ The presence of common EV protein markers in per-fraction proteomics, including CD63, CD81, CD9, FLOT1, and an absence of the cell protein marker calnexin confirmed successful EV isolation in our fractions of interest (Figure 1D, left and right panels respectively; Figure S5A). Several vesicle-related pathways were enriched in plaque zones compared to marginal zones (Figure S5B), which may suggest increased EV processes taking place. We confirmed EV morphology via direct visualization using cryogenic electron microscopy (cryo-EM), which revealed bilayered nanoparticles with dense cores, as well as several muti- compartmented vesicles, recently seen to be released by cells (Figure 1E).^42^ We did not observe any morphological or size differences between EV isolates from plaque and marginal zones (Figure 1F). Paired plaque regions liberated similar EV size distribution, but a larger quantity of EVs per milligram of tissue relative to marginal zones (Figure 1G). Notably, we found minimal lipoprotein (APOA1^43^ and APOB100, enzyme-linked immunosorbent assay, Figure S4C- D) and no whole-cell (flow cytometry, Figure S4F) inclusion in vesicle preparations. Given EVs in marginal and plaque zones have the potential to contribute to the local milieu, we next compared the biological cargo of EVs between these two anatomical locations to help delineate the potential functional effects of the EV-vesiculome in carotid atherosclerotic disease progression.

### Carotid plaque EV-vesiculome (miRNA transcriptome and proteome) is distinct in plaque versus marginal zones and drives pathways known to govern plaque progression

Given that EV miRNA and protein cargo can mediate their functional effects on recipient cells in local environments,^32–34^ we performed next-generation miRNA sequencing and proteomic analysis of EVs enriched from carotid plaque zones and marginal zones (Figures S6, S7).

Exploratory analysis with donor-corrected principal component analysis (PCA) revealed that the EV-vesiculome clustered by location; specifically plaque zone (PZ, purple) and marginal zones (MZ, pink; Figure 2A-B; donor uncorrected Figure S7F). Differentially enriched protein pathway analysis predicted biological interconnectability in plaque-derived EVs, as seen by greater than expected protein-protein interactions (Figure 2C). Differential expression analysis between EV-miRNA in plaque versus marginal zones revealed 371 EV-miRNA enriched in plaque zones and 290 EV-miRNA enriched in marginal zones, across 20 paired patient samples (Figure 2D). Top differentially expressed upregulated miRNA in the plaque zones included miR-146a, miR-155, let-7a, miR-200b, miR-223, and miR181b, known to have roles in secretion and response to inflammatory mediators and cell proliferation and differentiation.^44–54^ Among differentially enriched proteins, we found 282 proteins enriched in plaque zones, and 379 enriched in marginal zones, across 23 paired patient samples (Figure 2E). Like the EV-miRNAs, several known mediators of plaque progression including metalloproteinases (MMP9), cell adhesion proteins (ICAM1), lipoprotein metabolism proteins (LPL), complement and coagulation proteins (F2, F10), and inflammatory mediators (CRP, SAA2-4, BLVRA, BLVRB) were more abundant in EVs from plaque environments (Figure 2E).

**Figure 2.**
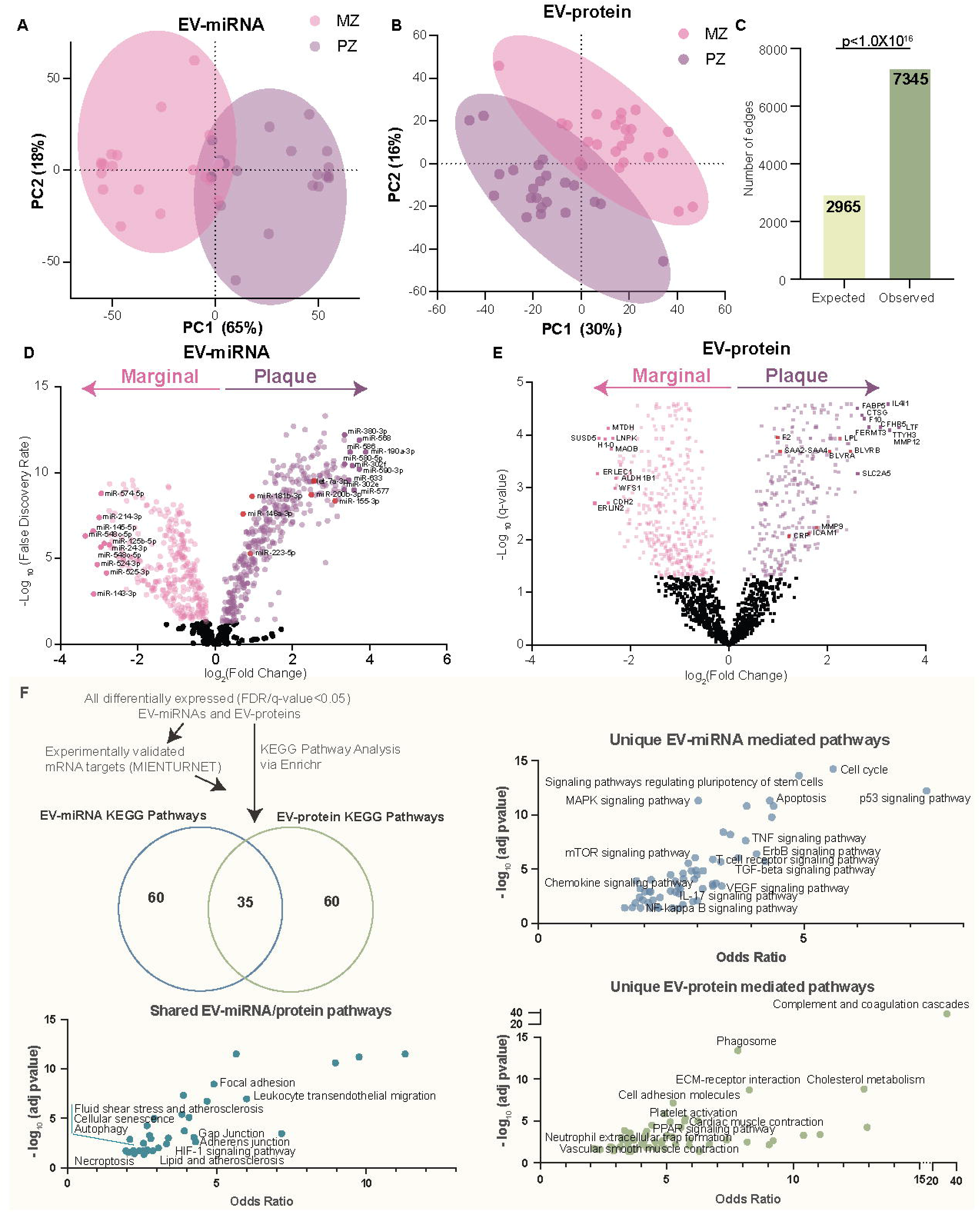
Carotid plaque EV-vesiculome is zone-specific (plaque versus marginal) with differentially expressed EV-miRNA and proteins associated with pro-atherogenic functions. A-B. Donor corrected principal component analysis (PCA) showing EV-miRNA **(A)** and EV-protein **(B)** profiles of EVs enriched from carotid plaque zones (PZ, purple) versus marginal zones (MZ, pink) (miRNA *n*=20 pairs; protein *n=*23 pairs). **C.** Edge quantification of protein-protein interaction network with all differentially enriched EV-proteins (n=661) in plaque versus marginal zones. Represents full STRING network wherein edges indicate both functional and physical protein interactions with a medium confidence interaction score and nodes represent proteins. **D-E.** Volcano plot of carotid tissue derived EV-miRNA **(D)** and EV-protein **(E)** contents with positive and negative fold change (FC) representing miRNAs and proteins enriched in plaque versus marginal zones, respectively (FDR step up/q-value<0.05). Highlighted in red are miRNAs and proteins known to function in inflammation. **F.** Kyoto Encyclopedia of Genes and Genomes (KEGG) pathway analysis of all differentially expressed (FDR/q-value <0.05) predicted EV-miRNA targets (mRNA, miRTarBase via MIENTURNET) and EV-proteins. Venn diagram of unique and shared KEGG pathways between predicted EV-miRNA targets and EV-proteins (top left panel). Bubble plots of KEGG pathways that were significantly (FDR<0.05) modulated by all differentially enriched plaque EV-miRNAs (top right panel) and EV-proteins (bottom right panel), with all pathways that were shared by EV-miRNA and EV-proteins shown in bottom left panel. Labeled are pathways of interest (FDR<0.05). Cancer-, and infection- associated pathways were excluded from analysis.

To explore the potential functions of EV-miRNA and protein cargo, we queried KEGG pathways using the Enrichr interface comparing plaque zone versus marginal zone EV cargo.^55^ Carotid plaque EV-cargo represented 155 unique KEGG pathways (FDR<0.05), with 60 pathways specific to EV-miRNAs and 60 pathways specific to EV-proteins (Figure 2F, Venn diagram). Top unique pathways predicted to be modulated by plaque EV-miRNA included cell cycle, apoptosis, and inflammatory signaling pathways (TNF, T cell receptor, TGF-β, chemokine, NFkB; Figure 2F, top right panel). Conversely, top unique pathways of differentially expressed plaque EV-proteins include complement and coagulation, phagosome, cholesterol metabolism, cell adhesion, and vascular smooth muscle contraction (Figure 2F, bottom right panel). Key pathways involved in atherosclerotic plaque progression shared between the EV-miRNA and EV-proteome included cellular senescence, cell adhesion and migration, leukocyte transendothelial migration, fluid shear stress and atherosclerosis, lipid and atherosclerosis, and amino acid and pyruvate metabolism (Figure 2F, bottom left panel). These data suggest that in the plaque the differentially enriched EV-vesiculome (i.e., protein and miRNA) may function together, as well as independently, to modulate several pathways known to be involved in driving plaque biology. Given EVs are a vector of cell-cell communication between donor and recipient cells, we next attempted to determine the cellular origin of human plaque EVs to delineate the cellular contributors to plaque EV and further, how they might change with disease progression from marginal to plaque zones.

### Total EV-proteomics suggests diverse EV cell of origin and potential targets modulated in recipient cells

EVs carry cargo and membrane proteins that are reflective of their cell of origin.^56,57^ Cells secrete EVs with distinct biological cargo to relay specific messages to recipient cells. To date, little is known about the cellular communication occurring *in vivo* within the plaque niche via EVs. We used two complementary approaches to infer the EV cell of origin using our EV proteomics. We started by exploring our total EV proteomics (i.e. plaque and marginal regions combined) within the TISSUES database^39^ which integrates evidence on tissue expression from proteomic and transcriptomic screens. The predicted cellular EV origin included a range of expected cells including endothelial (neointima), several immune (B-lymphoblastoid, plasma cell, bone marrow cell, and foam cell), and smooth muscle cells (Figure 3A; Figure S8A,C). The analysis was then refined with EV-protein signature enrichment analysis among carotid artery cells specifically, by integrating our complete list of total detected EV-proteins with a publicly available scRNA-seq dataset containing paired plaque and marginal region samples (n=6 samples, Figure S9A-G).^20^ Data integration corroborated the EV cell of origin, identifying strong EV signatures from vascular smooth muscle cells, endothelial cells, dendritic cells, macrophages and lymphocytes (Figure 3B). Likewise, examination of EV-miRNA normalized counts in our plaque and marginal zones identified several miRNAs that are frequently ascribed to endothelial,^45,58–62^ smooth muscle,^63–65^ and lymphocyte cells,^47,66–70^ suggesting EV cell of origin signatures that align with those predicted by the EV-proteome (Figure S10A).

**Figure 3.**
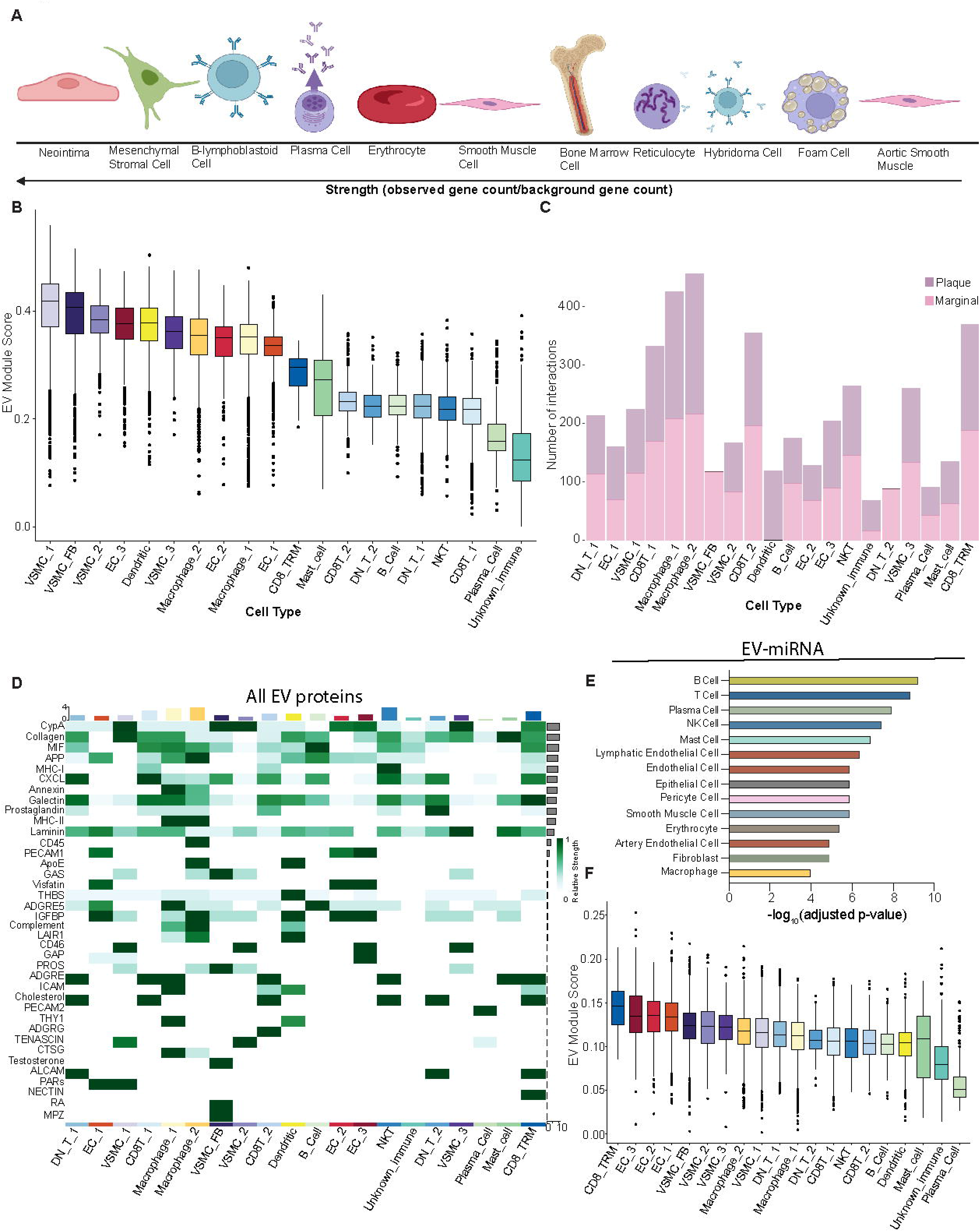
*In silico* analysis of human atherosclerotic tissue derive EV-cell-of-origin and global communication strategies in plaque and marginal zones. A. Pictorial representation of cell of origin using the TISSUES database with all EV-proteins. Top 11 cells by strength metric calculated as a ratio of observed and background gene count after inclusion of cell types only (tissue annotations filtered out; further details in Figure S8A). **B.** Box plot of global EV-protein signature enrichment among cells from atherosclerotic plaques, informed by integrating all detected EV-proteins (global EV signature) with single cell RNA sequencing (scRNA-seq) of carotid atherosclerotic plaques. **C.** Stacked bar plot visualizing total number of predicted cell-cell communication interactions in plaque (purple) and marginal (pink) zones with CellChat. **D.** Heatmap depicting EV-cell signaling strength employed by all EV-proteins to both plaque and marginal single cells. Strength of EV-derived signaling molecule and cell type shown on right and top bar graph, respectively, and represents communication probability. **E.** Cell type annotations derived using the Tabula Sapiens database (vascular tissue) by inputting all significant (adjusted p-value <0.05) predicted mRNA targets of all differentially expressed EV- miRNA in plaque and marginal zones combined (FDR<0.05). **F.** Box plot of predicted EV- miRNA-mRNA target signature enrichment among cells of the atherosclerotic plaque, informed by integrating differentially detected EV-miRNA - mRNA targets with scRNA-seq of carotid atherosclerotic plaques.

Beyond the suspected cell of origin, EV surface proteins and protein cargo deliver ligands to recipient cells within the atherosclerotic plaque environment – the biological consequences of this interaction will depend on the protein repertoire of the recipient cell. Using CellChat^71^, we predicted EV ligand communication (incoming signaling) with recipient cells that would be found in atherosclerosis using the same publicly available scRNA-seq dataset as above. Global EV-cell communication analysis was performed by specifically filtering for known receptor-ligand interactions where ligands were derived from EV proteomics and receptors from scRNA-seq.

When the total EV proteome was applied to scRNA-seq data annotated as plaque or marginal, it identified the predicted distribution of cellular EV source by each region (Figure 3C). Inferred cell communication pathways used total EV-associated ligands with total scRNA-seq (i.e. present in both plaque and marginal regions) (Figure 3D), as well as total EV-ligands separated by marginal and plaque regions (Figure S10C). The strongest global EV-cell interactions were seen among immune cells including macrophages and T-cells, with the enrichment of inflammatory signaling patterns (CypA, MIF, MHC-I, CXCL, prostaglandin, MHC-II). The predicted distribution of enriched global EV signaling pathways place ApoE, tenascin, cholesterol, and visfatin more likely within plaque zones (Figure S10B), while cord diagrams demonstrate predicted global EV ligand-receptor pairs with endothelial, macrophage, and vascular SMC populations (Figure S10D left, middle, and right panels, respectively).

Approaching EV-miRNA communication within tissue is more complex. Using the predicted messenger RNA (mRNA) targets of EV-miRNA, the genes that could be altered can be determined and correlated with single cell transcriptomics to understand potential EV-miRNA communication to recipient cells. We similarly used two approaches to annotate expected cellular targets of plaque EV-miRNAs - Tabula Sapiens database^72^ (vascular tissue) and the publicly available scRNA-seq carotid dataset as above (Figure 3E-F). All differentially expressed EV-miRNA had over 1956 (FDR<0.05) experimentally validated (miRTarBase)^73^ mRNA targets expressed in lymphocytes, endothelial cells, SMCs and macrophages (Figure 3E). Plaque EV-miRNA to mRNA gene-set enrichment analysis with the carotid scRNA-seq dataset corroborated these findings, wherein targets of plaque EV-miRNAs were enriched in lymphocytes, endothelial cells, vascular SMCs, and macrophages (Figure 3F).

### Plaque and marginal zone EV-proteomes predict different EV cells of origin, recipient cell repertoires, and expected communication with vascular cells

As the microenvironments of plaque and marginal zones are likely distinct (also implicated by initial exploration, Figure 3C), we recapitulated our approach to infer EV cellular origin and predicted targets in recipient cells by region using differentially enriched EV-proteins and differentially expressed EV-miRNA between plaque versus marginal zones. When delineated by region, plaque zones appear to have greater representation of macrophages as sources of EVs than marginal zones, where vascular SMCs predominated (Figure 4A-B). TISSUES database supported this finding, with our plaque and marginal zones highlighting diverse cellular EV sources that ranked differently by region (Figure 4C-D, S8B-C), predominantly reflecting cells that would be found in atherosclerosis despite the broader tissue sources included in this dataset.

**Figure 4.**
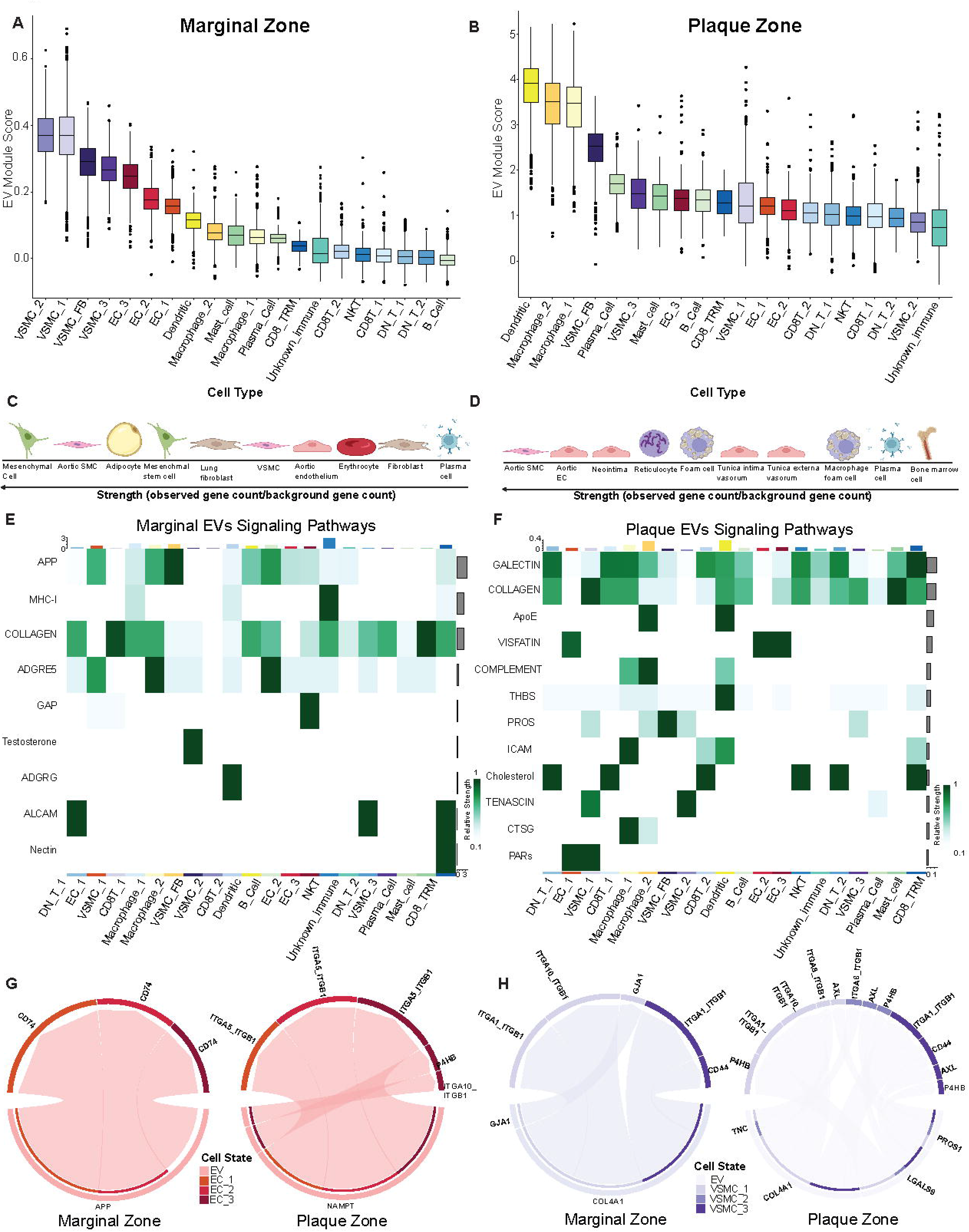
Site specific (plaque versus marginal zone) differences in EV cell of origin, recipient cell communication strategies and directed communication with vascular cells (ECs, SMCs, macrophages). A-B. Box plot of differential marginal **(A)** and plaque **(B)** EV-protein signature enrichment among cells of the atherosclerotic plaque, informed by integrating differentially enriched EV-proteins with scRNA-seq of carotid atherosclerotic plaques. **C-D.** Pictorial representation of cell of origin using the TISSUES database with differentially expressed EV-proteins in marginal **(C)** and plaque **(D)** zones. Top 10 cells by strength metric calculated as a ratio of observed and background gene count after filtering for tissues (further details in Figure S8B). **E-F.** Heatmap depicting differential marginal and plaque EV to cell signaling strength employed by differentially enriched EV proteins in marginal **(E)** and plaque **(F)** zones to all atherosclerotic single cells in plaque and marginal regions. Strength of EV-derived signaling molecule and cell type shown on right and top bar graph, respectively. **G-H.** Chord diagrams representing differentially expressed EV ligands to EC **(G)** and SMC **(H)** ligand-receptor pairs to single cells in marginal zones (left panels) and plaque zones (right panels). Arrows represent EV-ligand interactions. Individual cell populations annotated in legend.

We then pursued EV-cell communication signals within plaque and marginal regions separately, by filtering EV ligands as either plaque-enriched EV-proteins or marginal-enriched EV-proteins. Known ligand-receptor interactions were then explored using CellChat as above. In comparison to marginal zones (Figure 4E), plaque enriched EV-protein ligands were predicted to participate in more diverse and distinct cellular communication strategies when paired with total scRNA-seq (i.e. present in both plaque and marginal regions) (Figure 4F). For example, several predicted inflammatory ligand-receptor interactions emerged in the plaque zone with ApoE, complement, ICAM, cathepsin (CTSG), and cholesterol signatures. Similar increased communication diversity was predicted in plaque zones when analysis was restricted to single cells in marginal and plaque regions separately (Figure S11A-B). Chord communication diagrams were then created to explore region-specific cell-cell communication. Differentially enriched EV-protein ligands from our marginal and plaque zones were paired with publicly available single cell transcripts (potential receptor recipients) from either endothelial cells (Figure 4G), vascular SMCs (Figure 4H), or macrophages (Figure S11C) found in respective marginal regions (left panels) or plaque regions (right panels). For each cell type examined, plaque zones exhibited increased predicted ligand-receptor interactions relative to marginal zones. Notably, proposed plaque EV-cell communication included endothelial integrins (needed for cell motility), vascular SMC integrins and AXL (phagocytosis), and macrophage complement and cell adhesion (C3AR1, CD44).

Together, EVs are predicted to communicate with several atherosclerosis-relevant cell types, with regional (plaque and marginal) differences in communication patterns.

### Carotid plaque EVs are increased in number and contain EV-cargo (miRNA and protein) that are differentially enriched based on plaque vulnerability

Understanding the biological potential and EV-cell communication that takes place along the continuum of asymptomatic to symptomatic carotid disease will provide new insights into this complex clinical problem. We assessed differences in EV-cargo between asymptomatic (n=10 patients) and symptomatic (n=10-13 patients) groups by anatomical zones (marginal and plaque; Figure 5A). Blinded pathological grading performed by a staff pathologist (Dr. M. Seidman) of representative plaque samples (n=5) demonstrated fibroatheroma and calcification in all (AHA grade V-VI),^38,74^ with evidence of intraplaque hemorrhage reported in all symptomatic patients (Figure 5B, S12A). While both asymptomatic and symptomatic plaque zones contained more EVs per milligram of tissue compared to their matched marginal zones, only symptomatic patients achieved statistical significance (Figure 5C-D, S12B). Morphologically, carotid EVs were similar in symptomatic and asymptomatic groups (size and cryo-EM, Figure 5E). Direct comparisons by symptom status within plaque and marginal zones did not yield significant differences in EV quantification highlighting the importance of matched marginal zone controls (Figure S12C-D).

**Figure 5.**
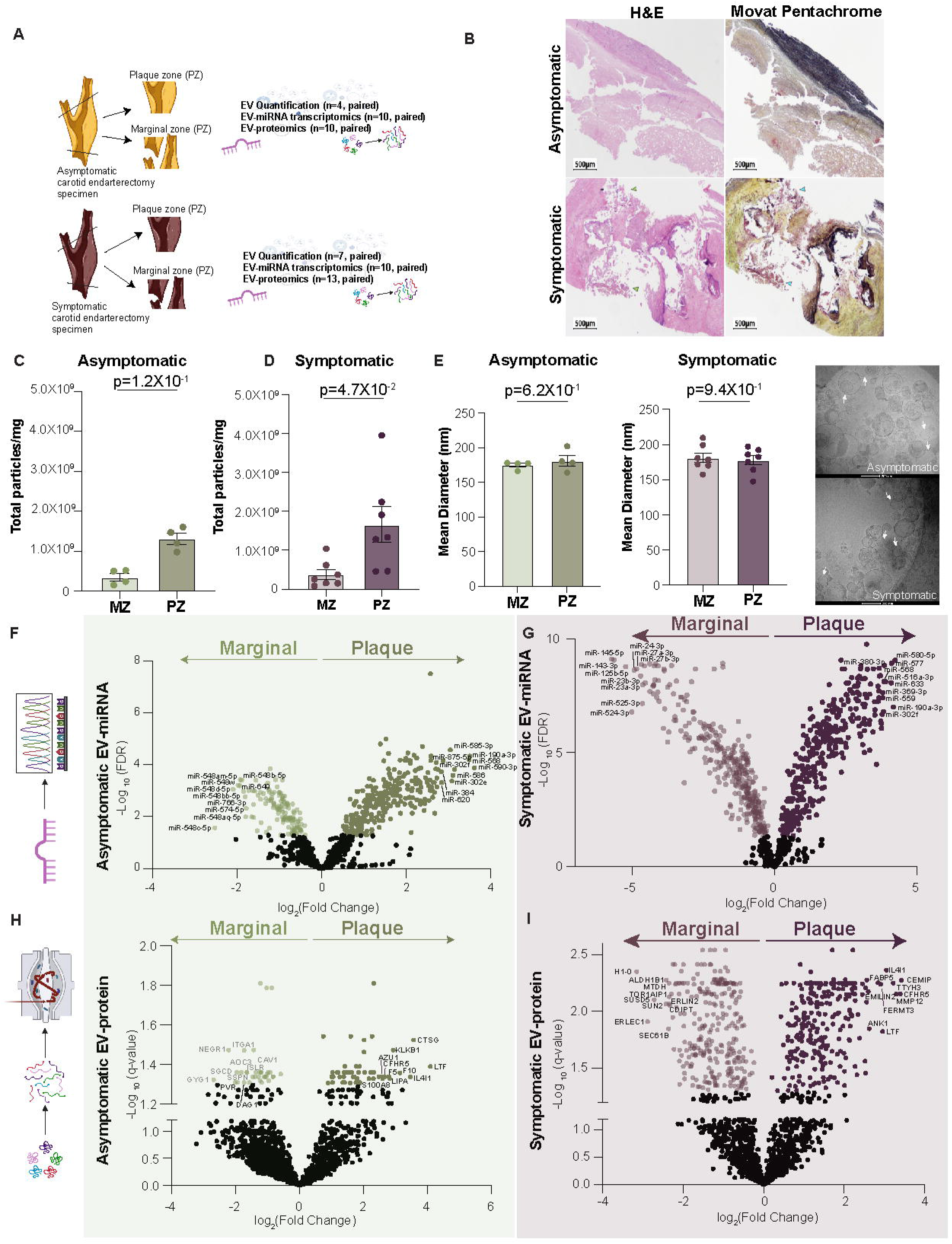
EV concentration and greater differential EV cargo (miRNA and protein) discriminates between symptomatic and asymptomatic atherosclerotic plaques. A. Schematic of experimental design. Carotid endarterectomy specimens from symptomatic and asymptomatic patients were dissected into plaque and marginal zones, followed by EV- enrichement, quantification, and miRNA transcriptomics and proteomics. **B.** Hematoxylin and eosin (H&E) and Movat pentachrome stain of asymptomatic (top panel) and symptomatic (bottom panel) of carotid plaque specimens (Patient 1 and 2). Large, calcified plaque with calcified/necrotic center, but with no significant intraplaque hemorrhage (top panel). Large, calcified plaque with calcified/necrotic center with intraplaque hemorrhage (bottom panel). H&E staining with intraplaque hemorrhage indicated by green arrowheads (bottom left panel) and Movat pentachrome staining with intraplaque hemorrhage indicated by cyan arrowheads (bottom right panel). Scale bar (500 µm) as shown. **C-D.** Nanoparticle tracking analysis (NTA) of nanoparticle concentration in asymptomatic **(C)** and symptomatic **(D)** atherosclerotic plaque versus marginal zones. **E.** Assessment of EV size and morphology. NTA of mean diameter of nanoparticles enriched from asymptomatic (left panel) and symptomatic (middle panel) carotid plaques. Cryogenic electron microscopy (cryo-EM) of EVs from asymptomatic (right panel, top) and symptomatic (right panel, bottom) carotid plaques (*n*=1-2 per group). Arrows indicate select EV structures as bilayered nanoparticles with dense cores. Scale bar = 200nm. **F-G.** Volcano plot of carotid tissue resident EV-miRNA transcriptome in asymptomatic **(F)** and symptomatic **(G)** with positive and negative fold change (FC) representing miRNAs enriched in plaque versus marginal zones (FDR step up<0.05). **H-I.** Volcano plot of carotid tissue resident EV-proteome in asymptomatic **(H)** and symptomatic **(I)** with positive and negative fold change (FC) representing EV-proteins enriched in plaque versus marginal zones (q-value<0.05). Bar graphs show mean ± SEM. Statistical significance assessed by non-parametric paired Wilcoxon test **(C-E)**.

When analysis of EV-cargos was performed, there were consistently larger numbers of differentially enriched EV-miRNA and EV-proteins in the symptomatic group in both marginal zones (delineated by negative fold change) and plaque zones (delineated by positive fold change) (Figure 5F-I), suggesting increased biological complexity in clinically vulnerable plaques and their corresponding marginal regions for symptomatic patients.

We next assessed specific differences in EV-cargo between asymptomatic plaques versus paired marginal zones and separately, EV-cargo differences between symptomatic plaques versus paired marginal zones (Figure 6). There were 306 unique differentially expressed EV-miRNAs in symptomatic tissue compared to only 33 unique EV-miRNA in asymptomatic tissue (Figure 6A). Overlap was seen with 364 differentially expressed EV-miRNAs in the two groups. Pathway analysis of unique EV-miRNA symptomatic predicted targets revealed more GO and KEGG pathways (Figure 6B). Unique signatures in asymptomatic tissues included steroid biosynthesis, ABC transporters, and the AMPK signaling pathway (Figure 6C). Pathways derived using unique symptomatic EV-miRNAs were numerous and included lipid and atherosclerosis, inflammatory signalling (HIF-1, TGFβ, TNF) and NOTCH signaling (endothelial angiogenesis) (Figure 6D). Similarly, 435 unique differentially enriched EV-proteins were seen in symptomatic tissue, compared to 6 in asymptomatic (Figure 6E) and functioned with a high degree of biological interconnectibility (Figure S12E). Similar to the EV-miRNA analysis, there were an increased number of GO and KEGG pathways modulated by the symptomatic plaque EV-protein contents (Figure 6F). KEGG pathway analysis of the unique EV-proteins from asymptomatic tissue identified 6 KEGG pathways functioning in amino acid and nutrient metabolism, antigen presentation, renin secretion, and complement and coagulation with only a single EV-protein (annotated in figure) contributing to each pathway protein (Figure 6G). The unique EV-proteins from symptomatic tissue identified several biological pathways with the top differentially expressed pathways including complement and coagulation, cell motility (regulation of actin cytoskeleton, focal adhesion, ECM-receptor interaction, leukocyte transendothelial migration), and cholesterol metabolism (Figure 6H, left panel). Several proteins were enriched in the symptomatic plaque EVs that demonstrated protein-protein interactions, forming three units (complement and coagulation cascades, cholesterol metabolism, and regulation of actin cytoskeleton) predicted to modulate these pathways (Figure 6H, right panel). Together, these vesiculomic data highlight the potential for EVs to govern several well-known aspects of plaque biology, with increasing complexity as the disease continuum progresses.

**Figure 6.**
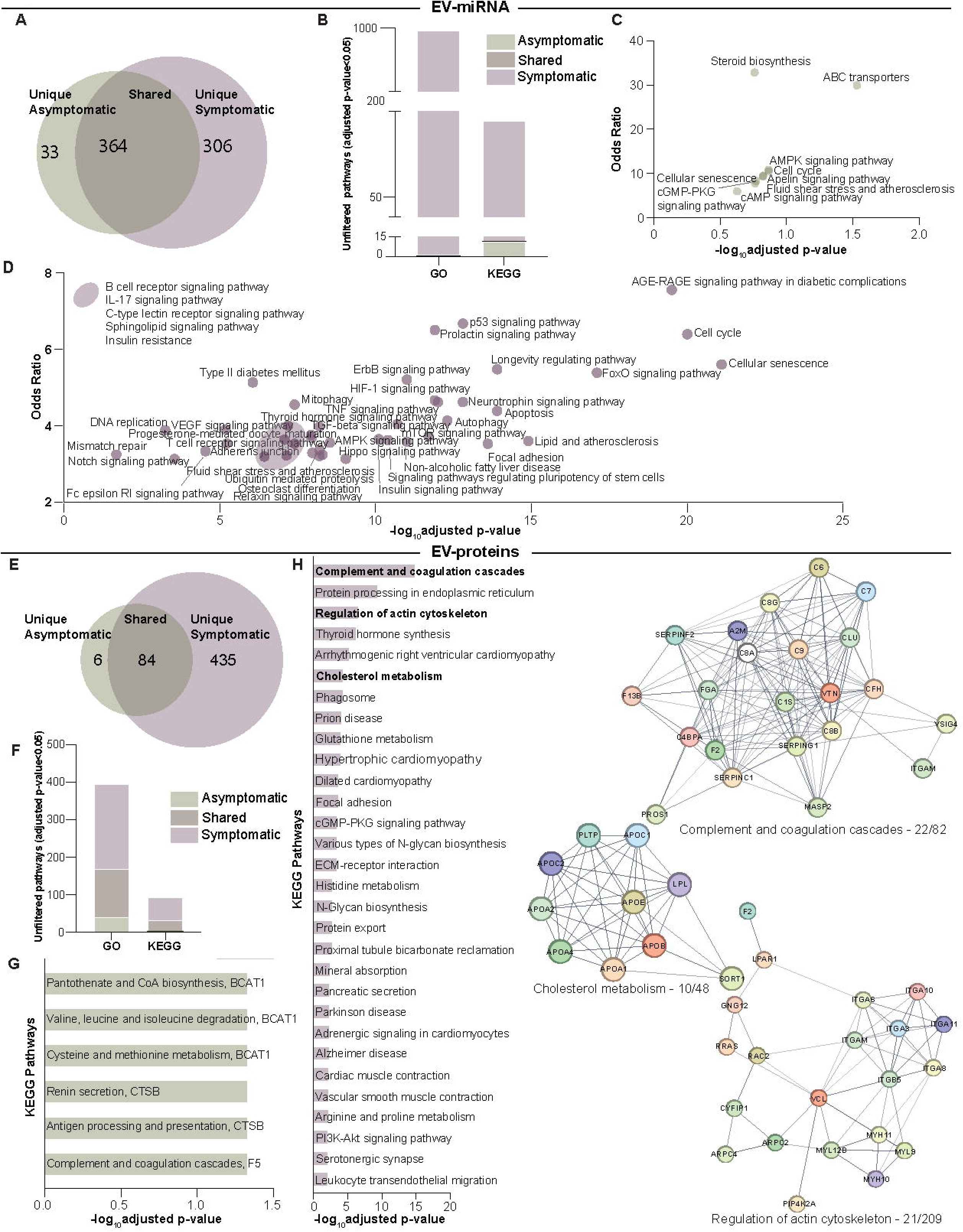
Differential plaque EV-secretome highlights key pathways involved in plaque vulnerability. A. Venn diagram depicting shared and unique plaque derived EV-miRNA in asymptomatic (green) and symptomatic (purple) conditions (FDR step-up<0.05, filtered for miRNAs<1000). **B.** Total numbers of significant Gene Ontology (GO) and Kyoto Encyclopedia of Genes and Genomes (KEGG) pathways (adjusted p-value<0.05) modulated by the predicted mRNA targets (miRTarBase via MIENTURNET, threshold of minimum interactions 1; FDR<0.05) of differentially expressed plaque EV-miRNAs in asymptomatic (green), symptomatic (purple), and shared (brown) conditions. **C.** Dot plot of all KEGG pathways derived from unique asymptomatic plaque-derived predicted EV-miRNA mRNA targets (p-value<0.05). Labelled are significant (adjusted p-value<0.05) pathways graphed by odds ratio. **D.** Dot plot of KEGG pathway analysis from unique symptomatic plaque-derived predicted EV-miRNA mRNA targets. Graphed are top 42 significant (adjusted p-value<0.05) pathways by odds ratio (total 120 pathways with adjusted p-value<0.05). **E.** Venn diagram depicting shared and unique plaque derived EV-proteins in asymptomatic (green) and symptomatic (purple) conditions (q- value<0.05). **F.** Total numbers of significant GO and KEGG pathways (adjusted p-value<0.05) modulated by differentially expressed plaque EV-proteins (q-value<0.05) in asymptomatic (green), symptomatic (purple), and shared (brown) conditions. **G.** Bar graph of all significant KEGG pathways (adjusted p-value<0.05) from unique asymptomatic plaque-derived EV- proteins (adjusted p-value<0.05). **H.** Bar graph of all significant KEGG pathways (adjusted p- value<0.05) from unique symptomatic plaque-derived EV-proteins (adjusted p-value<0.05). STRING (db version 12.0) used to generate protein-protein interaction maps (right panel) of pathways of interest. Cancer-, and infection- associated pathways were excluded from analysis.

### Predicted EV-based cell communication in the symptomatic plaque reveals the emergence of an endothelial cell signature with functional effects on angiogenesis

The differences in EV cargo between asymptomatic and symptomatic groups led us to query differences in predicted cellular communication based on symptom status. EV cellular identity was inferred by integrating our enriched EV-protein signatures from either marginal or plaque zones according to symptom status with carotid marginal and plaque scRNA-seq data (Figure 7A-D). Greater endothelial cell (EC) enrichment was predicted in symptomatic plaque signatures relative to asymptomatic plaque signatures (purple bars highlight each EC subpopulation, with increasing prevalence noted as a shift to the left; Figure 7B,D), possibly suggesting a heightened role of endothelial cell-derived EVs in governing EV-cell communication with atherosclerotic plaque progression. No notable differences in cell type enrichment was observed for marginal signatures (Figure 7A,C). Potential recipient cell repertoires available in asymptomatic and symptomatic plaques was also explored through EV-miRNA-mRNA predicted targets (Figure S13).

**Figure 7.**
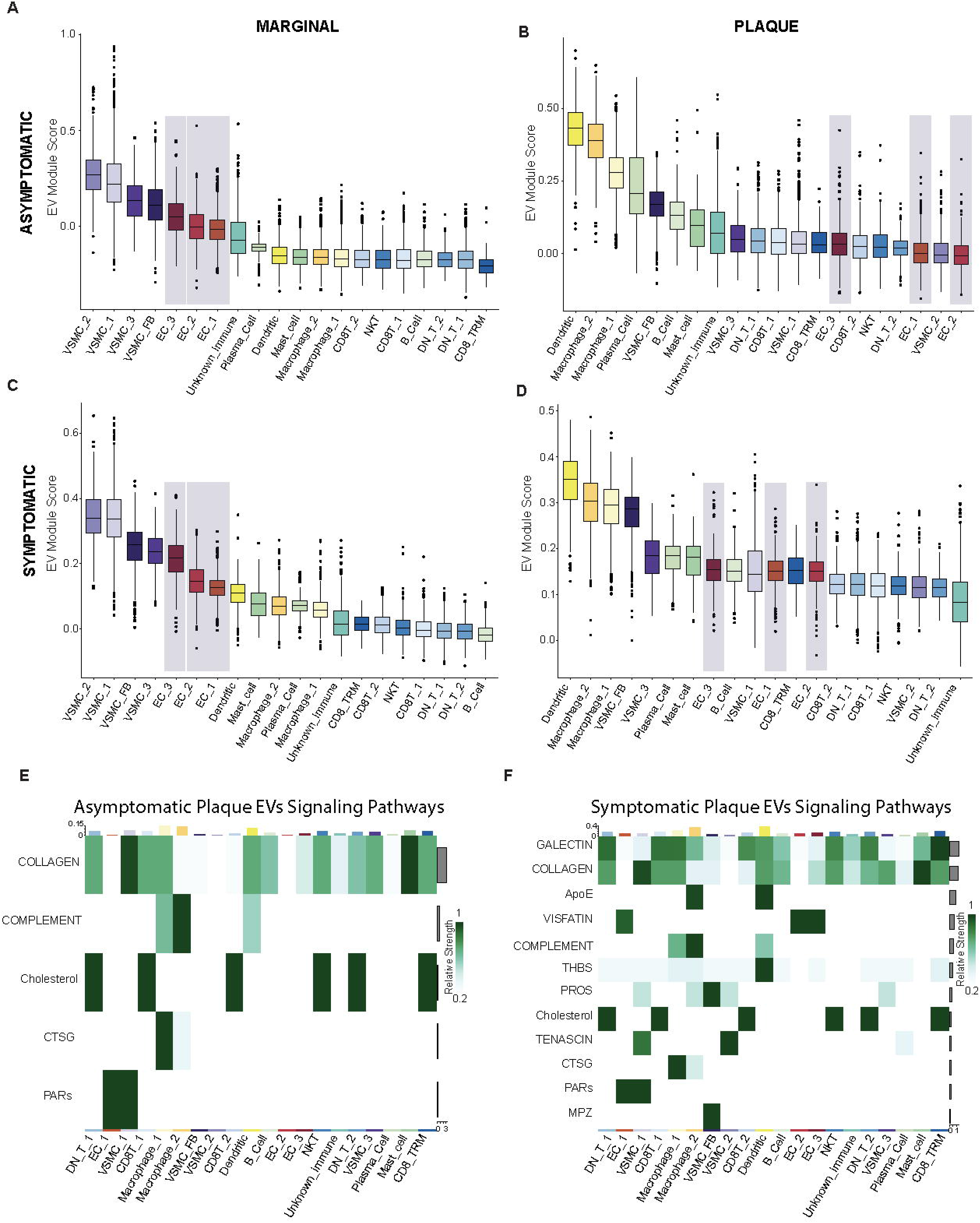
Endothelial cells (ECs) are cellular players modulating cell-cell communication through EVs in vulnerable plaques. A-D. Box plots of symptom and regional EV-protein signature enrichment among cells of the atherosclerotic plaque, informed by integrating differentially expressed EV-proteins in asymptomatic and symptomatic, marginal versus plaque zones with scRNA-seq of carotid atherosclerotic plaques. Asymptomatic marginal **(A)** and plaque **(B)** zone signature enrichment among cells of the atherosclerotic plaque. Symptomatic marginal **(C)** and plaque **(D)** zone signature enrichment among cells of the atherosclerotic plaque. **E-F.** Heatmap depicting differential asymptomatic plaque- **(E)** and symptomatic plaque-**(F)** EV-cell signaling strength employed by differentially expressed EV-proteins in asymptomatic and symptomatic cohorts with plaque cells. Strength of EV-derived signaling molecule and cell type shown on right and top bar graph, respectively.

We then explored differences in asymptomatic and symptomatic plaque EV-cell communication with CellChat, by similarly examining EV-proteins as the ligands in known ligand-receptor interactions, where receptors were predicted from scRNA-seq in marginal and plaque zones (Figure 7E-F). Although asymptomatic plaques shared all EV-cell signaling pathways with the symptomatic group (Figure 7E-F), symptomatic plaque EV-protein ligands also exhibited unique signaling involving ApoE, galectin, and visfatin. Specifically, EV/visfatin to endothelial cell-based communication was unique to symptomatic plaques – notable given angiogenesis can be modulated through visfatin-based secretion of vascular endothelial growth factor (VEGF) and fibroblast growth factor 2 (FGF-2),^75,76^ monocyte recruitment through monocyte chemoattractant protein 1 (MCP-1),^77^ and extracellular matrix remodelling via matrix metalloproteinase expression.^78,79^ We then explored specific ligand-receptor pairs between asymptomatic and symptomatic plaque EVs and recipient cell repertoires that might be available in ECs (Figure 8A, S14A), SMCs (Figure S14B-D), and macrophages (Figure S14E-G). Cell-cell communication strategies predicted in the asymptomatic plaques were mostly recapitulated in the symptomatic group, with symptomatic plaques exhibiting more complex EV-recipient cell signaling (Figure 8A, S14D,G). As an approach to externally validate some of our cell communication predictions, we examined the EV marker CD63 as well as CD63-normalized visfatin and CD63-normalized VEGFA in a cohort of 18 carotid endarterectomy specimens (Biobank of Karolinska Endarterectomy, BiKE) where proteomics had been completed on plaque zones and marginal zones (*n*=9 symptomatic, *n*=9 asymptomatic).^37^ While these were bulk tissue proteomics, more of the common EV marker CD63 protein was detected in symptomatic plaques and increased visfatin (NAMPT) and VEGFA was detected in symptomatic carotid tissue when normalized to CD63 protein by zone (Figure S15A-C).

**Figure 8.**
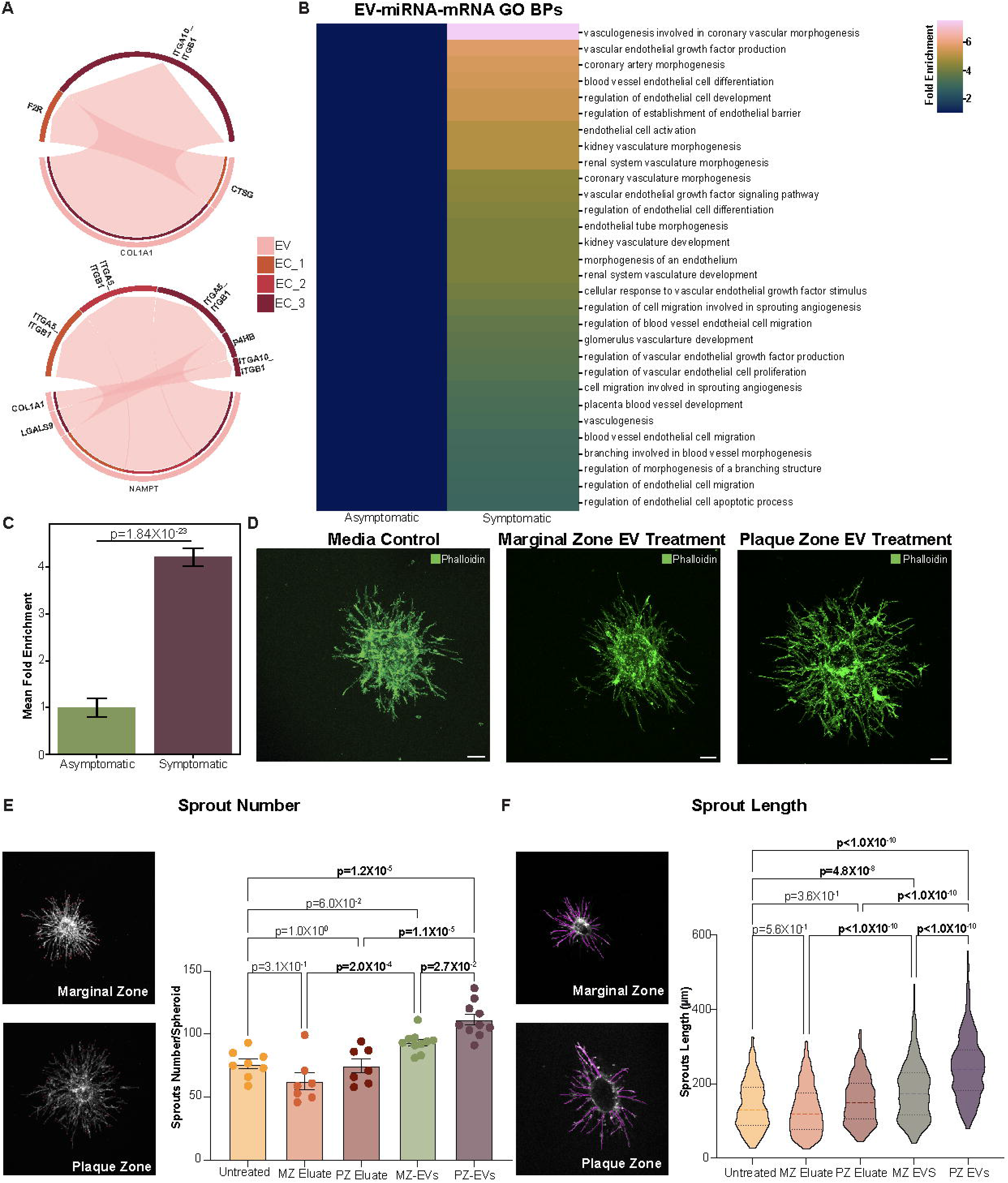
Overrepresentation analysis identifies angiogenesis and neovascularization as functional effects of vulnerable (symptomatic) plaque EVs with support from a functional EC sprouting assay. A. Chord diagrams representing ligand-receptor pairs between differentially enriched EV-proteins (ligands) to ECs (receptor) in asymptomatic (top panel) and symptomatic (bottom panel) plaques. Red arrows represent EV-ligand to receptor interactions. Individual cell populations annotated in legend. **B.** Filtered inquiry of EC, angiogenesis, and neovascularization specific pathways from Gene Ontology (GO) overrepresentation analysis of differentially expressed predicted EV-miRNA-mRNA targets in asymptomatic and symptomatic groups. Fold enrichment represented in legend. **C.** Pairwise comparison of mean fold enrichment of EC-specific pathways in symptomatic versus asymptomatic plaques. **D-F.** Functional effect of symptomatic marginal zone (MZ) and plaque zone (PZ) EVs on endothelial cell angiogenesis. Representative confocal microscopy images of HUVEC-spheroid angiogenesis after addition of EVs (2-4X10^10^ EVs; 24 h), or assay control **(D)**. Scale bar 100 µm. Representative grayscale images with annotations for sprout number **(E, left panel)** and quantification of sprout number per spheroid (*n*=7-10 from 3 independent experiments; **E, right panel**). Representative grayscale images with tracings representing individual sprout lengths **(F, left panel)** and quantification (*n*=188-331 sprouts from 7–10 spheroids from 3 independent experiments). Bar graphs shows the mean±SEM. Violin plots show median ± interquartile range. Statistical significance assessed by 1-way ANOVA with Sidak multiple comparisons test (adjusted p-values shown; **E,F)**. P-values meeting significance (<0.05) shown in bolded text.

Given the distinct EV-based communication strategies predicted in symptomatic versus asymptomatic tissues, we then assessed the potential functional consequences through analysis by GO and KEGG pathways for EV-miRNA, EV-proteins, and EV-vesiculome

(integrated analysis) (Figure S16A,B). We found differences in pathways predicted to be modulated by EVs in symptomatic compared to asymptomatic plaques. Key EV-miRNA based pathways enriched in symptomatic plaque included inflammatory signaling (TCR signaling, NFkB, HIF-1, TNF, TGFβ), apoptosis, cell senescence, and cell migration/motility (adherens junctions, VEGF) (Figure S16A,B left panels). Enriched pathways in symptomatic plaques predicted to be modulated by EV-proteins included complement and coagulation (includes fibrinolysis, iron metabolism), cholesterol metabolism (includes lipid transport, response to low density lipoprotein particle stimulus), and amino acid metabolism (Figure S16 A,B middle panels). Network analysis integrating EV-miRNA and EV-protein content identified the importance of metabolic pathways (amino acid biosynthesis, RNA synthesis and turnover, steroid biosynthesis, cholesterol), complement and coagulation, and inflammatory pathways (antigen processing and presentation, macrophage proliferation, neutrophil degranulation) as shared targets of EV-contents in symptomatic plaques (Figure S16 A-B right panels). Likewise, overrepresentation analysis of differentially enriched EV-miRNA and EV-protein predicted differential representation of GO and KEGG pathways that based on symptom status (Figure S17A-D). Upon closer examination there was a clear endothelial and angiogenic signature in EV-miRNA-mRNA overrepresentation analysis in symptomatic versus asymptomatic EVs with overrepresented predicted pathways in angiogenesis, blood vessel development, sprouting angiogenesis and VEGF signaling (Figure 8B, S18A-B), while EV-protein and EV-vesiculomics overrepresentation analysis were more varied (Figure S18C-F, S19A-D).

As significantly more endothelial angiogenesis-related pathways were identified in symptomatic plaque zones versus their respective marginal zones (Figure 8C, p-value = 1.84X10^-23^), we tested the functional effects of symptomatic plaque EVs (n=3) using an endothelial spheroid-based sprouting assay (Figure 8D-F, S19E-J). Endothelial sprouts were significantly increased in number (110.2±4.34 versus. 92.3±2.59; p=2.7X10^-2^) (Figure 8E) as well as total sprout length (244.9±5.06 versus 179.7±4.27 μm; p<1.0X10^-10^) (Figure 8F) and average sprout length (Figure S19G) after treatment with symptomatic plaque EVs compared to matched marginal zone EVs. Controls included eluates (negative control, Figure 8E-F, S19F-G) and VEGF (positive control, Figure S19F,H-J). Taken together, these results highlight the biologically relevant EV-protein and EV-miRNA cargos that exist within the plaque and marginal zones of carotid tissue along the continuum of asymptomatic to symptomatic disease.

## Discussion

It is currently unknown what renders a carotid atherosclerotic plaque vulnerable to rupture and leads to stroke. Cell-cell communication – including through extracellular vesicles (EVs)^31^ – may play a role. Our study evaluates carotid tissue EV-transcriptomics and EV-proteomics to understand changes in EV-cargo in diseased states (plaque versus marginal zones) along the clinical continuum (asymptomatic to symptomatic) to predict EV source, potential recipient cell responses, and a relevant biological functional effect based upon these predictions. Increased numbers of EVs containing distinct cargos were discovered in carotid plaque versus matched adjacent marginal zones, implicating their role in governing plaque biology. Given EV-based cell-cell communication necessitates an EV donor and an EV recipient cell, we integrated publicly available datasets (TISSUES^39^, Tabula Sapiens^72^ and carotid plaque scRNA-seq datasets^20^) with our region-specific carotid EV-vesiculomics to gain insight into the cellular drivers of EV-based communication within the plaque. As plaque vulnerability increased from asymptomatic to symptomatic disease states, there was a greater degree of EV communication suggested in the symptomatic plaques. Our *in silico* analysis suggested an increased abundance of endothelial cell-derived EVs with plaque progression and predicted a greater number of unique EV-endothelial cell ligand-receptor pairs in symptomatic plaques known to function in angiogenesis, with downstream overrepresentation analysis corroborating increased endothelial-based pathways. Intriguingly, we discovered an EV-derived endothelial/neovascularization ‘signature’ in symptomatic carotid plaques and then showed for the first time, that human symptomatic carotid plaque EVs induce angiogenesis in a model of endothelial sprouting. Taken together, these exciting data illustrate a complex communication network, with renewed focus on the role of EVs and the endothelium.

EVs are mediators of cardiovascular disease states and have previously been shown to be increased in patient samples.^31,80–84^ Here, we optimized and validated a protocol to enrich tissue- entrapped atherosclerotic plaque-derived EVs using size exclusion chromatography and validated our EVs in accordance with the MISEV2023 criteria,^26^ in alignment with our previous EV isolation approaches using similar methodology.^31^ We found increased EVs per milligram of tissue from plaque regions in comparison to their matched adjacent marginal regions. Recent work has delineated an increase in EV signature based on pathway analysis of differential protein expression from diseased areas (carotid plaque and aortic valve).^31^ EVs mediate their effects via delivering their cargo such as miRNAs and proteins to recipient cells.^32–34^ In carotid tissue, this has been difficult to test empirically. To date, there is evidence that carotid plaque-derived microparticles are thrombogenic^27^ and that candidate EV-cargo (from multiomics analysis of carotid tissue) can increase calcification in cultured human carotid SMCs.^31^ Given the challenges working with EVs^31,85^, there has been no study to date using human carotid plaque derived EVs in functional assays to determine the biological consequences of altered

EV-cargo in diseased states. By examining the EV-vesiculome within our carotid plaque versus marginal zones, plaque EV-contents were predicted to be involved in several pro-atherogenic pathways such as complement and coagulation, fluid shear stress, cellular senescence, cell proliferation and differentiation, and immune pathways. As EVs carry proteins that are indicative of their cell of origin,^56,86^ we harnessed an opportunity to examine our unique dataset within publicly available single cell RNA -sequencing datasets to explore potential EV-based cell-cell communication in the carotid atherosclerotic plaque milieu.

To assess EV-based communication in the plaque microenvironment, we layered our EV- vesiculomics with the only published carotid plaque scRNA-seq dataset that contained site-specific data (plaque zone and marginal zone) to reveal potential EV donor cell identities and communication partners in our two zones of interest.^20,56,86^ Site-specific layering is important in atherosclerotic disease given the cellular plasticity seen in plaque progression.^12,87,88^ Leveraging tissue-specific scRNA-seq data to inform the probable cell of origin of EVs has recently gained acceptance as a starting point in complex human tissue.^57^ The differences in EV cell of origin identity with vascular SMCs predominating in marginal zones and immune cells in plaques zones is in keeping with recent reports of different identities and quantities of various cell populations found in plaques versus the adjacent region.^20^ The role of SMCs in regulating atherosclerotic plaque progression through phenotypic switching in atherogenic environments^11,87^ is well characterized and our data now suggest that SMCs could also be important sources of EVs in marginal zones, beyond their roles within advanced plaques.^12,20,88,89^ The role of immune cells in governing advanced plaque biology is similarly acknowledged, and likely explains the increased EV contribution predicted from these cells in plaque regions.^90,91^ Intriguingly, the ligand-receptor communication strategies (using EV-protein cargo as ligands and recipient cell repertoires from scRNA-seq) that were predicted in the plaque versus marginal zones suggested increasing complexity of the biological milieu along the disease continuum.^20,31^ Our analysis implicates EVs as integral mediators of atherosclerotic plaque progression and EV-based communication as a potential target to modulate plaque biology.

Assessment and treatment of vulnerable carotid atherosclerotic plaques represents a continuing clinical challenge with opportunities to improve.^92^ With symptomatic plaques representing culprit lesions with advanced vulnerable atherosclerotic disease where participation from resident and circulating cells that are phenotypically dynamic contribute to the EV-pool, disease progression lends itself to multifaceted communication patterns.^93^ We defined a predicted role for endothelial cells at both facets (donor and recipient) of the EV cell-cell communication paradigm in symptomatic plaques, where endothelial cells featured prominently as both potential sources of EVs and responders to EVs – specifically with communication signaling known to function in neovascularization and angiogenesis. Neovascularization in atherosclerotic plaque occurs as a result of hypoxemia and leads to the growth of immature and leaky vessels causing intra-plaque hemorrhage, a well known contributor to plaque instability and rupture.^37,94–96^ While not unique to symptomatic plaques, it does predominate in this group of patients.^97^ With this insight, we corroborated our predictions functionally and showed for the first time that symptomatic human carotid plaque zone EVs induce angiogenesis using an optimized low-input human endothelial spheroid sprouting assay.

The importance of endothelial EV-based communication in driving atherosclerotic plaque instability represents an opportunity to understand the disease process from the less-studied endothelial-centric lens. Endothelial cell-EVs are intimately involved in autocrine and paracrine communication with cells of the atherosclerotic plaque where they orchestrate upregulation of adhesion molecules, and activation, proliferation, and migration of vascular cells.^7,98–104^ In the atherosclerotic milieu, endothelial cells are exposed to disturbed flow and inflammatory mediators, and can undergo endothelial-to-mesenchymal transition wherein they lose endothelial identify and adopt mesenchymal properties.^9,10,105^ In the scRNA-seq dataset from Alsaigh *et al*, there were three endothelial cell populations contributing to plaque-derived EVs as seen in the original scRNA-sequencing dataset that represented quiescent, activated, and transdifferentiated endothelial cells.^20^ Given the quiescent endothelium has been shown to participate in atheroprotective EV-messaging^104^, it would be beneficial to resolve the differences in communication between these cell types and understand how to shift the messaging to a quiescent phenotype. The important role of endothelial cell-based EV communication in carotid atherosclerotic plaque combined with the accessibility of the endothelial layer, emphasizes the potential of endothelial-EV targeted therapeutics in atherosclerotic vascular disease.

Our study explored EV-based cell-cell communication in human atherosclerotic disease and how this might dictate plaque vulnerability in part through the functional role of EVs in endothelial angiogenesis. There are limitations, however. While we completed tissue EV- transcriptomics and EV-proteomics, it was not possible to generate matched scRNA-sequencing that would have allowed us to annotate the cells and communication occurring within the same carotid tissue.^57^ The external dataset selected was the only publicly available scRNAseq on plaque and marginal zones, however it did not contain symptomatic patients. Given the cell identity changes seen with plaque progression, cell annotations and ligand-receptor interactions may not have captured the diversity of the symptomatic plaque cell populations specifically. *In silico* analysis only enables predictions for delineating EV cells of origin and potential recipient cell responses, which could be a limitation across all samples – regardless of patient symptom status. Lastly, EVs are one of many extracellular signals in the plaque microenvironment. While it is encouraging that increased CD63 (common EV marker) was similarly detected in symptomatic plaque zones in the BiKE dataset, and was significantly increased in the most severe Cluster 3 carotid plaques described by Mokry et al,^106^ the role of lipoproteins, non-EV inflammatory or angiogenic processes, and metabolites cannot be excluded.^107^ Our work does not preclude the role of other signalling molecules, nor can it account for the dynamic, spatial, and other physical elements taking place. As technologies progress, it may become possible to validate our findings at a single EV-level.

## Conclusions

This work was pursued with the challenging goal of establishing an important conversation around EVs in carotid plaque biology. Crucially, vulnerable symptomatic carotid disease was characterized by an increased concentration of EVs in plaque zones, with increased contribution from endothelial cell sources. Furthermore, EVs harbored a signature for specific EV-based communication with endothelial cells, with predicted impact on angiogenic pathways. These predictions were corroborated functionally by demonstrating symptomatic carotid plaque zone EVs increase endothelial sprouting angiogenesis – providing a mechanism for neovascularization and plaque rupture potential. These provocative results underscore the importance of the endothelium in governing vulnerable biology and highlight unique opportunities to consider the role of EVs as a therapeutic target in plaque stabilizing therapies.

## Acknowledgements

The authors thank the patients, surgical teams, and stroke neurology colleagues at University Health Network and Sunnybrook Health Sciences. For assistance with pathology: Ms. Melanie Peralta (UHN). Electron microscopy imaging of EVs was performed in the Harvard Medical School Electron Microscopy Facility.

## Disclosures

Natalie Galant is co-founder and CEO of Paradox Immunotherapeutics, but the content of this manuscript is unrelated to the work of this company.

**Supplemental Material** Detailed Methods Figures S1-S19 Supplemental Legends References 108-136

## Non-Standard Abbreviations

cryo-EM: cryogenic electron microscopy EC endothelial cell
EV: extracellular vesicle
GO: Gene Ontology
KEGG: Kyoto Encyclopedia of Genes and Genomes
NTA: nanoparticle tracking analysis miRNA microRNA MISEV Minimal information for studies of extracellular vesicles
MZ: marginal zone
PZ: plaque zone
scRNA-: seq single cell RNA-sequencing
SEC: size exclusion chromotography
SMC: smooth muscle cell
TEM: transmission electron microscopy

## References

1. Libby P. The changing landscape of atherosclerosis. Nature. 2021;592(7855):524-533.

2. Libby P, Buring JE, Badimon L, et al. Atherosclerosis. Nature Reviews Disease Primers. 2019;5(1):56.

3. Shahjouei S, Sadighi A, Chaudhary D, et al. A 5-decade analysis of incidence trends of ischemic stroke aRer transient ischemic attack: a systematic review and meta-analysis. JAMA neurology. 2021;78(1):77–87.

4. Howe KL, Cybulsky M, Fish JE. The Endothelium as a Hub for Cellular Communication in Atherogenesis: Is There Directionality to the Message? Front Cardiovasc Med. 2022;9:888390.

5. Yu L, Xu L, Chu H, et al. Macrophage-to-endothelial cell crosstalk by the cholesterol metabolite 27HC promotes atherosclerosis in male mice. Nat Commun. 2023;14(1):4101.

6. Yurdagul A, Jr. Crosstalk Between Macrophages and Vascular Smooth Muscle Cells in Atherosclerotic Plaque Stability. Arterioscler Thromb Vasc Biol. 2022;42(4):372–380.

7. Hergenreider E, Heydt S, Tréguer K, et al. Atheroprotective communication between endothelial cells and smooth muscle cells through miRNAs. Nature cell biology. 2012;14(3):249–256.

8. Raju S, Botts SR, Blaser MC, et al. Directional Endothelial Communication by Polarized Extracellular Vesicle Release. Circulation Research. 2024;134(3):269–289.

9. Chen P-Y, Qin L, Baeyens N, et al. Endothelial-to-mesenchymal transition drives atherosclerosis progression. The Journal of clinical investigation. 2015;125(12):4514–4528.

10. Evrard SM, Lecce L, Michelis KC, et al. Endothelial to mesenchymal transition is common in atherosclerotic lesions and is associated with plaque instability. Nature communications. 2016;7(1):11853.

11. Gomez D, Owens GK. Smooth muscle cell phenotypic switching in atherosclerosis. Cardiovascular research. 2012;95(2):156–164.

12. Alencar GF, Owsiany KM, Karnewar S, et al. Stem cell pluripotency genes Klf4 and Oct4 regulate complex SMC phenotypic changes critical in late-stage atherosclerotic lesion pathogenesis. Circulation. 2020;142(21):2045–2059.

13. Brott TG, Hobson RW, Howard G, et al. Stenting versus endarterectomy for treatment of carotid- artery stenosis. New England Journal of Medicine. 2010;363(1):11–23.

14. Ferguson GG, Eliasziw M, Barr HW, et al. The North American symptomatic carotid endarterectomy trial: surgical results in 1415 patients. Stroke. 1999;30(9):1751–1758.

15. Group ECSTC. Randomised trial of endarterectomy for recently symptomatic carotid stenosis: final results of the MRC European Carotid Surgery Trial (ECST). The Lancet. 1998;351(9113):1379–1387.

16. Walker MD, Marler JR, Goldstein M, et al. Endarterectomy for asymptomatic carotid artery stenosis. Jama. 1995;273(18):1421–1428.

17. Hackam DG. Optimal medical management of asymptomatic carotid stenosis. Stroke. 2021;52(6):2191–2198.

18. Committee W. European Society for Vascular Surgery (ESVS) 2023 Clinical Practice Guidelines on Radiation Safety. European Journal of Vascular and Endovascular Surgery. 2022.

19. Goessens BM, Visseren FL, Kappelle LJ, Algra A, van der Graaf Y. Asymptomatic carotid artery stenosis and the risk of new vascular events in patients with manifest arterial disease: the SMART study. Stroke. 2007;38(5):1470–1475.

20. Alsaigh T, Evans D, Frankel D, Torkamani A. Decoding the transcriptome of calcified atherosclerotic plaque at single-cell resolution. Communications biology. 2022;5(1):1084.

21. Theofilatos K, Stojkovic S, Hasman M, et al. Proteomic atlas of atherosclerosis: the contribution of proteoglycans to sex differences, plaque phenotypes, and outcomes. Circulation Research. 2023;133(7):542–558.

22. Tomas L, Edsfeldt A, Mollet IG, et al. Altered metabolism distinguishes high-risk from stable carotid atherosclerotic plaques. European heart journal. 2018;39(24):2301–2310.

23. Vaisar T, Hu JH, Airhart N, et al. Parallel murine and human plaque proteomics reveals pathways of plaque rupture. Circulation research. 2020;127(8):997–1022.

24. Colombo M, Raposo G, Théry C. Biogenesis, secretion, and intercellular interactions of exosomes and other extracellular vesicles. Annual review of cell and developmental biology. 2014; 30:255–289.

25. Théry C, Witwer KW, Aikawa E, et al. Minimal information for studies of extracellular vesicles 2018 (MISEV2018): a position statement of the International Society for Extracellular Vesicles and update of the MISEV2014 guidelines. Journal of extracellular vesicles. 2018;7(1):1535750.

26. Welsh JA, Goberdhan DC, O’Driscoll L, et al. Minimal information for studies of extracellular vesicles (MISEV2023): From basic to advanced approaches. Journal of extracellular vesicles. 2024;13(2):e12404.

27. Leroyer AS, Isobe H, Lesèche G, et al. Cellular origins and thrombogenic activity of microparticles isolated from human atherosclerotic plaques. Journal of the American College of Cardiology. 2007;49(7):772–777.

28. Rautou P-E, Leroyer AS, Ramkhelawon B, et al. Microparticles from human atherosclerotic plaques promote endothelial ICAM-1–dependent monocyte adhesion and transendothelial migration. Circulation research. 2011;108(3):335–343.

29. Haraszti RA, Didiot M-C, Sapp E, et al. High-resolution proteomic and lipidomic analysis of exosomes and microvesicles from different cell sources. Journal of extracellular vesicles. 2016;5(1):32570.

30. Stamatikos A, Knight E, Vojtech L, et al. Exosome-mediated transfer of anti-miR-33a-5p from transduced endothelial cells enhances macrophage and vascular smooth muscle cell cholesterol efflux. Human gene therapy. 2020;31(3-4):219–232.

31. Blaser MC, Buffolo F, Halu A, et al. Multiomics of tissue extracellular vesicles identifies unique modulators of atherosclerosis and calcific aortic valve stenosis. Circulation. 2023;148(8):661–678.

32. Valadi H, Ekström K, Bossios A, Sjöstrand M, Lee JJ, Lötvall JO. Exosome-mediated transfer of mRNAs and microRNAs is a novel mechanism of genetic exchange between cells. Nature cell biology. 2007;9(6):654–659.

33. Jiang S, Li X, Li Y, et al. APOE from patient-derived astrocytic extracellular vesicles alleviates neuromyelitis optica spectrum disorder in a mouse model. Science Translational Medicine. 2024;16(736):eadg5116.

34. Antounians L, Catania VD, Montalva L, et al. Fetal lung underdevelopment is rescued by administration of amniotic fluid stem cell extracellular vesicles in rodents. Science translational medicine. 2021;13(590):eaax5941.

35. Barrett KM, Brott TG. Stroke caused by extracranial disease. Circulation research. 2017;120(3):496–501.

36. Goncalves I, Sun J, Tengryd C, et al. Plaque vulnerability index predicts cardiovascular events: a histological study of an endarterectomy cohort. Journal of the American Heart Association. 2021;10(15):e021038.

37. Matic LP, Jesus Iglesias M, Vesterlund M, et al. Novel multiomics profiling of human carotid atherosclerotic plaques and plasma reveals biliverdin reductase B as a marker of intraplaque hemorrhage. JACC: Basic to Translational Science. 2018;3(4):464–480.

38. Stary HC, Chandler AB, Dinsmore RE, et al. A definition of advanced types of atherosclerotic lesions and a histological classification of atherosclerosis: a report from the Committee on Vascular Lesions of the Council on Arteriosclerosis, American Heart Association. Circulation. 1995;92(5):1355–1374.

39. Palasca O, Santos A, Stolte C, Gorodkin J, Jensen LJ. TISSUES 2.0: an integrative web resource on mammalian tissue expression. Database. 2018;2018:bay003.

40. Forster DT, Li SC, Yashiroda Y, et al. BIONIC: biological network integration using convolutions. Nature Methods. 2022;19(10):1250–1261.

41. Hong P, Koza S, Bouvier ES. A review size-exclusion chromatography for the analysis of protein biotherapeutics and their aggregates. Journal of liquid chromatography & related technologies. 2012;35(20):2923–2950.

42. Petersen JD, Mekhedov E, Kaur S, Roberts DD, Zimmerberg J. Endothelial cells release microvesicles that harbour multivesicular bodies and secrete exosomes. Journal of Extracellular Biology. 2023;2(4):e79.

43. Organization WH. Strategy for optimizing national routine health information systems: strengthening routine health information systems to deliver primary health care and universal health coverage. World Health Organization; 2024.

44. Cheng HS, Besla R, Li A, et al. Paradoxical suppression of atherosclerosis in the absence of microRNA-146a. Circulation research. 2017;121(4):354–367.

45. Cheng HS, Sivachandran N, Lau A, et al. Micro RNA-146 represses endothelial activation by inhibiting pro-inflammatory pathways. EMBO molecular medicine. 2013;5(7):1017–1034.

46. Raitoharju E, Lyytikäinen L-P, Levula M, et al. miR-21, miR-210, miR-34a, and miR-146a/b are up- regulated in human atherosclerotic plaques in the Tampere Vascular Study. Atherosclerosis. 2011;219(1):211–217.

47. Nazari-Jahantigh M, Wei Y, Noels H, et al. MicroRNA-155 promotes atherosclerosis by repressing Bcl6 in macrophages. The Journal of clinical investigation. 2012;122(11):4190–4202.

48. Yang Y, Yang L, Liang X, Zhu G. MicroRNA-155 promotes atherosclerosis inflammation via targeting SOCS1. Cellular Physiology and Biochemistry. 2015;36(4):1371–1381.

49. Huang S, Zeng Z, Sun Y, et al. Association study of hsa_circ_0001946, hsa-miR-7-5p and PARP1 in coronary atherosclerotic heart disease. International journal of cardiology. 2021;328:1–7.

50. Reddy MA, Jin W, Villeneuve L, et al. Pro-inflammatory role of microrna-200 in vascular smooth muscle cells from diabetic mice. Arteriosclerosis, thrombosis, and vascular biology. 2012;32(3):721–729.

51. Zhang F, Cheng N, Du J, Zhang H, Zhang C. MicroRNA-200b-3p promotes endothelial cell apoptosis by targeting HDAC4 in atherosclerosis. BMC cardiovascular disorders. 2021; 21:1–12.

52. An T-H, He Q-W, Xia Y-P, et al. MiR-181b antagonizes atherosclerotic plaque vulnerability through modulating macrophage polarization by directly targeting Notch1. Molecular Neurobiology. 2017; 54:6329–6341.

53. Di Gregoli K, Mohamad Anuar NN, Bianco R, et al. MicroRNA-181b controls atherosclerosis and aneurysms through regulation of TIMP-3 and elastin. Circulation research. 2017;120(1):49–65.

54. Sun P, Li L, Liu Y-Z, et al. MiR-181b regulates atherosclerotic inflammation and vascular endothelial function through Notch1 signaling pathway. European Review for Medical & Pharmacological Sciences. 2019; 23(7).

55. Kuleshov MV, Jones MR, Rouillard AD, et al. Enrichr: a comprehensive gene set enrichment analysis web server 2016 update. Nucleic acids research. 2016;44(W1):W90–W97.

56. Hallal S, Tűzesi Á, Grau GE, Buckland ME, Alexander KL. Understanding the extracellular vesicle surface for clinical molecular biology. Journal of extracellular vesicles. 2022;11(10):e12260.

57. He R, Zhu J, Ji P, Zhao F. SEVtras delineates small extracellular vesicles at droplet resolution from single-cell transcriptomes. Nature Methods. 2024;21(2):259–266.

58. Wang S, Aurora AB, Johnson BA, et al. The endothelial-specific microRNA miR-126 governs vascular integrity and angiogenesis. Developmental cell. 2008;15(2):261–271.

59. Fish JE, Santoro MM, Morton SU, et al. miR-126 regulates angiogenic signaling and vascular integrity. Developmental cell. 2008;15(2):272–284.

60. Vasa-Nicotera M, Chen H, Tucci P, et al. miR-146a is modulated in human endothelial cell with aging. Atherosclerosis. 2011;217(2):326–330.

61. McCann JV, Xiao L, Kim DJ, et al. Endothelial miR-30c suppresses tumor growth via inhibition of TGF-β-induced Serpine1. J Clin Invest. 2019;129(4):1654–1670.

62. Yang ZJ, Liu R, Han XJ, et al. Knockdown of the long non-coding RNA MALAT1 ameliorates TNF-α-mediated endothelial cell pyroptosis via the miR-30c-5p/Cx43 axis. Mol Med Rep. 2023;27(4).

63. Afzal TA, Luong LA, Chen D, et al. NCK Associated Protein 1 Modulated by mi RNA-214 Determines Vascular Smooth Muscle Cell Migration, Proliferation, and Neointima Hyperplasia. Journal of the American Heart Association. 2016;5(12):e004629.

64. Cordes KR, Sheehy NT, White MP, et al. miR-145 and miR-143 regulate smooth muscle cell fate and plasticity. Nature. 2009;460(7256):705–710.

65. Cheng Y, Liu X, Yang J, et al. MicroRNA-145, a novel smooth muscle cell phenotypic marker and modulator, controls vascular neointimal lesion formation. Circulation research. 2009;105(2):158–166.

66. O’Connell RM, Zhao JL, Rao DS. MicroRNA function in myeloid biology. *Blood*, The Journal of the American Society of Hematology. 2011;118(11):2960–2969.

67. Das A, Ganesh K, Khanna S, Sen CK, Roy S. Engulfment of apoptotic cells by macrophages: a role of microRNA-21 in the resolution of wound inflammation. The Journal of Immunology. 2014;192(3):1120–1129.

68. Maurya M, Barthwal MK. MicroRNA-99a: a potential double-edged sword targeting macrophage inflammation and metabolism. Cellular & Molecular Immunology. 2021;18(9):2290–2292.

69. Rodriguez A, Vigorito E, Clare S, et al. Requirement of bic/microRNA-155 for normal immune function. Science. 2007;316(5824):608-611.

70. Seeley JJ, Baker RG, Mohamed G, et al. Induction of innate immune memory via microRNA targeting of chromatin remodelling factors. Nature. 2018;559(7712):114-119.

71. Jin S, Guerrero-Juarez CF, Zhang L, et al. Inference and analysis of cell-cell communication using CellChat. Nature communications. 2021;12(1):1088.

72. Consor Tum TTS, Jones RC, Karakinos J, et al. The Tabula Sapiens: A multiple-organ, single-cell transcriptomic atlas of humans. Science. 2022;376(6594):eabl4896.

73. Hsu S-D, Lin F-M, Wu W-Y, et al. miRTarBase: a database curates experimentally validated microRNA–target interactions. Nucleic acids research. 2011;39(suppl_1):D163–D169.

74. Stary HC. Natural history and histological classification of atherosclerotic lesions: an update. Arteriosclerosis, thrombosis, and vascular biology. 2000;20(5):1177–1178.

75. Adya R, Tan BK, Punn A, Chen J, Randeva HS. Visfatin induces human endothelial VEGF and MMP-2/9 production via MAPK and PI3K/Akt signalling pathways: novel insights into visfatin- induced angiogenesis. Cardiovascular research. 2008;78(2):356–365.

76. Bae Y-H, Bae M-K, Kim S-R, Lee JH, Wee H-J, Bae S-K. Upregulation of fibroblast growth factor-2 by visfatin that promotes endothelial angiogenesis. Biochemical and biophysical research communications. 2009;379(2):206–211.

77. Adya R, Tan BK, Chen J, Randeva HS. Pre-B cell colony enhancing factor (PBEF)/visfatin induces secretion of MCP-1 in human endothelial cells: role in visfatin-induced angiogenesis. Atherosclerosis. 2009;205(1):113–119.

78. Adya R, Tan BK, Chen J, Randeva HS. Nuclear factor-κB induction by visfatin in human vascular endothelial cells: its role in MMP-2/9 production and activation. Diabetes care. 2008;31(4):758–760.

79. Kim S-R, Bae Y-H, Bae S-K, et al. Visfatin enhances ICAM-1 and VCAM-1 expression through ROS-dependent NF-κB activation in endothelial cells. Biochimica et Biophysica Acta (BBA)-Molecular Cell Research. 2008;1783(5):886–895.

80. Sinning J-M, Losch J, Walenta K, Böhm M, Nickenig G, Werner N. Circulating CD31+/Annexin V+ microparticles correlate with cardiovascular outcomes. European heart journal. 2011;32(16):2034–2041.

81. Arteaga RB, Chirinos JA, Soriano AO, et al. Endothelial microparticles and platelet and leukocyte activation in patients with the metabolic syndrome. The American journal of cardiology. 2006;98(1):70–74.

82. Chironi G, Simon A, Hugel B, et al. Circulating leukocyte-derived microparticles predict subclinical atherosclerosis burden in asymptomatic subjects. Arteriosclerosis, thrombosis, and vascular biology. 2006;26(12):2775–2780.

83. Koga H, Sugiyama S, Kugiyama K, et al. Elevated levels of VE-cadherin-positive endothelial microparticles in patients with type 2 diabetes mellitus and coronary artery disease. Journal of the American College of Cardiology. 2005;45(10):1622–1630.

84. Amabile N, Cheng S, Renard JM, et al. Association of circulating endothelial microparticles with cardiometabolic risk factors in the Framingham Heart Study. European Heart Journal. 2014;35(42):2972–2979.

85. Crescitelli R, Lässer C, Lötvall J. Isolation and characterization of extracellular vesicle subpopulations from Tissues. Nature Protocols. 2021;16(3):1548–1580.

86. Dixson AC, Dawson TR, Di Vizio D, Weaver AM. Context-specific regulation of extracellular vesicle biogenesis and cargo selection. Nature Reviews Molecular Cell Biology. 2023;24(7):454–476.

87. Shankman LS, Gomez D, Cherepanova OA, et al. KLF4-dependent phenotypic modulation of smooth muscle cells has a key role in atherosclerotic plaque pathogenesis. Nature medicine. 2015;21(6):628–637.

88. Pan H, Xue C, Auerbach BJ, et al. Single-cell genomics reveals a novel cell state during smooth muscle cell phenotypic switching and potential therapeutic targets for atherosclerosis in mouse and human. Circulation. 2020;142(21):2060–2075.

89. Bennett MR, Sinha S, Owens GK. Vascular smooth muscle cells in atherosclerosis. Circulation research. 2016;118(4):692–702.

90. Fernandez DM, Rahman AH, Fernandez NF, et al. Single-cell immune landscape of human atherosclerotic plaques. Nature medicine. 2019;25(10):1576–1588.

91. Milei J, Parodi J, Barone A, et al. Carotid atherosclerosis. Immunocytochemical analysis of the vascular and cellular composition in endarterectomies. Cardiologia (Rome, Italy). 1996;41(6):535–542.

92. Soehnlein O, Libby P. Targeting inflammation in atherosclerosis—from experimental insights to the clinic. Nature reviews Drug discovery. 2021;20(8):589–610.

93. Libby P. The biology of atherosclerosis comes full circle: lessons for conquering cardiovascular disease. Nature Reviews Cardiology. 2021;18(10):683–684.

94. Gao P, Chen Z-Q, Bao Y-H, Jiao L-Q, Ling F. Correlation between carotid intraplaque hemorrhage and clinical symptoms: systematic review of observational studies. Stroke. 2007;38(8):2382–2390.

95. Parma L, Baganha F, Quax PH, de Vries MR. Plaque angiogenesis and intraplaque hemorrhage in atherosclerosis. European journal of pharmacology. 2017;816:107–115.

96. Wadén K, Karlöf E, Narayanan S, et al. Clinical risk scores for stroke correlate with molecular signatures of vulnerability in symptomatic carotid patients. Iscience. 2022;25(5).

97. Larson AS, Benson JC, Brinjikji W, et al. Variations in the presence of carotid Intraplaque hemorrhage across age categories: what age groups are Most likely to benefit from plaque imaging? Frontiers in neurology. 2020; 11:603055.

98. Arderiu G, Peña E, Badimon L. Angiogenic microvascular endothelial cells release microparticles rich in tissue factor that promotes postischemic collateral vessel formation. Arteriosclerosis, thrombosis, and vascular biology. 2015;35(2):348–357.

99. Boyer MJ, Kimura Y, Akiyama T, et al. Endothelial cell-derived extracellular vesicles alter vascular smooth muscle cell phenotype through high-mobility group box proteins. Journal of extracellular vesicles. 2020;9(1):1781427.

100. Charla E, Mercer J, Maffia P, Nicklin S. Extracellular vesicle signalling in atherosclerosis. Cellular signalling. 2020;75:109751.

101. He S, Wu C, Xiao J, Li D, Sun Z, Li M. Endothelial extracellular vesicles modulate the macrophage phenotype: Potential implications in atherosclerosis. Scandinavian journal of immunology. 2018;87(4):e12648.

102. Jansen F, Yang X, Baumann K, et al. Endothelial microparticles reduce ICAM-1 expression in a micro RNA-222-dependent mechanism. Journal of cellular and molecular medicine. 2015;19(9):2202–2214.

103. Liang H, Li S, Chen H. GW28-e0938 Endothelial microparticles-mediated transfer of microRNA- 19b, a novel messenger in cell-cell communication, plays a key role in endothelial migration and angiogenesis. Journal of the American College of Cardiology. 2017;70(16S):C34–C34.

104. Njock M-S, Cheng HS, Dang LT, et al. Endothelial cells suppress monocyte activation through secretion of extracellular vesicles containing antiinflammatory microRNAs. *Blood*, The Journal of the American Society of Hematology. 2015;125(20):3202–3212.

105. Botts SR, Fish JE, Howe KL. Dysfunctional vascular endothelium as a driver of atherosclerosis: emerging insights into pathogenesis and treatment. Frontiers in pharmacology. 2021;12:787541.

106. Mokry M, Boltjes A, Slenders L, et al. Transcriptomic-based clustering of human atherosclerotic plaques identifies subgroups with different underlying biology and clinical presentation. Nature cardiovascular research. 2022;1(12):1140–1155.

107. Seeley EH, Liu Z, Yuan S, et al. Spatially resolved metabolites in stable and unstable human atherosclerotic plaques identified by mass spectrometry imaging. Arteriosclerosis, thrombosis, and vascular biology. 2023;43(9):1626–1635.

108. AbuRahma AF, Avgerinos ED, Chang RW, et al. Society for Vascular Surgery clinical practice guidelines for management of extracranial cerebrovascular disease. Journal of vascular surgery. 2022;75(1):4S–22S.

109. Gaba K, Syed M, Raza Z. Reducing the delay for carotid endarterectomy in South-East Scotland. The Surgeon. 2014;12(1):11–16.

110. Gocan S, Bourgoin A, Shamloul R, Sivakumar B, Dowlatshahi D, Stotts G. Early vascular imaging and key system strategies expedite carotid revascularization aRer transient ischemic attack and stroke. Journal of Vascular Surgery. 2020;72(5):1728–1734.

111. Lim ET, Laws PE, Beresford TP, Khanafer AM, Roake JA. Timeliness and outcomes of carotid endarterectomy for symptomatic carotid artery stenosis: a single centre audit. ANZ Journal of Surgery. 2020;90(3):345–349.

112. Den Hartog A, Moll F, van der Worp H, Hoff R, Kappelle L, de Borst G. Delay to carotid endarterectomy in patients with symptomatic carotid artery stenosis. European Journal of Vascular and Endovascular Surgery. 2014;47(3):233–239.

113. Meyer D, Karreman E, Kopriva D. Factors associated with delay in carotid endarterectomy for patients with symptomatic severe internal carotid artery stenosis: a case–control study. Canadian Medical Association Open Access Journal. 2018;6(2):E211–E217.

114. Barnett HJ, Taylor DW, Eliasziw M, et al. Benefit of carotid endarterectomy in patients with symptomatic moderate or severe stenosis. New England Journal of Medicine. 1998;339(20):1415–1425.

115. Peeters W, Hellings W, de Kleijn D, et al. Carotid Atherosclerotic Plaques Stabilize ARer Stroke. 2008.

116. Ching T, Huang S, Garmire LX. Power analysis and sample size estimation for RNA-Seq differential expression. Rna. 2014;20(11):1684–1696.

117. Movat HZ. Demonstration of all connective tissue elements in a single section; pentachrome stains. AMA archives of pathology. 1955;60(3):289–295.

118. TiÅord M. The long history of hematoxylin. Biotechnic & histochemistry. 2005;80(2):73–78.

119. Tyanova S, Temu T, Sinitcyn P, et al. The Perseus computational plaÅorm for comprehensive analysis of (prote) omics data. Nature methods. 2016;13(9):731–740.

120. Jeppesen DK, Fenix AM, Franklin JL, et al. Reassessment of exosome composition. Cell. 2019;177(2):428–445. e418.

121. Emoto T, Lu J, Sivasubramaniyam T, et al. Colony stimulating factor-1 producing endothelial cells and mesenchymal stromal cells maintain monocytes within a perivascular bone marrow niche. Immunity. 2022;55(5):862–878. e868.

122. Love MI, Huber W, Anders S. Moderated estimation of fold change and dispersion for RNA-seq data with DESeq2. Genome biology. 2014;15(12):1–21.

123. Benjamini Y, Hochberg Y. Controlling the false discovery rate: a practical and powerful approach to multiple testing. Journal of the Royal statistical society: series B (Methodological). 1995;57(1):289–300.

124. Huang H-Y, Lin Y-C-D, Li J, et al. miRTarBase 2020: updates to the experimentally validated microRNA–target interaction database. Nucleic acids research. 2020;48(D1):D148–D154.

125. Licursi V, Conte F, Fiscon G, Paci P. MIENTURNET: an interactive web tool for microRNA-target enrichment and network-based analysis. BMC bioinformatics. 2019;20:1–10.

126. Chen EY, Tan CM, Kou Y, et al. Enrichr: interactive and collaborative HTML5 gene list enrichment analysis tool. BMC bioinformatics. 2013;14:1–14.

127. Xie Z, Bailey A, Kuleshov MV, et al. Gene set knowledge discovery with Enrichr. Current protocols. 2021;1(3):e90.

128. Christian LS, Wang L, Lim B, et al. Resident memory T cells in tumor-distant Tissues fortify against metastasis formation. Cell reports. 2021;35(6).

129. Bafor EE, Valencia JC, Young HA. Double negative T regulatory cells: an emerging paradigm shiR in reproductive immune tolerance? Frontiers in Immunology. 2022;13:886645.

130. Xu MM, Pu Y, Han D, et al. Dendritic cells but not macrophages sense tumor mitochondrial DNA for cross-priming through signal regulatory protein α signaling. Immunity. 2017;47(2):363–373. e365.

131. Alqassim EY, Sharma S, Khan ANH, et al. RNA editing enzyme APOBEC3A promotes pro- inflammatory M1 macrophage polarization. Communications biology. 2021;4(1):102.

132. Basatemur GL, Jørgensen HF, Clarke MC, Bennett MR, Mallat Z. Vascular smooth muscle cells in atherosclerosis. Nature reviews cardiology. 2019;16(12):727–744.

133. Muhl L, Genové G, Leptidis S, et al. Single-cell analysis uncovers fibroblast heterogeneity and criteria for fibroblast and mural cell identification and discrimination. Nature communications. 2020;11(1):3953.

134. John L, Poos AM, Brobeil A, et al. Resolving the spatial architecture of myeloma and its microenvironment at the single-cell level. Nature Communications. 2023;14(1):5011.

135. Arora R, Cao C, Kumar M, et al. Spatial transcriptomics reveals distinct and conserved tumor core and edge architectures that predict survival and targeted therapy response. Nature Communications. 2023;14(1):5029.

136. Sinha S, Rosin NL, Arora R, et al. Dexamethasone modulates immature neutrophils and interferon programming in severe COVID-19. Nature medicine. 2022;28(1):201–211.

