## Supplementary Material for "Multiomics unveils extracellular vesicle-driven mechanisms of endothelial communication in human carotid atherosclerosis"

Raju S, et al.

#### Methods

##### Clinical criteria for patient selection for carotid atherosclerotic plaque

Decisions for carotid endarterectomy (CEA) were based according to surgeon +/- multidisciplinary discussion (e.g. stroke neurology, neuroradiology) assessment alongside established clinical guidelines.[108] Briefly, symptomatic patients with extracranial carotid atheroembolic disease identified to vascular surgeons were selected on the basis of a clinically relevant event (i.e. amaurosis fugax, transient ischemic attack or stroke) with a confirmed radiographic diagnosis of the culprit lesion in the carotid artery (i.e. duplex ultrasound and computed tomography (CT) or magnetic resonance (MR) angiogram) +/- brain imaging (CT or MRI) in the absence of an alternate source (i.e. intracerebral haemorrhage or atrial fibrillation) within 30 days of their last neurologic event. *See below for additional details and surgical timing. Asymptomatic patients with no symptoms in the previous 6 months undergoing operative intervention based upon the surgeon’s recommendation (e.g. progressive stenosis in a young, otherwise medically optimized patient or severe stenosis with contralateral carotid occlusion) were included provided there was no history of radiation, prior ipsilateral endarterectomy, active cancer or infection.

**Timing:* As per clinical guidelines, symptomatic patients with a culprit carotid plaque lesion are recommended to undergo CEA within 2 weeks. Despite this goal, there are limitations (patient and system-derived factors)[109] to achieving revascularization within the 2 week period in many countries.[110-113] As per NASCET criteria[114], symptomatic disease is defined by an ipsilateral neurological event within the preceding 6 months. However, ruptured carotid plaques can heal during this 6 month period.[115] To better understand the biology of carotid plaques from symptomatic patients, plaque collection from this group was restricted to 4 weeks post neurological deficits, in order to capture sufficient patient samples while balancing system issues that can delay timing for intervention. Asymptomatic patients underwent CEA as part of elective surgical practice.
Patient carotid plaque endarterectomy samples and clinical data were obtained from the University Health Network and Sunnybrook Health Sciences Centre, Toronto, Canada in accordance with institutional research ethics board protocol (18-5282). The study and all protocols conform to the ethical guidelines set forth by the Helsinki Declaration (version 2008).
*Inclusion Criteria*

- Adult patients undergoing carotid endarterectomy (symptomatic or asymptomatic)
- Symptomatic plaques were defined as patients presenting with transient or permanent focal neurological deficits including ipsilateral amaurosis fugax, expressive or receptive aphasia, and/or dysarthria, contralateral paraesthesia, paresis, or paralysis of the face, upper extremity, and/or lower extremity, <4 weeks since initial onset.

*Exclusion Criteria*

- Pregnant or age <18 years
- Non-extracranial carotid artery sources of ischemic symptoms (e.g. sources from cardioembolic disease, haemorrhagic stroke, lacunar infarction)
- Prior neck radiation
- Re-do carotid endarterectomy on ipsilateral side
- Patients who met symptomatic criteria >4 weeks since last neurologic event (re: NASCET criteria would categorize a patient as symptomatic if neurologic event was within 6 months)
- Active cancer or systemic infection

**Human carotid plaque collection and processing**

Immediately after endarterectomy, carotid plaques were suspended in Hanks Balanced Salt Solution (Thermo Fisher Scientific, Waltham, MA, USA) and within an hour grossly dissected into two defined regions by a single trained surgeon (Dr. Sneha Raju): (i) plaque, and (ii) marginal zone(Figure 1A, S4A). Regions were delineated based on presence (plaque zone) or absence (marginal zone) of visible atheromatous disease. Tissues (separated into plaque and marginal zones) were weighed, a small segment of plaque was sectioned for histological review, and the reminder of sample was snap frozen in liquid nitrogen and placed in a -80°C for downstream processing. Adjacent carotid tissue (marginal zone) was used as an internal control for carotid plaque zone samples. The utilization of patient-matched tissue samples allowed marginal zone signatures to be used as an internal control that accounts for interpatient heterogeneity unrelated to plaque-rupture specific biology. Previous studies have shown that patient matched marginal tissue increases statistical power for differential gene expression.[116]

A total of 64 patient carotid endarterectomy samples were used in this study as detailed in Methods Figure 1. A total of 29 paired (plaque zone and matched marginal zone) patient samples were used for extracellular vesicle (EV)-vesiculomics (miRNA transcriptomics and proteomics). Separate samples were used to complete whole tissue proteomics (N=5), EV enrichment validation, quantification, visualization (cryogenic TEM), proteomics method validation, histopathology and *in vitro* experiments. Demographics and comorbidities were available for a subset of patients and are displayed in Table 1.

**Tissue Histology**

Tissue for histology was fixed in 4% paraformaldehyde for 24 hours, dehydrated in alcohols of increasing concentrations (50%, 70%, and 96%), processed in xylene (50-100 ml), immobilized in paraffin blocks, and sectioned into 5 µm thick sheets with a rotary microtome. The sheets were then placed on microscopy slides and stained with the Russell-Movat[117] and Hematoxylin and eosin stain.[118] Several samples were assessed and graded in a blinded fashion by a staff pathologist (Dr. Seidman). All images acquired on a Leica DM2500 LED microscope with a 2.5x Fluotar objective, using an OMAX A35180U3 digital microscope camera, 0.70x C-mount adapter, and ToupView software. Images were sharpened, annotated, and converted to TIFF format using the GNU Image Manipulator Program (GIMP) 2.10.14 for Windows.

**Tissue proteomics**

*Tissue proteomics processing*

Keeping the carotid endarterectomy samples frozen, tissue samples were processed into ~ 1$mm^3$ pieces using a blade. Tissue pieces were then pulverized in liquid nitrogen using a pestle and mortar. Protein was extracted by resuspending the pulverized tissue in RIPA buffer (ThermoFisher) supplemented with protease and phosphatase inhibitors (Roche, cOmplete Mini tablets and PhosSTOP), vortexing, incubating on wet ice for 1 hr followed by additional vortexing and 10 mins sonication in a benchtop ultrasonic water bath (Branson 5510). Samples were then centrifuged at 2000 x g for 5 min. The supernatant was collected and transferred to a new tube. Samples were stored at -80°C for liquid chromatography tandem mass spectrometry (LC-MS/MS). The protein concentration was determined using the Bicinchoninic acid (BCA) assay on a Nanodrop spectrophotometer reading at 562 nm against a standard curve. 10 µg of protein was used for proteomics preparation.

*Sample preparation*

Peptides for MS injection were prepared to the PreOmics iST kit protocol. In LYSE buffer, samples were heated at 95 °C for 10 minutes. Protein samples were digested using PreOmics provided Lys-C/trypsin DIGEST solution for 2 hours at 37 °C. Samples were speed-vacuumed at 45°C only until the peptides were dried and was subsequently stored at -80°C until LC-MS/MS sequencing. 17µL of LC-LOAD solution (PreOmics) was added to the dried peptides. Samples were centrifuged at 16 000 x g for 1 minute and then 15 µL were aspirated into a new tube and 2 µL was injected.

*Mass spectrometry*

An Orbitrap Exploris 480 mass spectrometer fronted with an EASY-Spray Source (heated at 45 ˚C), coupled to an Easy-nLC1200 HPLC pump (Thermo Scientific) was used to analyse the peptide samples. Peptides were fractionated using a dual column set-up: an Acclaim™ PepMap™ 100 C18 HPLC Columns, 75 µm X 70 mm; and an EASY-Spray™ HPLC Column, 75 µm X 250 mm. The analytical gradient was run at 300 nL/min from 5% to 21% Solvent B (acetonitrile/0.1% formic acid) for 60 minutes, followed by 10 minutes of 21% to 30% Solvent B, and another 10 minutes of a 95%-5% jigsaw wash. The MS1 resolution was set to 120 K (maximum injection time, 25 ms), and the TopS precursor ions (within a 3.0 second cycle time) were subject to high-energy collision induces dissociation (HCD) isolation width 1.2 m/z, dynamic exclusion enabled (60 seconds), and MS2 resolution was set to 60 K (maximum injection time, auto; normalized collision energy, 24%, 26% and 28%). The MS1 and MS2 acquisition range was set to 400 m/z – 1500 m/z and 120 m/z – 1200 m/z, respectively.

*Mass spectral annotation*

The acquired peptide spectra from the 12 samples were searched with Proteome Discoverer package (PD, Version 2.5) using the SEQUEST-HT search algorithm against the Human UniProt database (101043 entries, updated January 2022) to identify the proteins in the samples. Trypsin was set as the digestion enzyme, allowing up to 2 missed cleavages and a minimum peptide length of 6 amino acids. Oxidation (+15.995 Da) of methionine; and acetylation (+42.011 Da) of the N-terminus, were set as variable modifications. Carbamidomethylation (+57.021 Da) of cysteine was set as a static modification. Furthermore, the precursor tolerance was set to 10 ppm and the fragment tolerance window to 0.02 Da. Peptides were filtered based on a false discovery rate of 1.0%, which was calculated using Percolator, provided by Proteome Discover and peptides were filtered based on a 1.0% FDR. Quantification utilized unique peptides (those assigned to a given Master protein group and not present in any other protein group) and razor peptides (peptides shared among multiple protein groups). Razor peptides were used to quantify only the protein with the most identified peptides and not for the other proteins they are contained in. The “Feature Mapper” was enabled in PD to identify peptide precursors that may not have been sequenced in all samples but were detected in the MS1. The chromatographic spectra were aligned while allowing for a maximum retention time shift of 10 minutes, mass tolerance of 10 ppm, and a minimum signal-to-noise ratio of 5.

Chromatographic intensities were used to establish precursor peptide abundance. Peptide abundance was normalized by total peptide amount. Each protein intensity was calculated using the sum of its peptide intensities. A minimum of at least 2 unique peptides for each protein was required for the protein to be included in the analyses.

*Proteome Analysis*
The quantified proteins were exported from Proteome Discoverer and further processed using Perseus.[119] Proteins were removed from analysis if they were not present in at least 9 of the 12 samples. The datasets were median normalized, log2 transformed, and missing data was imputed using a Gaussian distribution within each sample with a 1.96 standard deviation downshift within a 0.3 standard deviation range. Principal component analysis (PCA) and two-group differential enrichment analysis was performed using Qlucore Omics Explorer statistical software (Version 3.9). PCAs were unbiased and not statistically thresholded. For comparisons of the EV proteome of the plaque and the marginal region, the effect of donor was subtracted using a generalized linear model. Significantly differentially enriched proteins were calculated using a two-group comparison with a Benjamini-Hochberg false discovery rate (FDR)-corrected p-value (i.e., q-value) less than 0.05. Pathway analysis on differentially enriched proteins was completed as described below.

**Extracellular vesicle (EV) enrichment from snap frozen human carotid plaque tissue**

Crude EVs were isolated from the interstitial space of human tissue as previously established, with modifications.[85,120] Briefly, pieces of frozen carotid plaque or marginal zone tissue were thawed, roughly split into 1-5 $mm^3$ pieces, and underwent enzymatic digestion at 37°C in 250 units/mL Collagenase type I (Worthington LS004210) for 45-60 minutes. Subsequently, entire sample was dissociated using gentle mortar and pestle pulverization, centrifuged (400 x g, 4°C 10 minutes) and strained through a 0.8 mM syringe filter (Millipore Sigma SLAA033SS) to remove cells and debris.

EVs were then enriched from the supernatant using by size exclusion chromatography (SEC) for miRNA experiments with an added density gradient ultracentrifugation step for proteomics (detailed below). SEC was completed using the legacy qEV 70 nm columns (Izon Science Ltd, Christchurch, NZ). Supernatants were concentrated to 500 µL using Amicon 10K MWCO filters and loaded onto the 10 ml bed volume qEV columns, followed by elution with 0.22 µM filtered PBS-/-. After collection of a 3 mL void volume, four fractions of 500 µL volumes were collected from the column. The EV-enriched fractions were concentrated to 25-50 µL via ultrafiltration using the Amicon 10K MWCO filters (Millipore, UFC801096D) and validated as described below.

**EV validation and characterization**

EV isolation was confirmed via western blot analysis of common EV markers as described previously.[8] Briefly, EV samples were lysed in 5X RIPA prior to protein quantification via Micro BCA (ThermoFisher, 23235) and SDS gel electrophoresis. Proteins were separated on 4-20% precast polyacrylamide gels (Mini-Protean TGX Precast Protein gels, 4561094), transferred to PVDF membranes, blocked, and probed for markers of interest. Primary antibodies utilized were anti-CD63 (ABclonal, A5271), anti-CD81 (ABclonal, A5270), anti-CD9 (SantaCruz, sc-13118), anti-Flotillin 1 (SantaCruz, sc-74566) and anti-Calnexin (ABclonal, A4846). Membranes were then incubated with anti-rabbit or anti-mouse HRP (Cell Signalling, 7074S, 7076S), developed using SuperSignal West Femto Maximum Sensitivity Substrate ECL (ThermoFisher, 34094), and imaged on the Bio-Rad ChemiDoc system.

Following SEC, EV-enriched samples were quantified and visualized via nanoparticle tracking analysis (NTA, NS300 Malvern Panalytical Ltd, 532 nm laser) and cryogenic transmission microscopy (cryo-EM) as previously detailed.[8] Briefly, samples for NTA were diluted to a concentration between $10^6$-$10^9$ particles/mL prior to recording and three videos (60 sec duration each) were recorded for each sample and measurements averaged across runs by NTA software (NTA 3.4 Build 3.4.4). Samples for cryo-EM were vitrified in liquid ethane using a Vitrobot Mark IV (Thermo Grids were imaged with a Talos L120C TEM (Thermo Scientific) using a high tension of 120 kV with a 4k x 4k BM-Ceta CMOS camera. At least 10 images of each sample were taken at magnifications 28,000x, 57,000x, and 120,000x yielding a pixel size of 510 pm, 249 pm, and 121 pm, respectively.

Assessment of lipoprotein co-isolation was completed using the Quantikine Human Apolipoprotein B (R&D Systems, DAPB00) and Apolipoprotein A1 (DAPA10) solid phase ELISA (enzyme-linked immunosorbent assay), as per the manufacturer's instructions. Undiluted SEC fractions were utilized. The lack of whole cell contamination was validated via flow cytometry analysis with EV-enriched samples and bone marrow cells isolated from mouse femur[121] stained with Hoechst 33342 (0.6 μg/mL; ThermoFisher Scientific 62249).

**EV MicroRNA (miRNA) Sequencing and Analysis**

EVs were enriched via SEC from paired plaque and marginal zones from asymptomatic and symptomatic patient cohorts (detailed in Methods Figure 1). HTG Molecular Diagnostics EdgeSeq technology (HTG Molecular Diagnostics, Inc, Tucson, AZ, USA) was utilized to determine concentration of 2,083 known miRNAs, as described previously.[8] Briefly, samples were prepared for sequencing by 1:1 addition of EV sample to HTG plasma lysis buffer, followed by addition of Proteinase K (Invitrogen) for 180 minutes. All samples were run using the HTG EdgeSeq miRNA WT assay, which includes 2,083 functional DNA nuclease protection probes (NPP) designed to bind to human miRNA. Library preparation was completed with DNA probe amplification using multiple PCR cycles with primers designed to recognize the NPP wing sequencies. The library was quantified using HTG EdgeSeq KAPA Library Quantification for Illumina Sequencing with samples quantified in triplicate. Post qPCR quantification, HTG EdgeSeq RUO Library Calculator was used to ensure adequate sample concentration for library pooling and dilutions. Library preparations were sequenced on the Illumina NextSeq 500/550 platform. A minimum of three biological replicates were run for all experimental groups.

**Differential EV-miRNA expression and pathway analysis**

Partek Flow (v10.0.23.0326) was used for differential miRNA expression analysis as detailed previously.[8] Adaptors (GATCGGAAGAGCACACGTCTGAACTCCAGTCACCGATGTA TCTCGTATGCCGTCTTCTGCTTG) were trimmed from the 3' end and mapped to the genome using Bowtie with the hg38 assembly (Homo Sapiens, BioProject PRJNA31257). Unaligned reads were filtered and aligned reads were quantified using miRBase mature microRNAs (version 22) annotation model with no strand specificity and 100% minimum read overlap. Principal component analysis (PCA) was performed on Qlucore Omics Explorer statistical software (version 3.9). PCAs were unbiased and not statistically thresholded. PCA was completed using median normalized counts, thresholded at 0.0001, log2 transformed, and donor effects subtracted using a generalized linear model. Differential expression of miRNA in experimental conditions was determined via DESeq2[122], without fold-change shrinkage and with adjustment for multiple comparisons using Benjamini and Hochberg's FDR step-up.[123] An adjusted p-value $< 0.05$ was considered significant. MiRNAs with ID >1000 were removed for downstream analysis. Predicted messenger RNA (mRNA) targets of differentially expressed EV-miRNA were identified using MicroRNA Enrichment Turned Network (MIENTURNET version 2019-11-25; minimum number of miRNA target interactions 1, FDR<0.05) using the miRTarBase database.[124,125]

#### EV proteomics, differential EV-protein expression, and pathway analysis

*Iodixanol density gradient ultracentrifugation EV enrichment for proteomics analysis*
Following SEC, tissue EV-enriched samples (Fractions 8 and 9) were frozen at -80°C and transferred to Brigham and Women's Hospital for downstream tissue EV enrichment processing and subsequent proteomics analysis, using a previously validated protocol for carotid artery tissue EV enrichment for proteomic analysis.[31] Samples (n=23 paired plaque and marginal zones from 13 symptomatic and 10 asymptomatic patients; detailed in Methods Figure 1) were thawed on ice and centrifuged at 10,000 x g for 10 minutes at 4°C. The supernatant was transferred, and ultracentrifuged at 100, 000 x g for 1 hr at 4°C (Beckman Coulter, Optima MAX-UP, fixed-angle rotor MLA-55) in polycarbonate ultracentrifuge tubes (Beckman Coulter). The resultant pellet containing EVs was resuspended in 1.5mL of NTE buffer (137 mM NaCl, 1 mM EDTA, 10 mM Tris, pH 7.4, 0.22um filtered) with protease inhibitor (Roche, cOmplete Mini tablets, 4693159001) and layered on top of a linear 5-step 10-30% iodixanol gradient (OptiPrep Density Gradient Media, Sigma-Aldrich D1556, and NTE buffer as diluent) with 1.5 mL per gradient step. The iodixanol gradient was moved horizontally for 1 hr at 4°C, and then vertically for at least 30 minutes before the sample was added to create a continuous gradient. The iodixanol gradient with the sample was ultracentrifuged at 250,000 x g for 40 minutes at 4°C (rotor MLA-55), and the top 2.4 mL were collected, as these have previously been shown to be the EV enriched fractions in carotid plaque tissues.[31] The sample was topped up to a volume of 9 ml with NTE buffer, and underwent ultracentrifugation at 100,000 x g for 1 hr at 4°C. Supernatant was removed and sample was resuspended in LYSE buffer (PreOmics GmbH, Planegg/Martinsried, Germany) and frozen at -80°C for downstream proteomics analysis or resuspended in NTE buffer for transmission electron microscopy (TEM) or NTA in a subset (N=3) of samples.

*Sample preparation*
From isolated EVs, peptides for mass spectrometer (MS) injection were prepared as per the PreOmics iST kit (PreOmics GmbH, Germany) protocol with modification for low abundance protein samples. In LYSE buffer, samples were heated at 95°C for 10 minutes. Protein samples were digested using PreOmics provided Lys-C/trypsin DIGEST solution for 2 hours at 37°C using 10 μL of LYSE solution (PreOmics) from sample and 10 μL of DIGEST solution. Samples were speed-vacuumed at 45°C only until the peptides were dried and was subsequently stored at -80°C until liquid chromatography tandem mass spectrometry (LC-MS/MS) sequencing. 17 mL of LC-LOAD solution (PreOmics) was added to the dried peptides. Samples were set on ice for 1 hour and then sonicated in an ultrasonic waterbath for 5 minutes. Samples were centrifuged at 16 000 x g for 1 minute and then 15 mL were aspirated into a new tube and 2 mL was injected.

*Mass spectrometry*
An Orbitrap Exploris 480 MS fronted with an EASY-Spray Source (heated at 45˚C), coupled to an Easy-nLC1200 HPLC pump (Thermo Scientific) was used to analyze the peptide samples. Peptides were fractionated using a dual column set-up: an Acclaim™ PepMap™ 100 C18 HPLC Columns, 75 µm X 70 mm; and an EASY-Spray™ HPLC Column, 75 µm X 250 mm. The analytical gradient was run at 300 nL/min from 5% to 21% Solvent B (acetonitrile/0.1% formic acid) for 60 minutes, followed by 10 minutes of 21% to 30% Solvent B, and another 10 minutes of a 95%-5% jigsaw wash. The MS1 resolution was set to 120 K (maximum injection time, 25

ms), and the TopS precursor ions (within a 3.0 second cycle time) were subject to high-energy collision induces dissociation (HCD) isolation width 1.2 m/z, dynamic exclusion enabled (60 seconds), and MS2 resolution was set to 60 K (maximum injection time, auto; normalized collision energy, 24%, 26% and 28%). The MS1 and MS2 acquisition range was set to 400 m/z – 1500 m/z and 120 m/z – 1200 m/z, respectively.

*Mass spectral annotation*

The acquired peptide spectra from the 49 samples were searched with Proteome Discoverer package (PD, Version 2.5) using the SEQUEST-HT search algorithm against the Human UniProt database (101043 entries, updated January 2022) to identify the proteins in the samples. Trypsin was set as the digestion enzyme, allowing up to 2 missed cleavages and a minimum peptide length of 6 amino acids. Oxidation (+15.995 Da) of methionine; and acetylation (+42.011 Da) of the N-terminus, were set as variable modifications. Carbamidomethylation (+57.021 Da) of cysteine was set as a static modification. Furthermore, the precursor tolerance was set to 10 ppm and the fragment tolerance window to 0.02 Da. Peptides were filtered based on a false discovery rate of 1.0%, which was calculated using Percolator, provided by PD and peptides were filtered based on a 1.0% FDR. Quantification utilized unique peptides (those assigned to a given Master protein group and not present in any other protein group) and razor peptides (peptides shared among multiple protein groups). Razor peptides were used to quantify only the protein with the most identified peptides and not for the other proteins they are contained in. The "Feature Mapper" was enabled in PD to identify peptide precursors that may not have been sequenced in all samples but were detected in the MS1. The chromatographic spectra were aligned while allowing for a maximum retention time shift of 10 minutes, mass tolerance of 10 ppm, and a minimum signal-to-noise ratio of 5. Chromatographic intensities were used to establish precursor peptide abundance. Proteins were quantified using the Total Peptide strategy provided by PD: Each protein intensity was calculated using the sum of its peptide intensities. A minimum of at least 2 unique peptides for each protein was required for the protein to be included in the analyses.

*Proteome analysis*

The quantified proteins were exported from Proteome Discoverer and further processed using Perseus.[119] Proteins were removed from analysis if they were not present in at least 70% of samples within a comparator group. The comparator groups were asymptomatic plaque (N=14), asymptomatic marginal (N=13), symptomatic plaque (N=12), asymptomatic marginal (N=10). The datasets were median normalized, log2 transformed, and missing data was imputed using a Gaussian distribution within each sample with a 1.96 standard deviation downshift within a 0.3 standard deviation range.

PCA and two-group differential enrichment analysis was performed using Qlucore Omics Explorer statistical software (Version 3.9; Qlucore, Sweden). PCAs were unbiased and not statistically thresholded. For comparisons of the EV proteome of the plaque and the marginal region, the effect of donor was subtracted using a generalized linear model and only samples with matched plaque and marginal regions (asymptomatic, N=10; symptomatic, N=13) were included in all presented analysis. Significantly differentially enriched proteins were calculated using a two-group comparison with a Benjamini-Hochberg false discovery rate (FDR)-corrected p-value (i.e., q-value) less than 0.05.

For comparison of SEC +/- density gradient ultracentrifugation (DGUC) proteomes on N=3 paired samples, a uncorrected $p<0.1$ was used threshold annotated of differentially enriched proteins to maximize sensitive of assessment of proteome changes that may have resulted from the methods. The list of conserved prototypical EV markers were generated from literature review, as previously reported.[31]

To identify EV containing SEC fractions, each fraction was centrifuged at 100,000 x g for 1hr, the fraction-specific proteome within the same sample were Z-score normalized across protein within each sample and plotted using a heat map for 4 representative tissues. These findings identified consistent enrichment in fractions 8-9 which were used in vesiculomics analysis.

*Nanoparticle tracking analysis*
Following DGUC, particle size and concentration was measured in EV enriched fractions using NTA (Malvern Instruments, NanoSight LM10). For each sample, five data collection windows (1 minute per window) were recorded during continuous injection by syringe pump with the following parameters: screen gain 1.0, 10.0 (capture, processing); camera level 9.0; detection threshold 2.0.

*Transmission electron microscopy*
Identification of EV particles and removal of contaminating extracellular matrix proteins was validated using TEM imaging completed by Harvard Medical School Electron Microscopy Core. 5 µl of the sample was adsorbed for 1 minute to a carbon coated grid (EMS, CF400-CU) that had been made hydrophilic by a 20 second exposure to a glow discharge (25 mA). Excess liquid was removed with a filter paper (Whatman #1), the grid was then floated briefly on a drop of water (to wash away phosphate or salt), blotted again on a filer paper, and then stained with 1% Uranyl Acetate (EMS catalog # 22400) for 20 seconds. After removing the excess stain with a filter paper, the grids were examined in a TecnaiG² Spirit BioTWIN and images were recorded with an AMT NanoSprint43 CCD camera.

**Pathway Analysis (EV-miRNA transcriptomics and EV-proteomics)**

Kyoto Encyclopaedia of Genes and Genomes (KEGG) and Gene Ontology (GO) analysis of targets of differentially expressed EV-miRNAs and EV-proteins (FDR/q-value<0.05) was completed using Enrichr with multiple testing correction using Benjamini-Hochberg procedure.[55,126,127] An adjusted p-value < 0.05 was considered significant. Tabula Sapiens was accessed through the Enrichr database platform and results were filtered to exclude all non-vascular tissues.[72] The TISSUES database was accessed through the STRING database platform (v12.0).[39] Cancer-, and infection- associated pathways were excluded from analysis.

**scRNA-seq data analysis**

*Quality filtering and data pre-processing*
An aggregate filtered feature-barcode matrix containing processed single-cell data from three patients with paired atherosclerotic and marginal region samples was pre-processed using Seurat (Version 5.0.1) for normalization, quality control, batch effect correction, dimensionality reduction, and Louvain clustering. We adopted the quality control criteria from Alsaigh et al,[20] by excluding cells expressing less than 200 or greater than 4000 features, and cells with >10% mitochondrial counts from downstream analysis. We also downsampled to 2850 cells across samples, due to previous concerns around dataset imbalance by Alsaigh et al. Samples were normalized using SCTransform in Seurat, with mitochondrial genes regressed out. Batch effect correction was performed with Harmony. Integration and dimensionality reduction was performed using standard Seurat pre-processing pipelines. Following Louvain clustering, we

annotated T-cells, Monocytes, B-cells, Plasma cells, Mast cells, VSMCs, and ECs using *CD2, AIF1, MS4A1, MZB1, TPSAB1, MYL9,* and *VWF* marker genes described in the original dataset paper, respectively.[20] *CD3E+CD4-CD8-* cells were annotated as double-negative (DN) T-cells, *CD3E+CD4-CD8+* cells were annotated as CD8+ T-cells, *CD3E+CD4-CD8+ITGAE+* cells were annotated as CD8+ tissue resident memory (TRM) T-cells,[128]*CD3E+CD4-CD8-NKG7+* cells were annotated as natural killer (NK) T-cells[129] while a group of *CD2+CD3E-CD4-CD8-NKG7-* cells were annotated as unknown immune cells. Monocyte subtypes were annotated using marker genes *ITGAX* and *SIRPA* for dendritic cells,[130] and marker genes *APOEB3A* for macrophages.[131] VSMC subtypes were annotated using the *MYH11* marker gene for VSMCs,[132] and *LUM* marker gene for fibroblast-like states.[133]

*EV cellular identity enrichment analysis*

We employed the AddModule score function in Seurat (Version 5.0.1) to assess the enrichment of EV protein signatures among different cell types.[134-136] We converted all EV protein cargo into associated genes for a comprehensive or 'global' EV signature enrichment. Plaque- and marginal zone specific signatures were established based on differentially expressed proteins (FDR<0.05, Log2FC>0). Additionally, predicted mRNA targets of differentially expressed EV-miRNA were generated using MicroRNA Enrichment Turned Network (MIENTURNET version 2019-11-25) leveraging the miRTarBase database,[124,125] to analyse EV-miRNA signature enrichment in cellular identities.

EV c*ell communication network inference*

We used the 'CellChat' R package (version 1.6.0) to infer cell interaction networks by filtering the CellChat cell-cell communication ligand database for genes associated with EV protein gene IDs. CellChat curates a signalling molecule interaction database that includes ligand-receptor interactions between soluble and membrane-bound molecules.[71] Predictions of global EV-cell communication was performed by filtering for 1502 genes associated with all EV protein cargo, while plaque- and marginal- zone specific EV-cell communication was performed by filtering for 282 and 379 genes associated with differentially expressed EV protein cargo (FDR<0.05), respectively. Heatmap diagrams visualizing incoming signalling patterns across different cell types were generated using the 'netAnalysis_signalingRole_heatmap' function. Chord diagrams were generated using the 'netVisual_chord_gene' function with modifications to visualize high probability ligand-receptor pairs through the 'reduce' parameter (and re-label ligand identities as 'EV'). Comparative analysis of predicted EV-cell communication between cells from plaque- and marginal-zone was performed with CellChat, with visualizations of overall information flow across different signaling patterns using the 'rankNet' function with default parameters.

**Overrepresentation and network analysis**

Comparing marginal and plaque zone tissue EV loads with an FDR cut-off of q<0.05, we identified differentially expressed proteins and miRNA for symptomatic and asymptomatic cohorts. Proteins were directly mapped to UniProtKB gene lists and miRNAs were mapped to gene lists from predicted mRNA target enrichment using MIENTURNET.[124,125]

The network based approach incorporated protein-protein from STRING, BioGRID, MINT, and IntAct, and protein-miRNA interaction data from RNAInter. Protein-protein interaction data was curated to include interactions with >30 experimental citations. Similarly, protein-miRNA interaction data was curated to include interactions with a confidence score >0.4. The curated data was then integrated into a multiomic network using BIONIC, a deep-learning based biological network integration algorithm.[40] BIONIC parameters were fine-tuned to ensure a stable reconstruction loss across 3000 epochs. The final network comprised 512 embedded features and was integrated using a learning rate of 5 x $10^{-6}$, with a GAT-head dimensionality of 196. The multiomic network was then clustered using the Louvain algorithm across values of ranging from 1 to 5 to form a set of modules. Modules enriched with differentially expressed proteins/miRNA were identified using Fisher's exact ($q<0.05$). Proteins and miRNA from the enriched modules were mapped to genes in the same way as above.

Gene lists were used in overrepresentation analyses to identify enriched GO biological processes and/or KEGG pathways. The overrepresentation analysis employed Fisher's exact test, with an FDR cut-off of $q<0.05$. In the case of the network-based approach, pathway enrichment was done on a per-module basis with each differentially expressed proteins/miRNA-enriched module representing a distinct gene list.

Code used in the scRNA sequencing and overrepresentation analysis is deposited at Github (https://github.com/npunw/carotid_plaque).

#### Cell Culture

Telomerase-Immortalized Human Umbilical Vein Endothelial Cells (TeloHUVECs; passage 16-24) were cultured in ECM (SC1001) containing 5% Fetal Bovine Serum (FBS), 1X endothelial growth supplement (SC1052), and 1X penicillin-streptomycin solution (SC0503). Cells were grown on polystyrene tissue culture plates coated with Type 1 Collagen (150 µg/mL, Sigma-Aldrich C7661) and seeded at a density of 5 x $10^3$ cells/cm$^2$. Cell viability was assessed by Trypan Blue exclusion test with 0.4% trypan blue (GibcoTM 15250061) mixed 1:1 with cells in suspension.

#### *In Vitro* Sprouting Angiogenesis Assay

TeloHUVECs were suspended in 0.2% methylcellulose (Sigma, M0512) mixture prepared in EBM$^{TM}$-2 Basal Medium (Lonza, CC-3156) at a concentration of 4 x $10^4$ cells/ml. Twenty µl cell suspension drops were pipetted on a 10cm plastic plate. Plates were flipped upside-down to form hanging droplets and incubated for 24 hours at 37°C to allow the formation of spheroids. Spheroids were collected by gently washing with a 10% FBS/PBS solution followed by centrifugation at 300 x g for 5 minutes. A collagen matrix was prepared on ice using 0.6% methylcellulose with 40% FBS, Type 1 collagen (Cultrex, Cat. 4447-02001), $NaHCO_3$ (15.6 mg/ml), and NaOH (1 M). The collected spheroids were resuspended in the collagen matrix and transferred into a 96-well plate. For polymerization, the matrix was incubated for 10 minutes at 37°C, followed by the addition of 50µl of complete ECM media. Spheroids were treated for 24 hours in a 37°C incubator with VEGF (50 ng/ml; Positive control), marginal zone eluate, plaque eluate, pooled (two patients) symptomatic marginal zone and symptomatic plaque zone EVs (2 x $10^{10}$ to 4 x $10^{10}$ sEV; 24 h), and untreated control. After 24 hours, spheroids were fixed with 4 % PFA (Fisherscientific, 15710) for 30 minutes and permeabilized with 0.5 % Triton (Sigma, SLBZ4187) for 30 minutes, followed by a wash with PBS. Spheroids were stained with phalloidin 488 (1:40 dilution, ThermoFisher Scientific, lot 2219253) and Hoechst33342 (1:500, Thermo Fisher Scientific, H3570) overnight. Spheroids were imaged using the NIKON AR1 confocal microscopy using 10X magnification. In a blinded fashion, the number of sprouts per spheroid and the sprout length per spheroid were manually quantified using Fiji.

**Bulk proteomics from carotid plaque and adjacent regions in BiKE**

Full methods have previously been published.[37] Briefly, CEA tissue was obtained from 18 patients (9 asymptomatic and 9 symptomatic) with a central plaque portion and adjacent peripheral ends of the sample for comparison used in proteomic analysis by liquid chromatography mass spectrometry/mass spectrometry (LC-MS/MS). Samples were crushed while frozen, lysed and sonicated in buffered solution, and then underwent centrifugation with supernatants collected for digestion. Resulting peptide mixtures were labelled with isobaric Tandem mass tags. Sample clean-up was performed with solid-phase extraction and pools were pre-fractionated by high resolution isoelectric focusing before LC-MS/MS analysis. Raw MS/MS files were converted to mzML format using msconvert from ProteoWizard tool suite. Spectra were searched using MSGF+ (v10072) and Percolator (v2.08) where 8 subsequent search results were grouped for Percolator target/decory analysis. Reference database used was the human protein subset of ENSEMBL 75. Peptide spectrum match (PSM) found at 1% PSM- and peptide-level false discovery rate (FDR) were used to infer gene identities, and median normalization of ratios on the PSM was performed. Protein level FDRs were calculated using the picked-FDR method.

**Statistical Analysis**

Statistical analyses were performed using Graphpad Prism 10.2.0 for Mac (GraphPad Sofware, Inc., La Jolla, CA, USA). All bar graphs were composed of 3-20 independent samples and data points are presented as the mean ± standard error of mean (SEM). Volcano plots display median and interquartile range. The Fisher's exact test tested the prevalence of GO terms in whole tissue proteomics. For $n<10$, a two-sided Mann-Whitney or Wilcoxon (for paired samples) test was completed with the assumption of a non-normal distribution. For $n \geq 10$, normality was tested using the Shapiro-Wilk test and for normally distributed data, differences between two groups were tested using a two-tailed unpaired Student's *t*-test. For comparisons between three or more groups, a one-way ANOVA was used with Šidák multiple comparisons test . Differences were significant for values of $p<0.05$ and are indicated in the figures. As stated above, the false discovery rate for multiple comparisons of the transcriptomics and proteomics data was controlled with the Benjamini–Hochberg procedure. Comparison of fold enrichments between cohorts regarding endothelial-related GO biological processes was performed using ANOVA and Tukey's honestly significant difference.

***Data Availability***

miRNA-seq data is deposited at the Gene Expression Omnibus (GSE270496, Reviewer Access olknkoamdrcldgz). The mass spectrometry proteomics data are deposited to the ProteomeXchange Consortium via the PRIDE partner repository (Project accession: PXD052938, Username:, Password: 7PEUstaq5l4v). Datasets are private for reviewers and will be made public after acceptance. The carotid scRNA-sequencing data and materials from this study are publicly accessible in the GEO database under accession code GSE159677. [20]

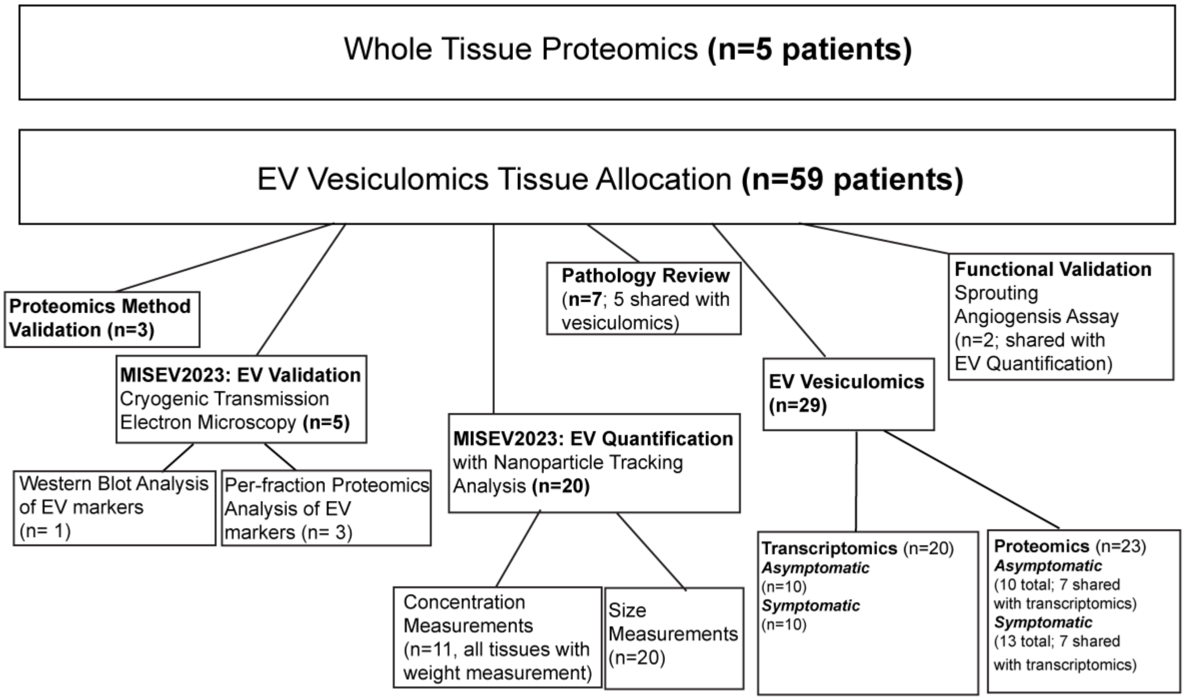


**Methods Figure 1. Human Carotid Endarterectomy Sample Allocation.** A total of 64 carotid endarterectomy samples were used in the study. Five paired plaque and marginal zones were used to complete whole tissue proteomics. The remaining 59 patient samples were used for proteomics methods validation (1 donor contributing plaque and marginal zones), MISEV2023 based EV validation and quantification, histopathology, vesiculomics, and functional validation. A portion of samples were utilized for more than one experiment. Two samples used in EV quantification were also used in functional validation studies. Five samples that underwent EV vesiculomics were also sectioned and stained for pathology review. Majority of samples (14 patient samples) used in vesiculomics underwent both transcriptomics and proteomics (tissue was divided in half), however there was an addition of nine samples for EV-proteomics (3 asymptomatic, 6 symptomatic).

**Methods Table 1. Available patient demographics**

| | **Symptomatic (n=12)** | **Asymptomatic (n=5)** |
|---|---|---|
| Sex (Male) | 12 (100%) | 5 (100%) |
| Age | 77.5± 8.91 | 73.4 ± 2.70 |
| Side of Carotid Surgery (Right) | 6 (50%) | 3 (60%) |
| Smoking Status | | |
| Present | 1 (8%) | 1 (20%) |
| Previous | 5 (42%) | 2 (40%) |
| COPD | 0 (0%) | 3 (60%) |
| Hypertension | 10 (83%) | 3 (60%) |
| Dyslipidemia | 7 (58%) | 1 (20%) |
| Diabetes Mellitus | 3 (25%) | 3 (60%) |
| Coronary Artery Disease | 8 (67%) | 3 (60%) |
| Chronic Kidney Disease | 3 (25%) | 1 (20%) |
| Antiplatelets | 11 (92%) | 1 (20%) |
| Anticoagulants | 6 (50%) | 4 (80%) |
| Statins | 12 (100%) | 5 (100%) |
| Beta-Blockers | 9 (75%) | 5 (100%) |
| Anti-hypertensives | 11 (92%) | 5 (100%) |
| Highest PSV (cm/s) | 262.14 ± 175.61 | 363.67 ± 17.50 |
| ICA:CCA Ratio | 3.66 ± 3.15 | 5 ± 0.62 |

**Supplemental Figure 1**

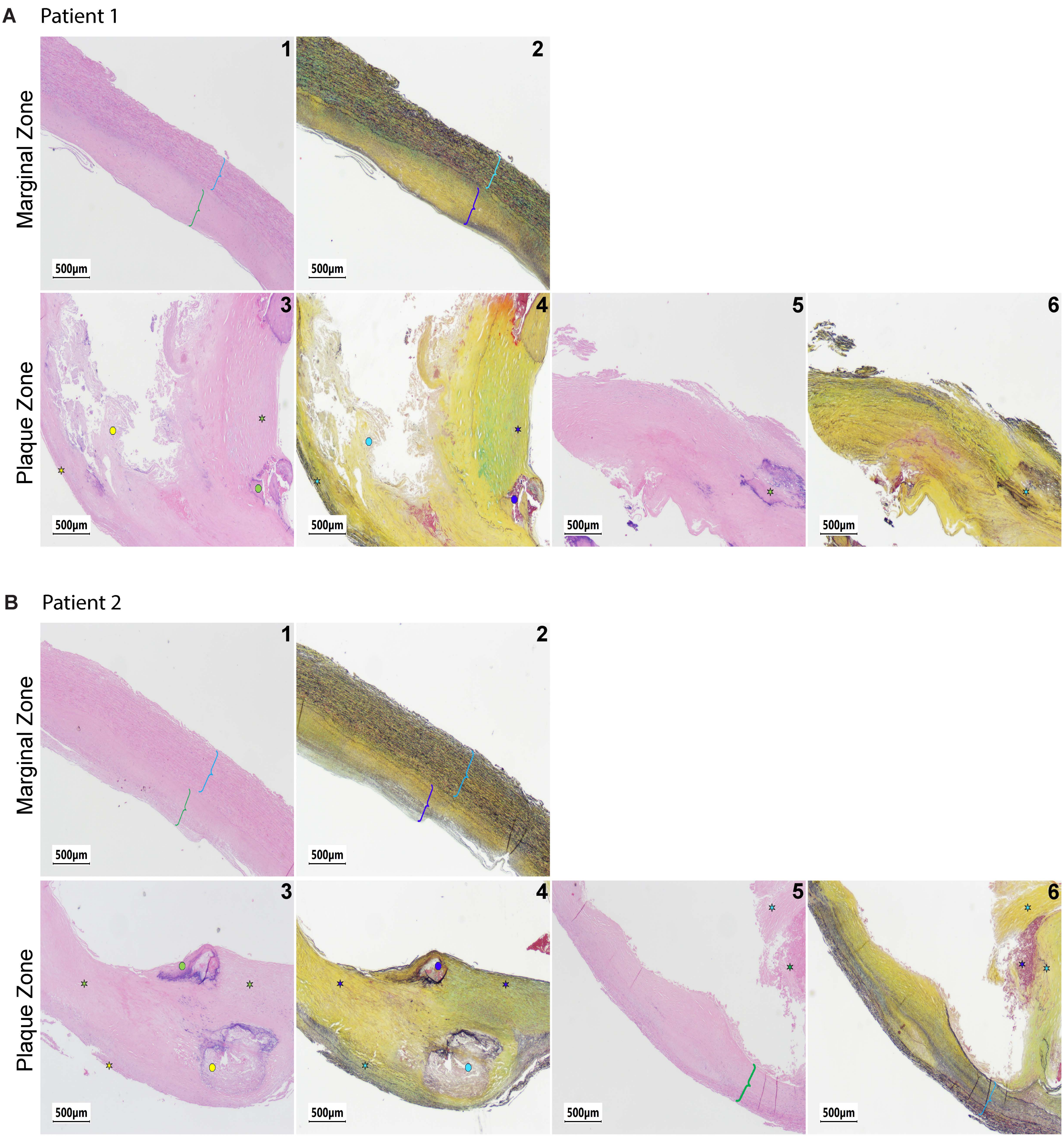

#### Supplemental Figure 2

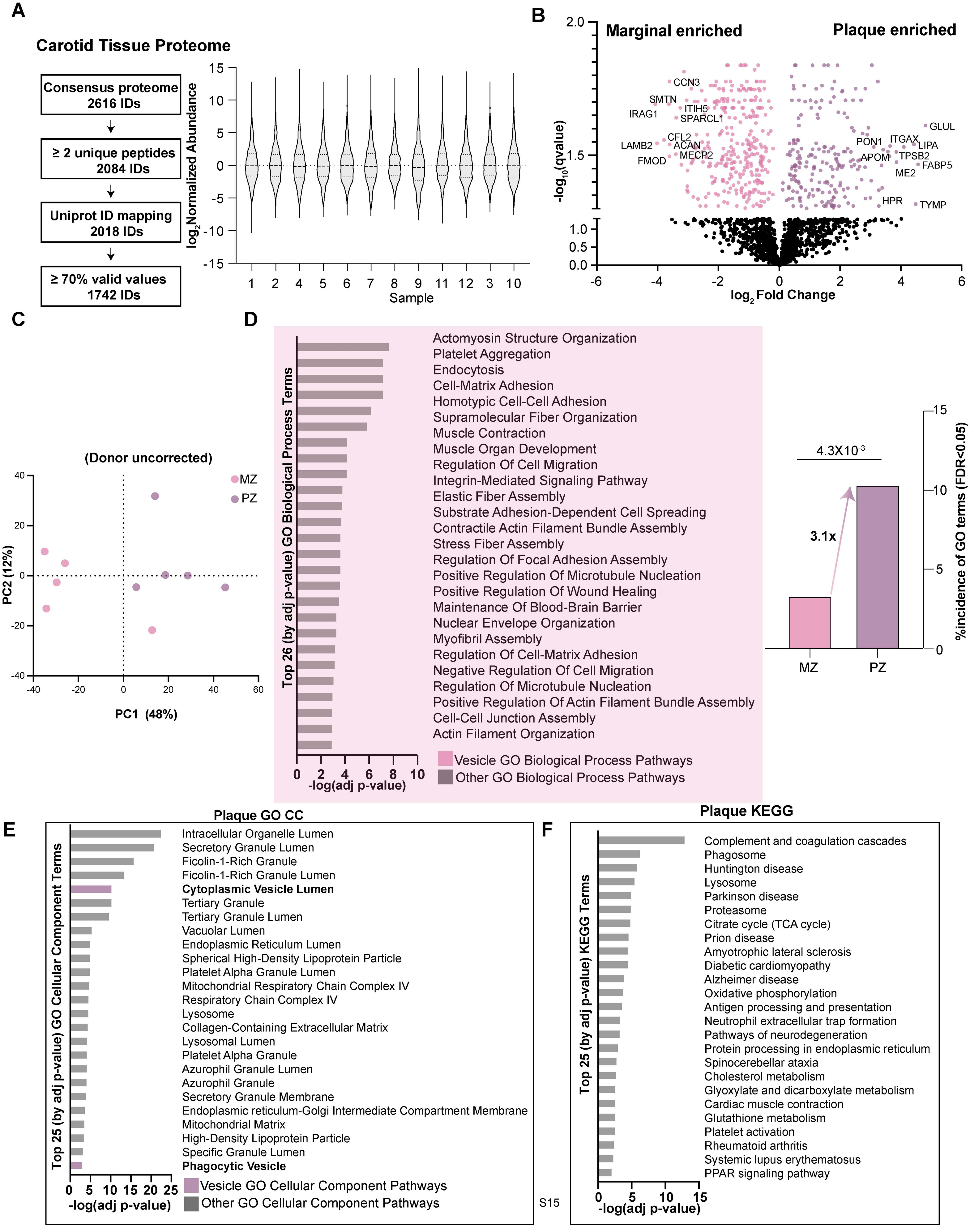

#### Supplemental Figure 3

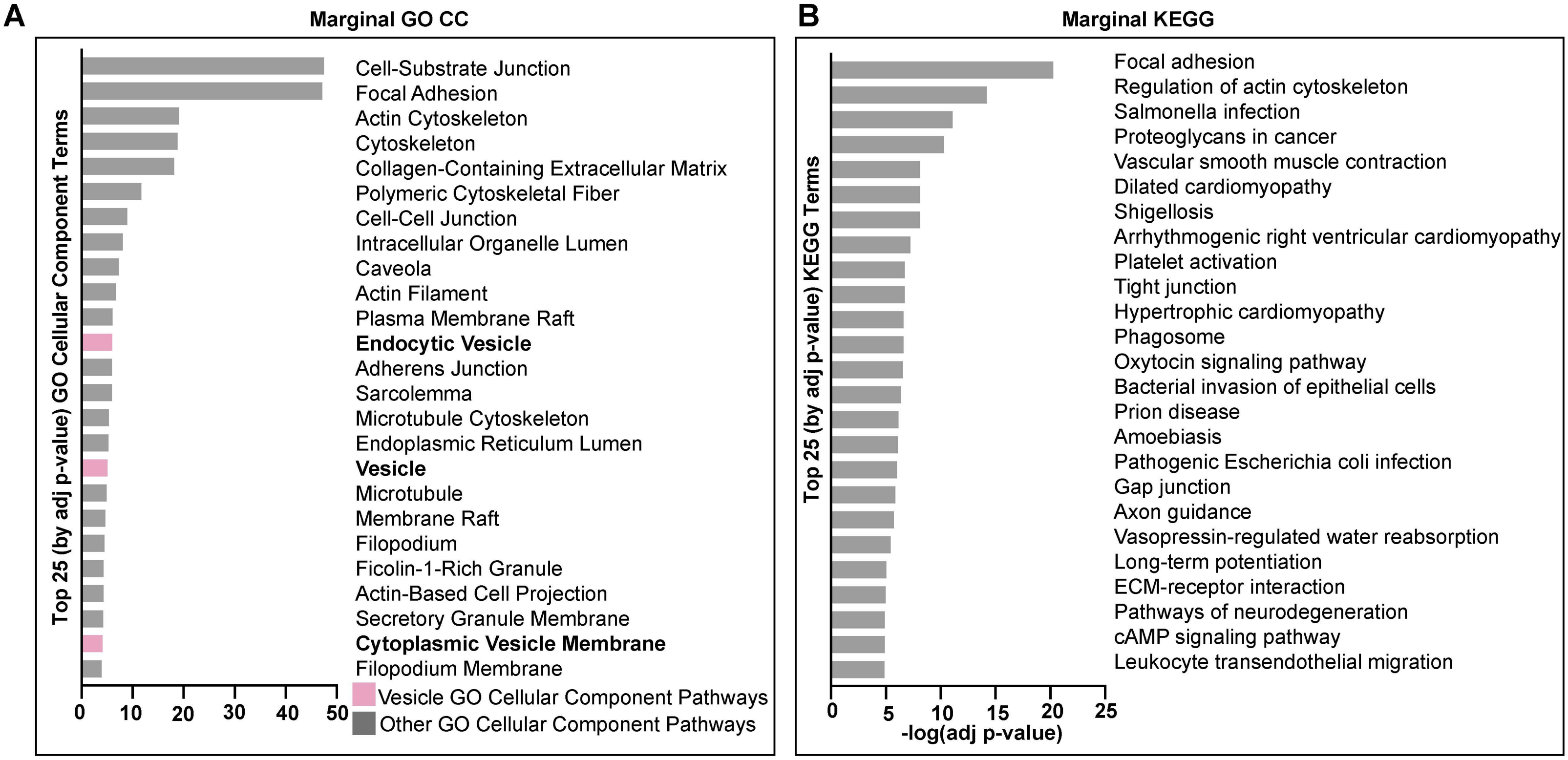

### Supplemental Figure 4

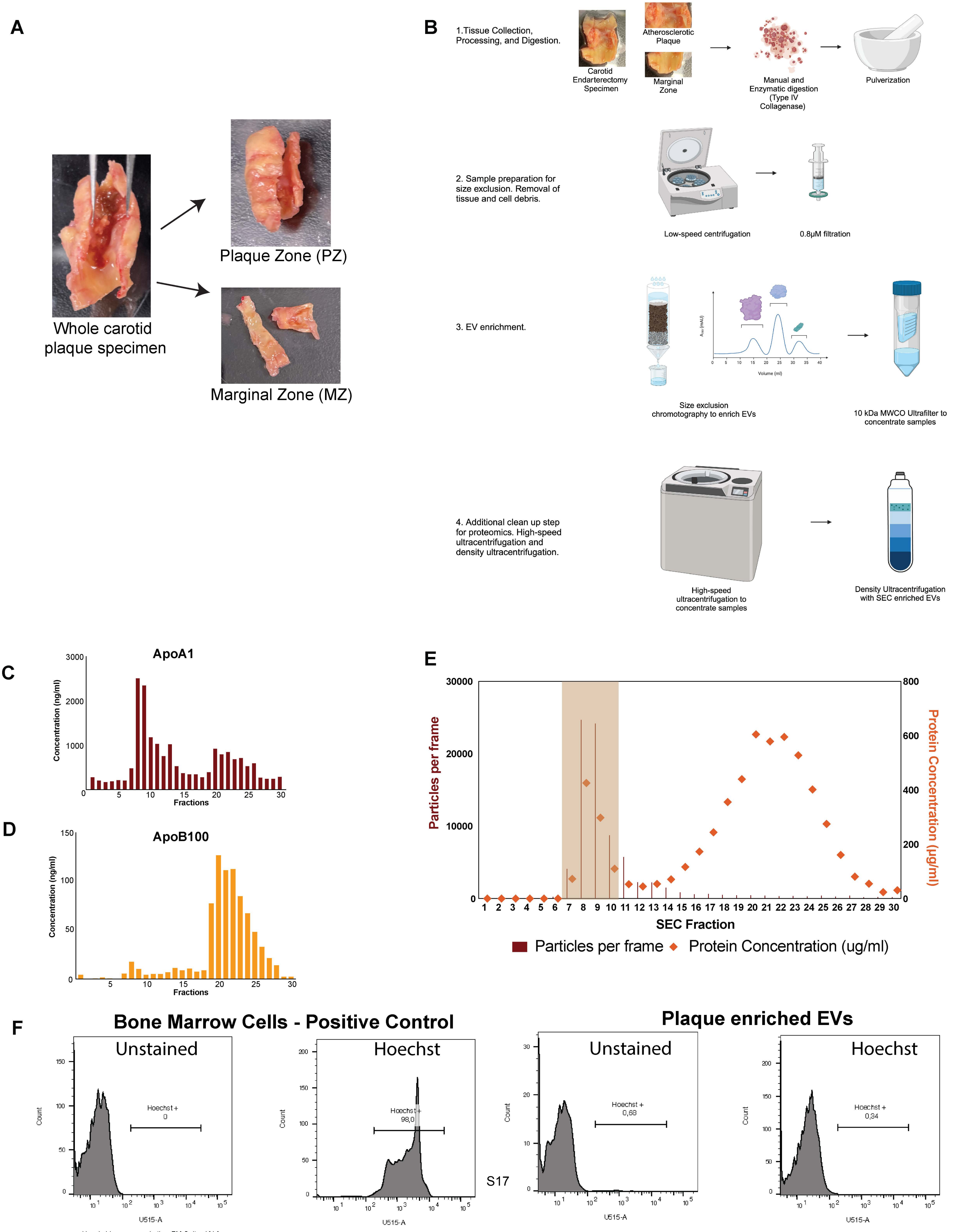

#### Supplemental Figure 5

A

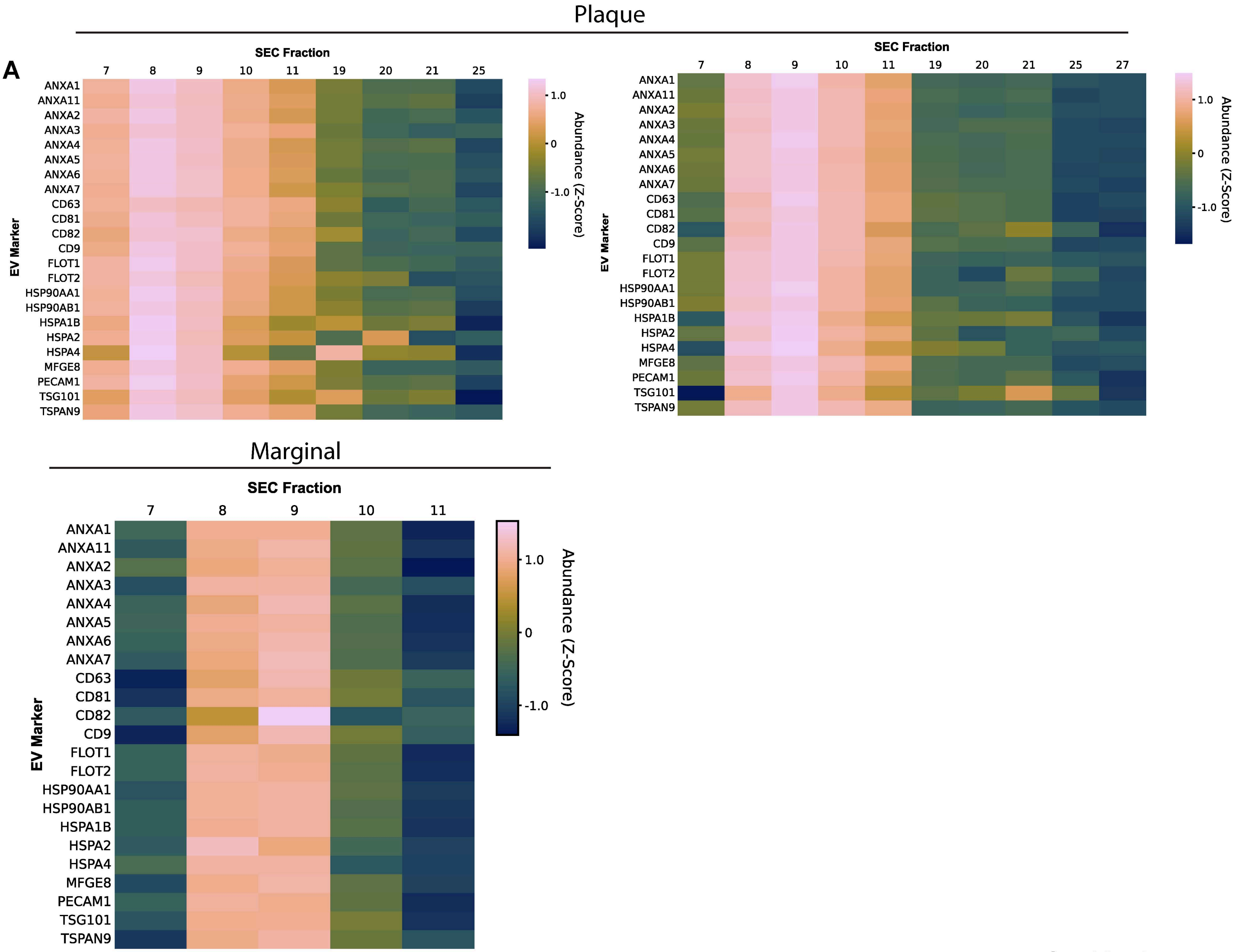


B

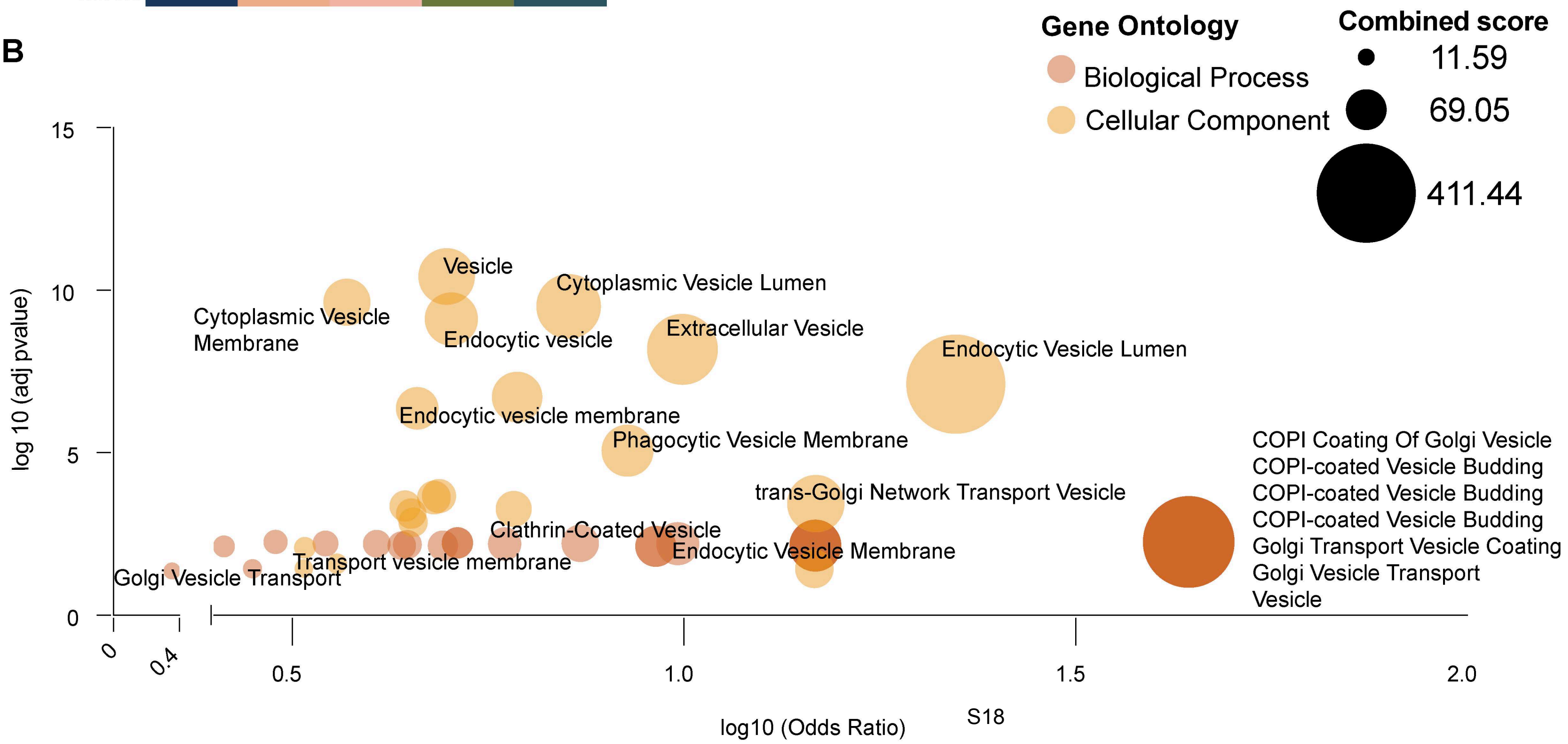

**Supplemental Figure 6**

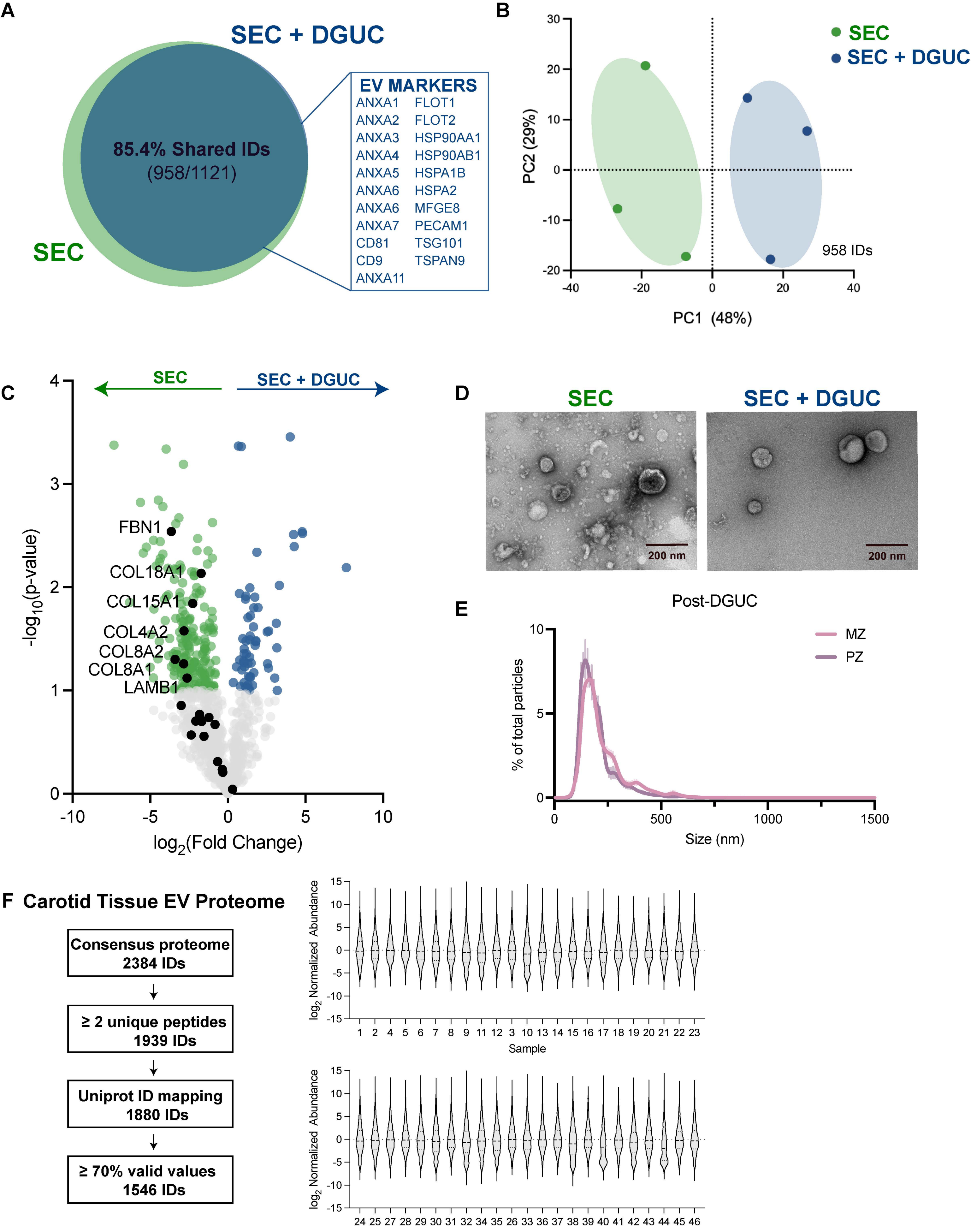

#### Supplemental Figure 7

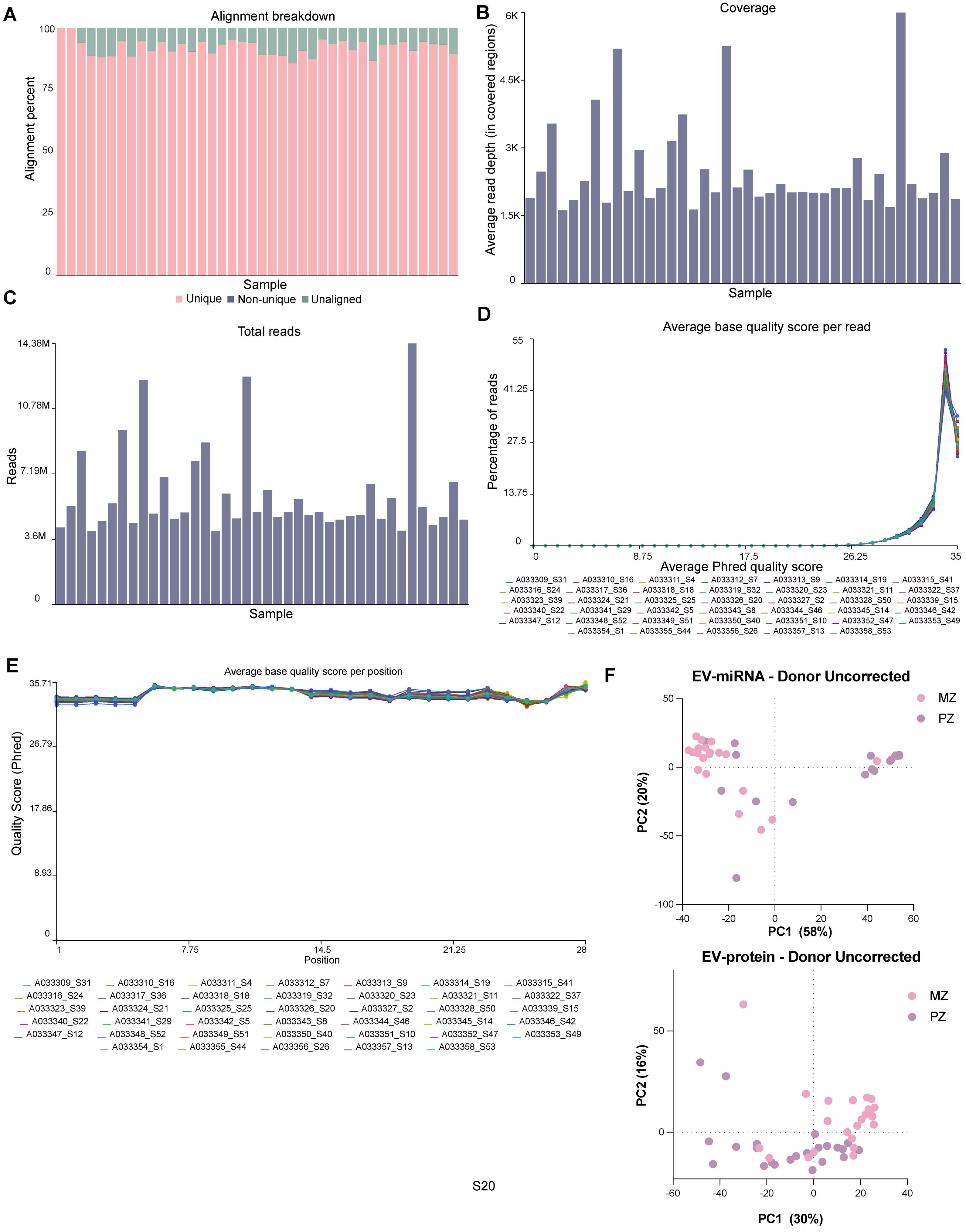

#### Supplemental Figure 8

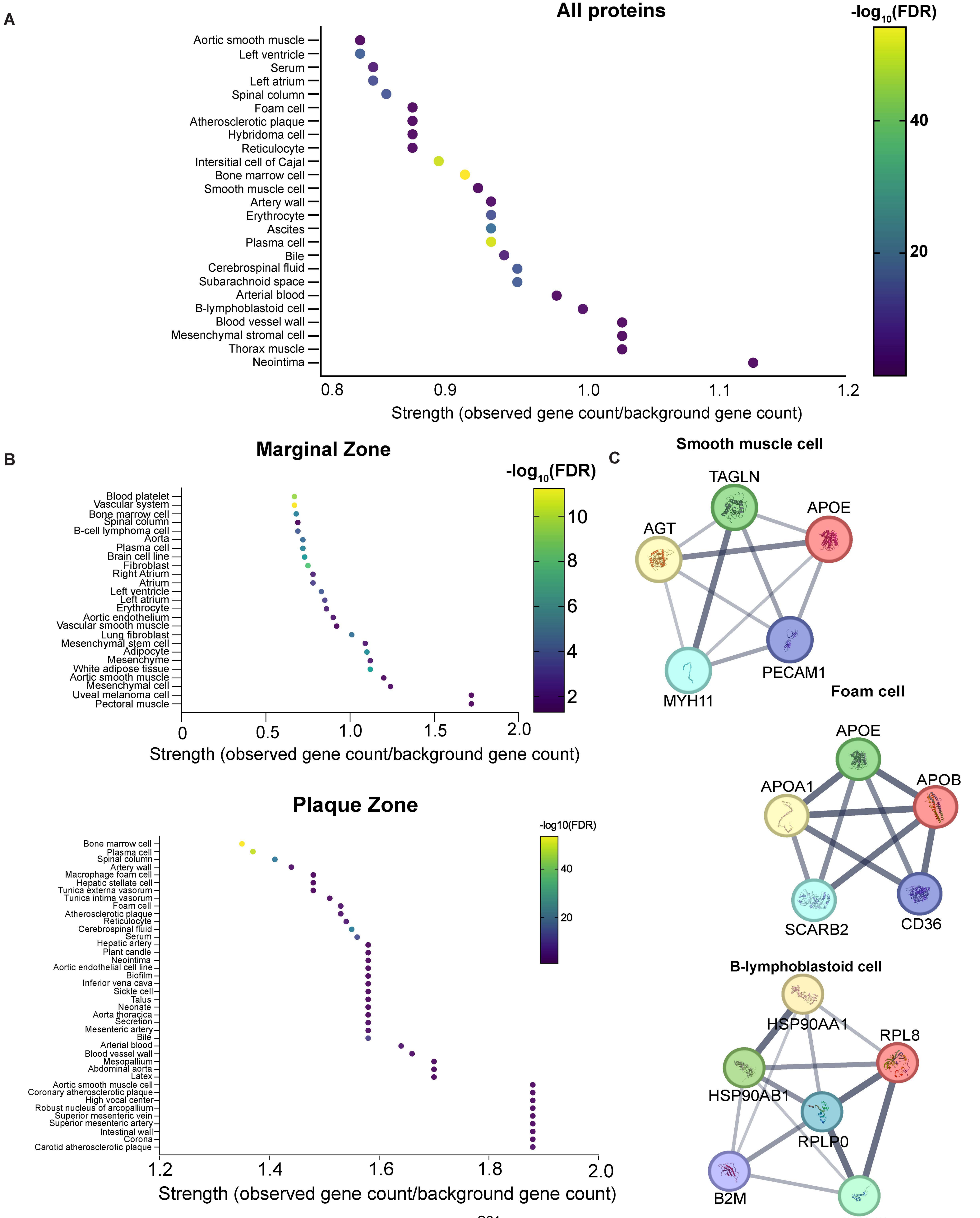

### Supplemental Figure 9

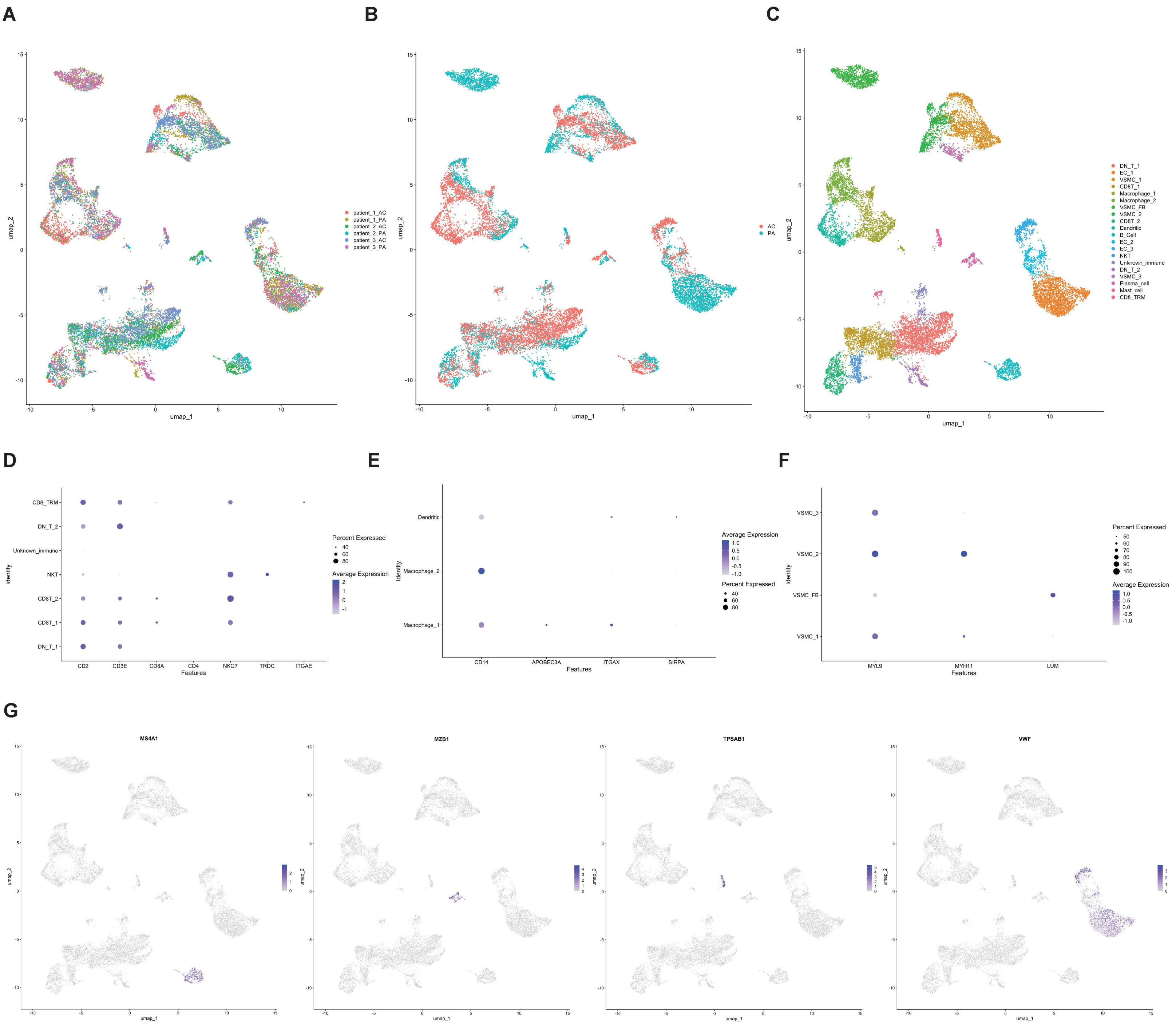

#### Supplemental Figure 10

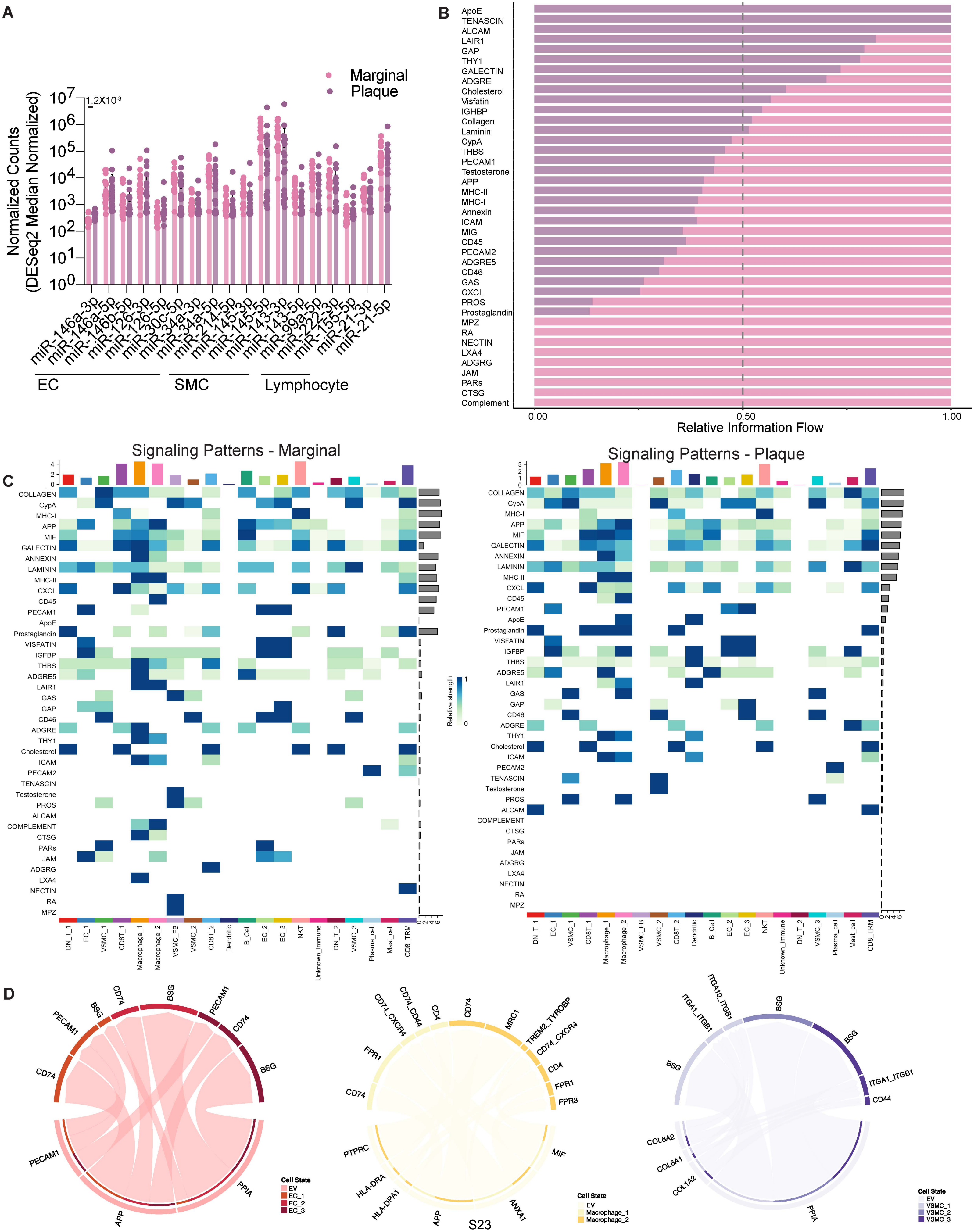

### Supplemental Figure 11

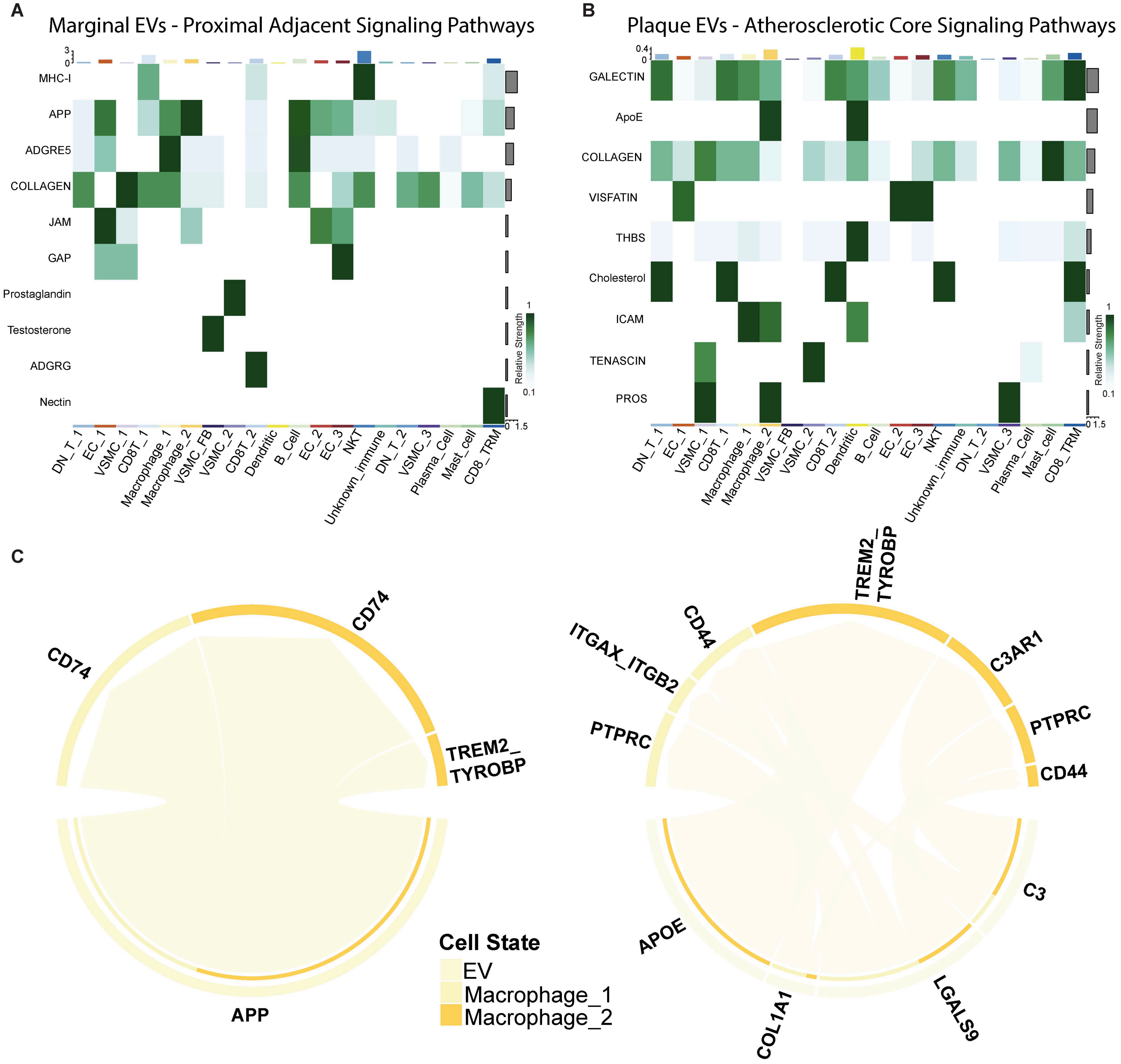

#### Supplemental Figure 12

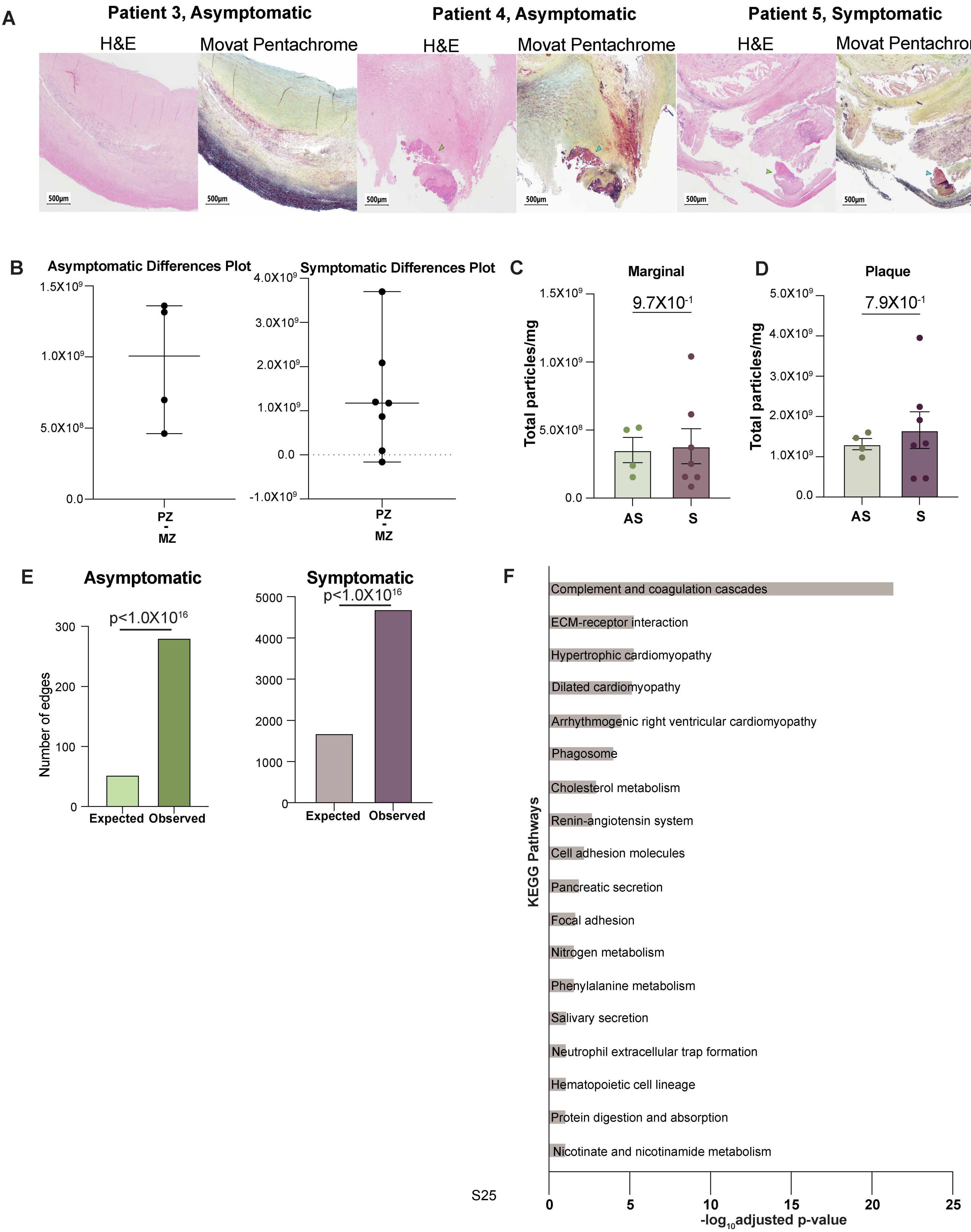

**Supplemental Figure 13**

A

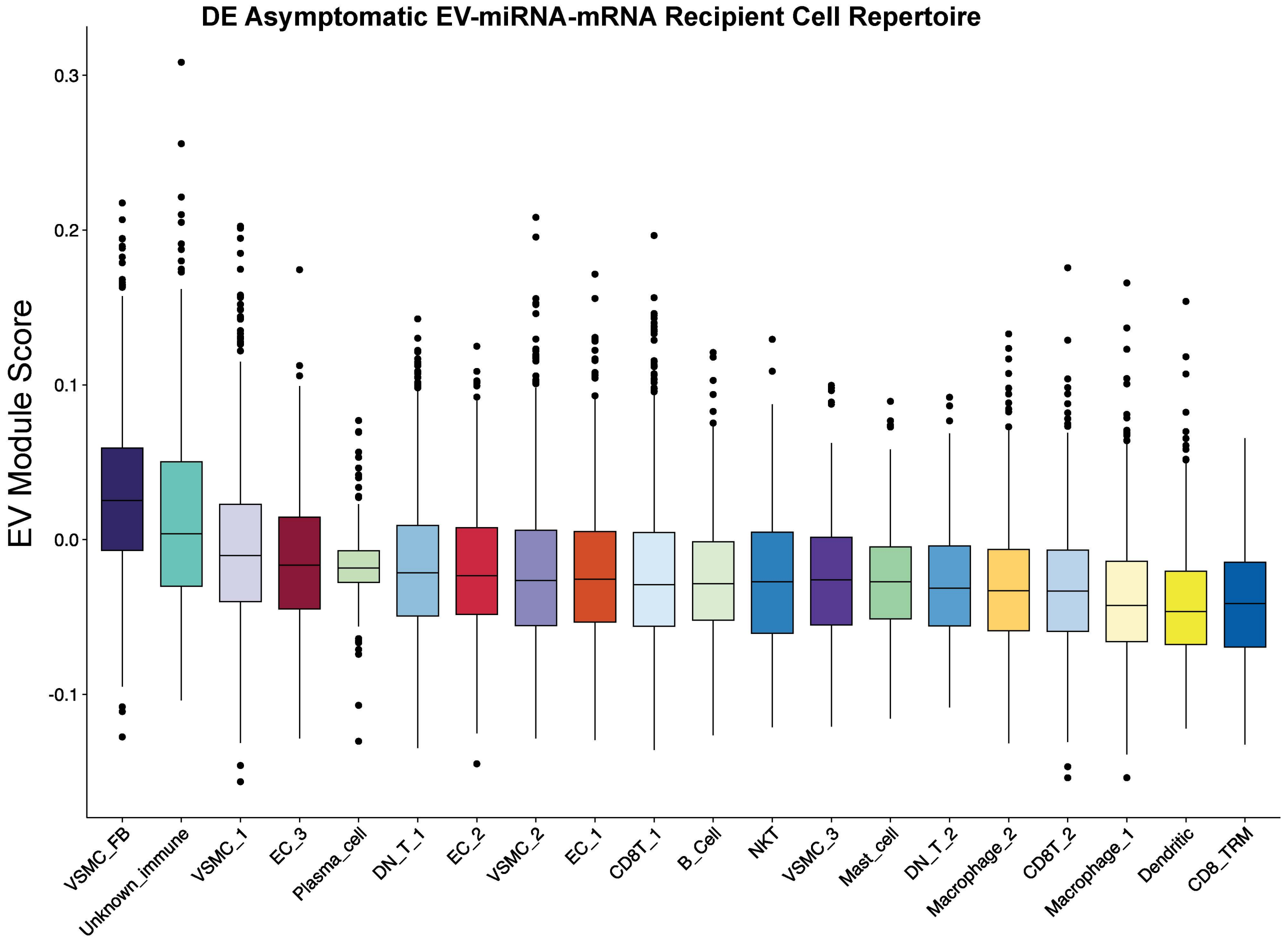


B

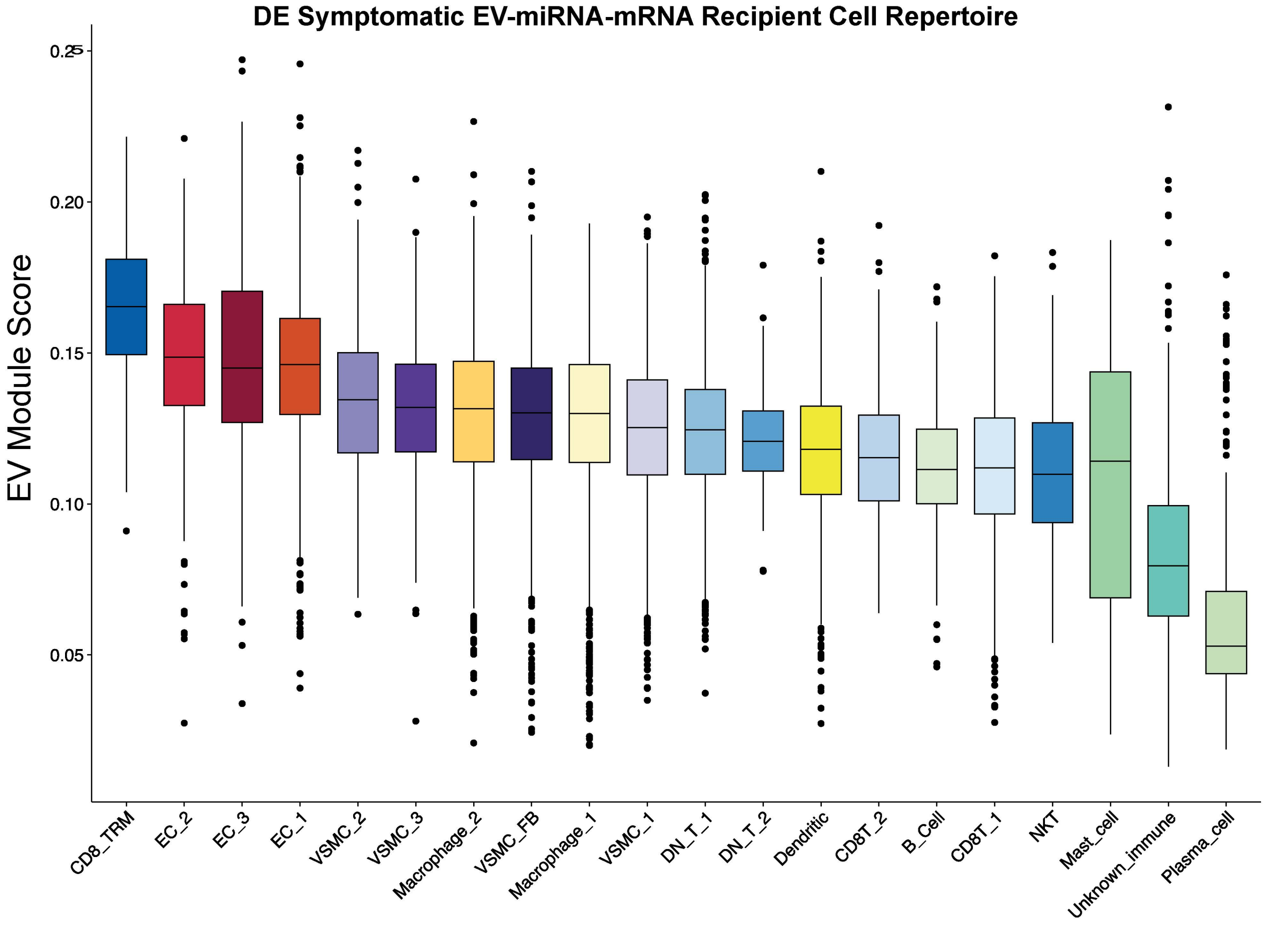

#### Supplemental Figure 14

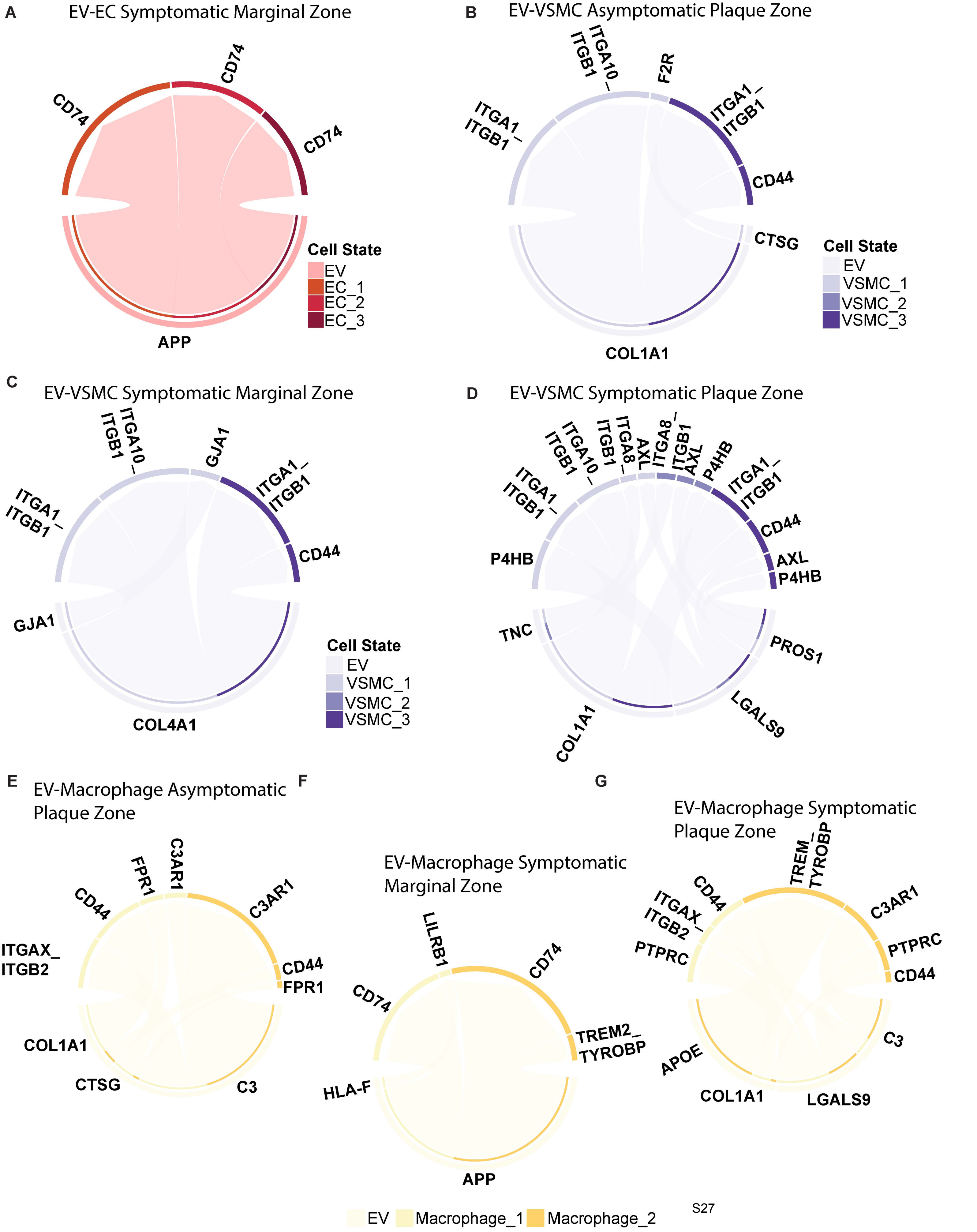

#### Supplemental Figure 15

EV marker CD63 in atherosclerotic plaques

Protein values normalized to CD63 levels

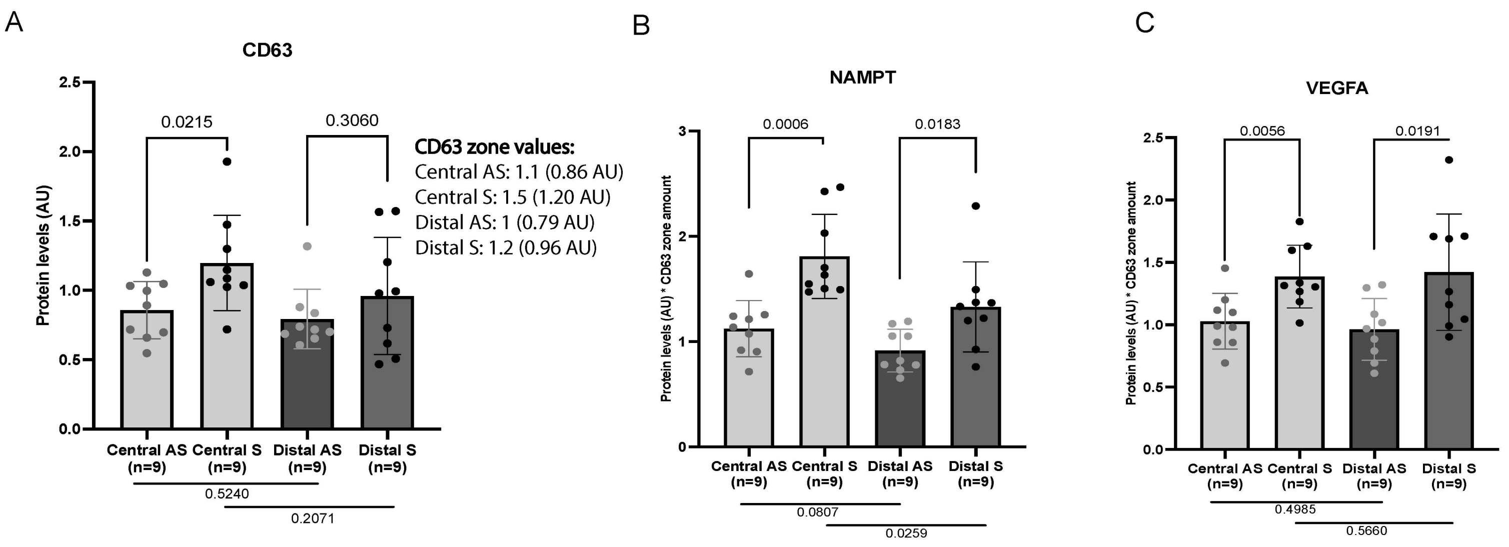

#### Supplemental Figure 16

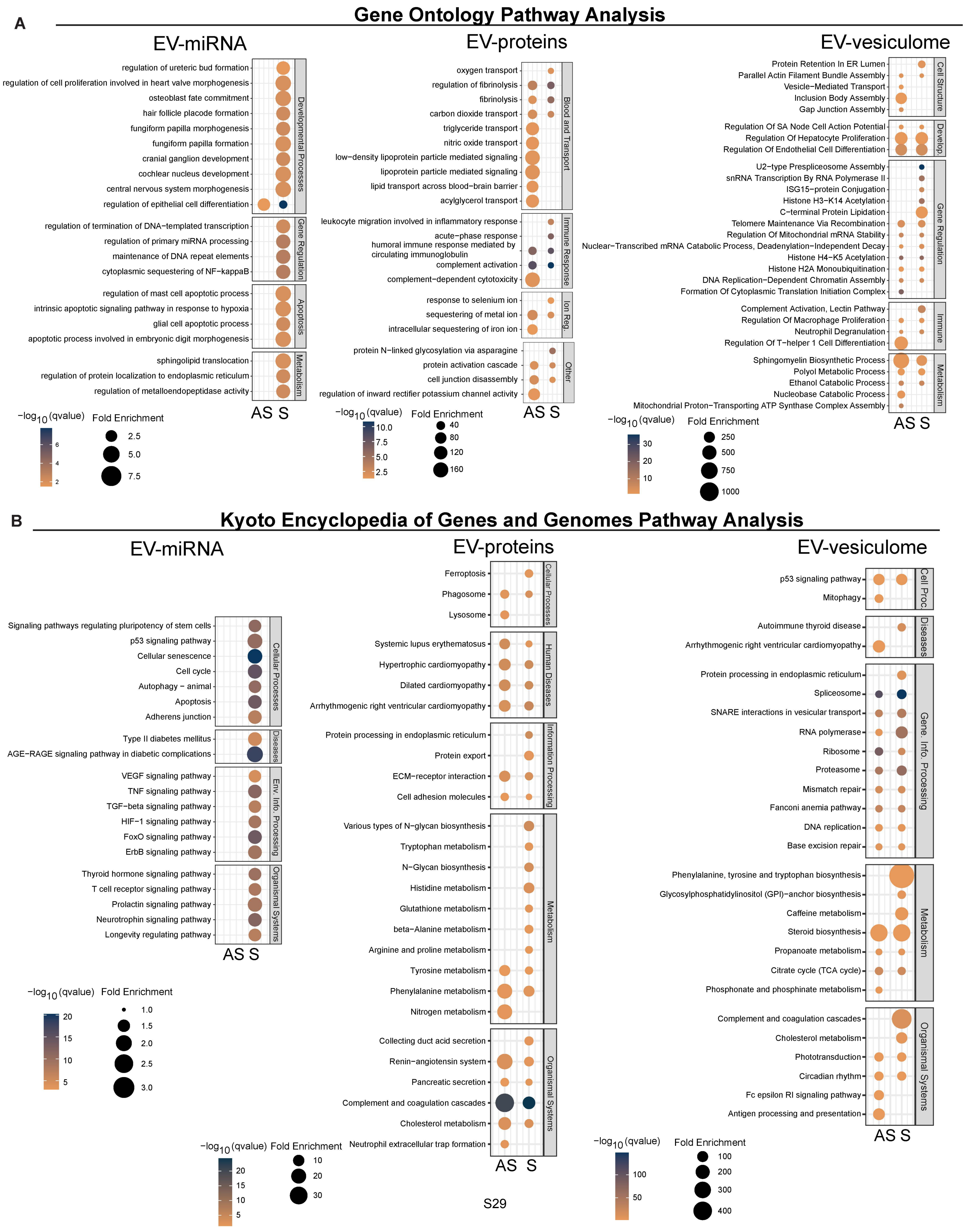

### Supplemental Figure 17

#### Gene Ontology Overrepresentation Analysis

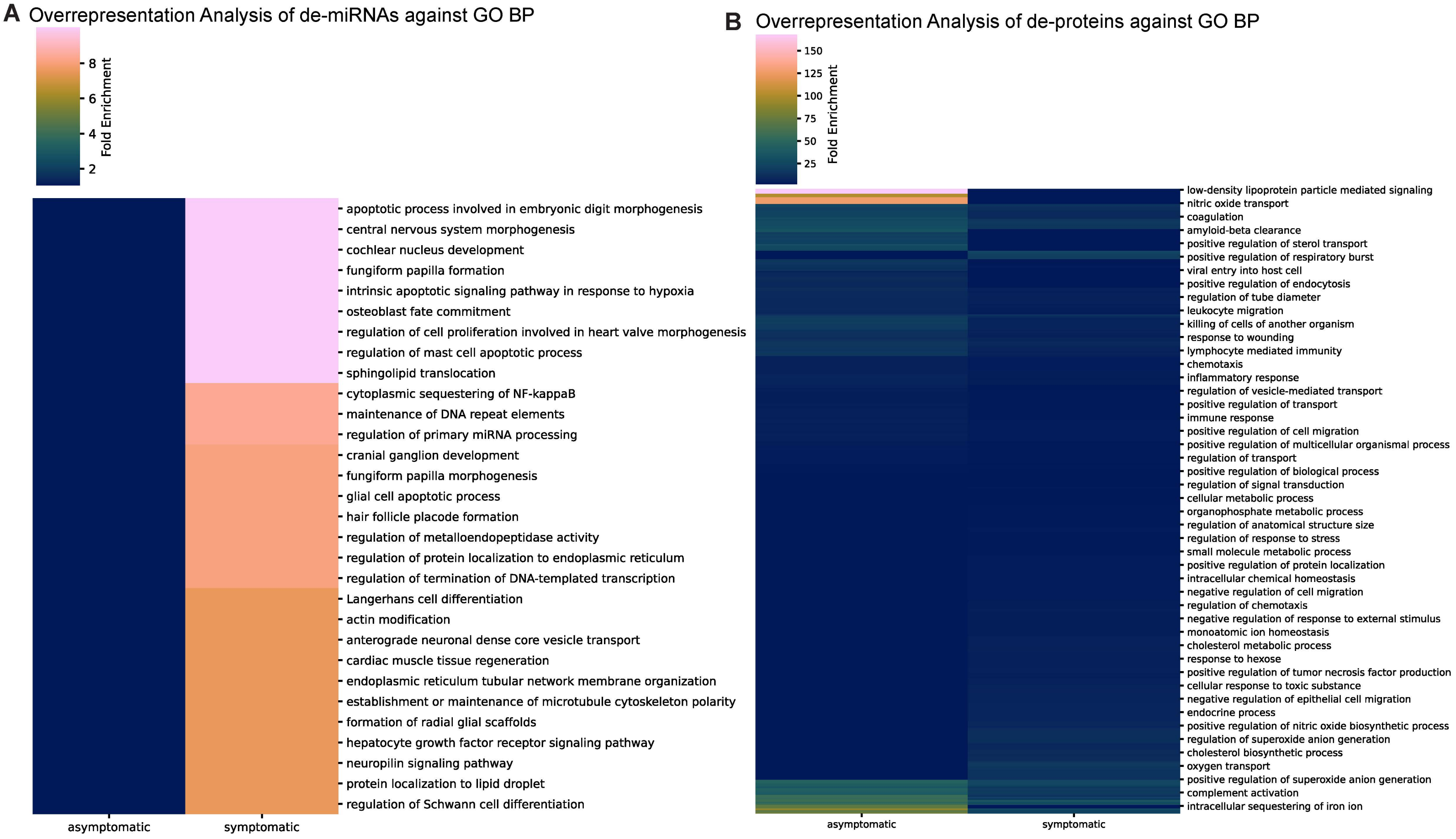


#### Kyoto Encyclopedia of Genes and Genomes Overrepresentation Analysis

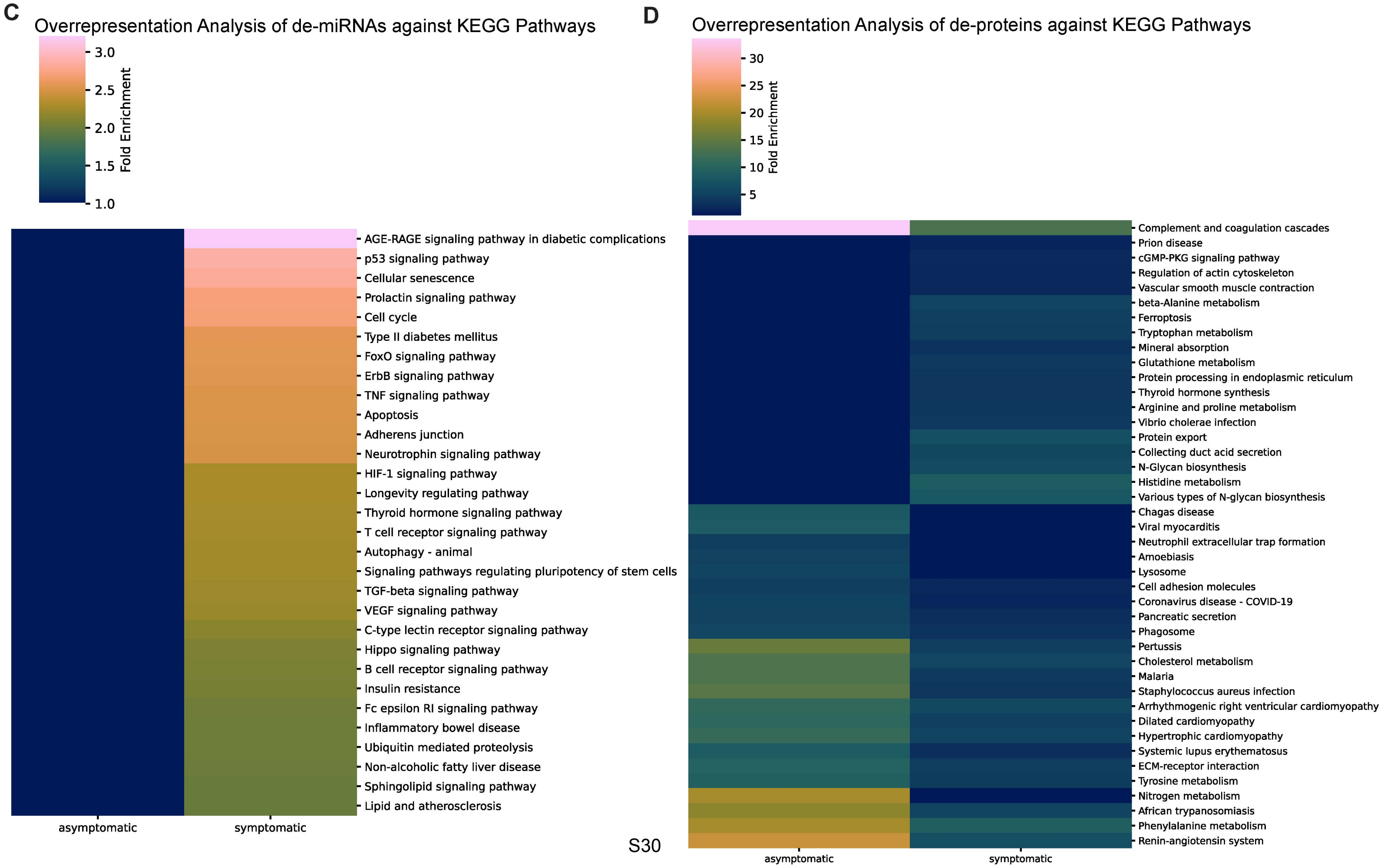

#### Supplemental Figure 18

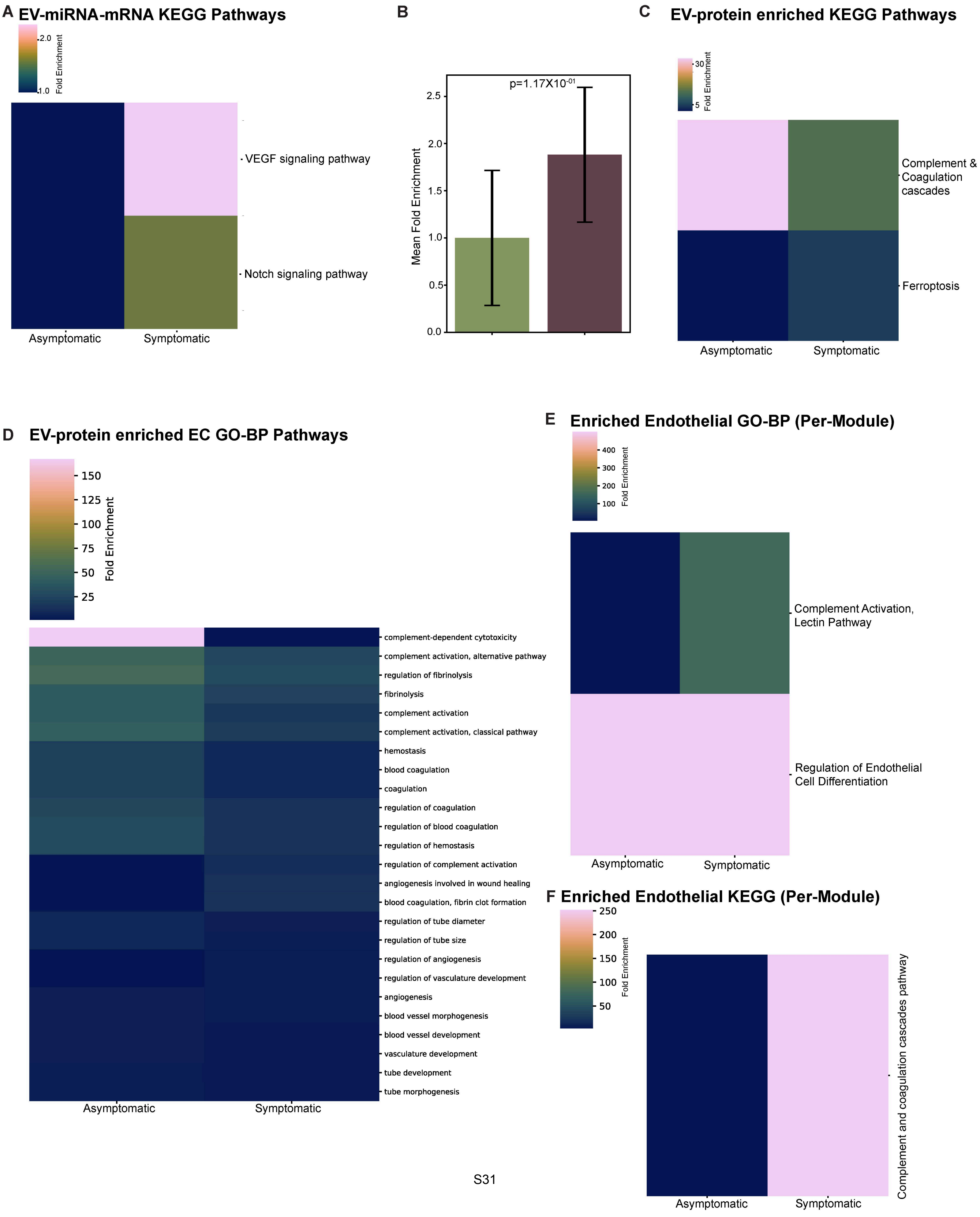

#### Supplemental Figure 19

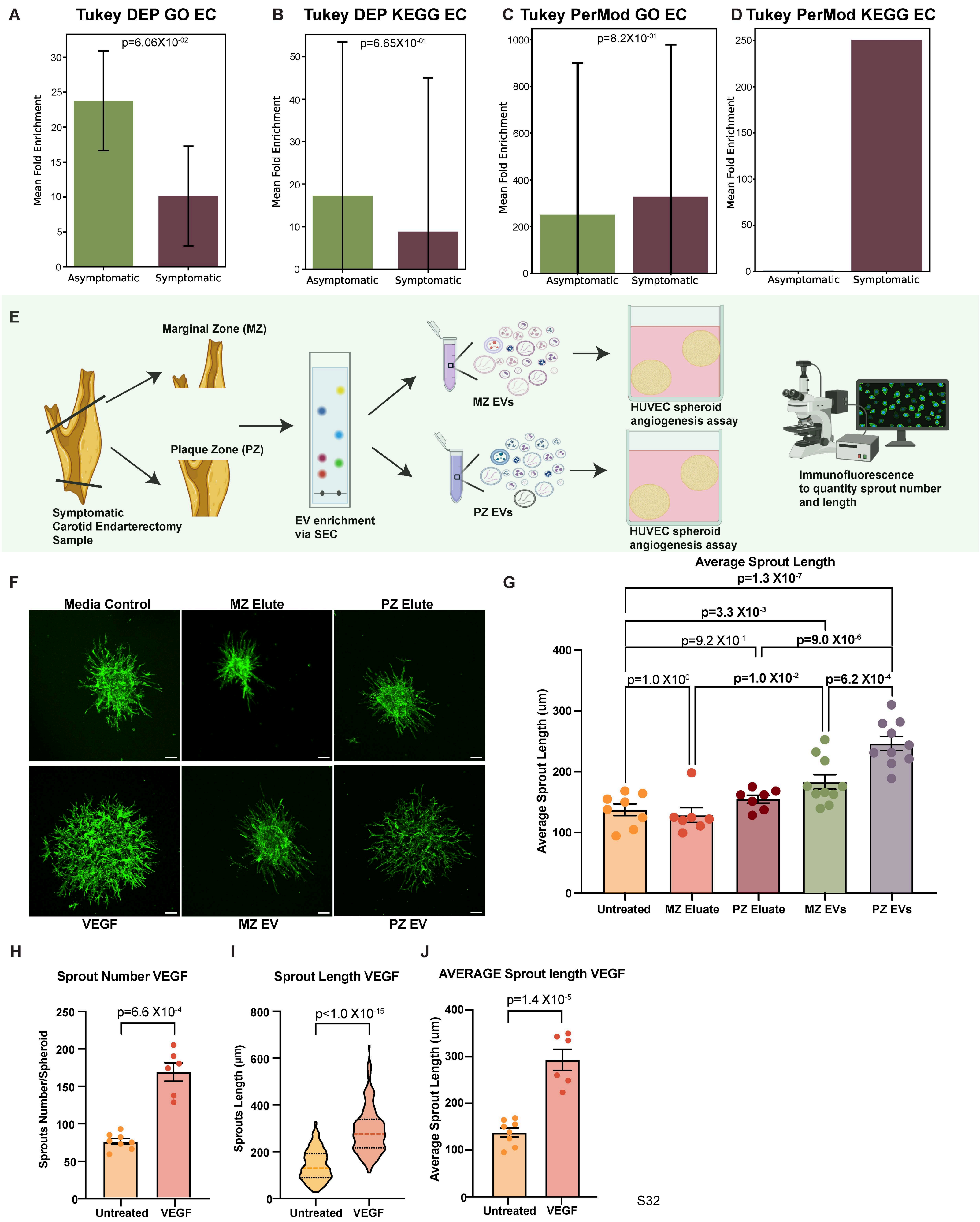

#### Supplemental Figure Legends

**Supplemental Figure 1. Histopathology validates tissue dissection (plaque and marginal zones) and identifies characteristic features of late-stage atherosclerosis predominantly in plaque zones. A.** Hematoxylin and eosin (H&E) and Movat pentachrome stain of plaque and marginal zones from patient one. Area adjacent to plaque (marginal zone; 1-2). H&E staining: intimal hyperplasia (green bracket), media with moderate degenerative changes (cyan bracket; 1). Movat pentachrome staining: intimal hyperplasia (blue bracket), media with moderate degenerative changes (cyan bracket; 2). Middle zone of large, calcified plaque (3-4). H&E staining: surface calcification (green octagon), fibrous cap (green star), and necrotic center (yellow octagon); media (yellow star; 3). Movat pentachrome staining: surface calcification (blue octagon), fibrous cap (blue star), necrotic center (cyan octagon), media (cyan star; 4). Edge zone of large, calcified plaque (5-6). H&E staining: calcification (green star; 5). Movat pentachrome staining: calcification (cyan star; 6). **B.** H&E and Movat pentachrome stain of plaque and marginal zones from patient two. Area adjacent to plaque from (marginal zone; 1-2) H&E staining: intimal hyperplasia (green bracket), media with moderate degenerative changes (cyan bracket;1). Movat pentachrome staining: intimal hyperplasia (blue bracket), media with moderate degenerative changes (cyan bracket; 2). Middle zone of large, calcified plaque (3-4). H&E staining: surface calcification (green octagon), fibrous cap (green stars), and necrotic center (yellow octagon); media (yellow star; 3). Movat pentachrome staining: surface calcification (blue octagon), fibrous cap (blue star), necrotic center (cyan octagon), media (cyan star, 4). Edge zone of large, calcified plaque (5-6). H&E staining: media with marked disruption of the subintimal layer by plaque (green bracket), plaque (artifactually displaced from the rest of the tissue) with fibrous cap (cyan star) and intraplaque hemorrhage (green star, 5). Movat pentachrome staining: media with marked disruption of the subintimal layer by plaque (cyan bracket), plaque (artifactually displaced from the rest of the tissue) with fibrous cap (cyan star) and intraplaque hemorrhage (blue star; 6). Scale bar (500 μm) as shown.

**Supplemental Figure 2. Carotid tissue proteomics delineates differential protein enrichment in plaque versus marginal zones with pathway analysis identifying atherogenic signatures. A.** Proteomic data pre-processing and per-sample violin plots for carotid tissue proteomes of the plaque zone and marginal zone (*n*=10). Following valid value filtering, data was median normalized, log2 transformed, and imputed from a Gaussian distribution. **B.** Volcano plot of carotid plaque and marginal zone proteome with purple and pink representing plaque and marginal enriched, respectively (q-value<0.05). **C.** Donor uncorrected principal component analysis of label-free proteomics of whole-tissue carotid plaque (purple) and marginal zones (pink; *n*=5). **D.** Percent incidence of Gene Ontology (GO; Biological Process and Cellular Component) terms (false discovery rate <0.05) containing "vesicle" resulting from all differentially expressed proteins (q-value<0.05) from plaque and marginal zones (right panel; *n*=5, repeated from Figure 1C). Left panel lists top 26 (by adjusted p-value) GO Biological Process terms and highlighted are vesicle-associated terms from marginal zones. **E.** Top 25 (by adjusted p-value) GO Cellular Component pathways derived from all differentially enriched proteins in plaque zones. **F.** Top 25 (by adjusted p-value) Kyoto Encyclopedia of Genes and Genomes pathways derived from all differentially enriched proteins in plaque zones. Statistical significance assessed by Fisher's exact test (D)

**Supplemental Figure 3. Marginal zone pathway analysis features vesicle related and atherogenic pathways. A.** Top 25 (by adjusted p-value) Gene Ontology Cellular Component pathways derived from all differentially enriched proteins in marginal zones. **B.** Top 25 (by adjusted p-value) Kyoto Encyclopedia of Genes and Genomes pathways derived from all differentially enriched proteins in marginal zones.

**Supplemental Figure 4. Carotid Plaque EV-enrichment protocol and validation of purity.** **A.** Representative images showing dissection of whole carotid plaque into marginal and plaque zones. **B.** Schematic for EV isolation from atherosclerotic carotid plaque tissue. Briefly, tissues were collected in Hanks balanced salt solution (HBSS) and dissected into plaque and adjacent marginal zones. Subsequently, tissues were prepared for size exclusion chromatography (SEC) by enzymatic digestion, and centrifugation and filtration to remove cellular debris. EVs were enriched via SEC in fractions 7-10 and concentrated using ultrafiltration to a desired volume. EVs destined for proteomics underwent an additional purification step with ultracentrifugation and density gradient ultracentrifugation. **C-D.** Lipoprotein co-isolation in EV preparations via enzyme-linked immunosorbent assay (ELISA) for ApoA1 (HDL; **C**) and ApoB100 (LDL; **D**; *n*=1 plaque). **E.** Fractional analysis of nanoparticle isolation via nanoparticle tracking analysis (NTA) and bicinchoninic acid assay (*n*=1). **F.** Flow Cytometry based assessment of whole cell co-isolation with Hoechst nuclei stain. Bone marrow cells (positive control) and plaque-derived EV fractions depicted on left and right panel, respectively. EV, extracellular vesicle (*n*=1).

**Supplemental Figure 5. Proteomics of EVs enriched from carotid tissue identifies several EV markers. A.** Prototypical EV markers are enriched in fractions 7-10. Abundance Z-score normalized by protein in four samples. **B.** Bubble plot of GO pathway analysis of all differentially expressed EV-proteins between plaque zone and marginal zone filtered by vesicle-related terms. Data points are sized by combined score, and size-scaled by genes altered in pathway/total number of unique genes in analysis and color-scaled by biological processes and cellular locations.

**Supplemental Figure 6. Density gradient ultracentrifugation (DGUC) after size exclusion chromatography (SEC) enriches for EVs and depletes extracellular matrix proteins.** N=3 matched sample pre- and post-DGUC. **A.** After DGUC, 85.4% (958/1121) of proteins IDs remain, 158 protein IDs were no longer identified, and 4 were uniquely identified. Markers of EVs were retained following DGUC. **B.** SEC and DGUC cluster separately in principal component analysis (PCA) after using a generalized linear model to account for matched sample variability. **C.** Extracellular matrix proteins were differentially enriched in samples prior to DGUC (Total: N=206 differentially enriched in SEC pre DGUC, N=58 differentially enriched post DGUC). p-value threshold of 0.1. **D.** Transmission electron microscopy confirms the removal of debris indicative of extracellular matrix and retention of EV structures. Representative image, *n*=3 samples. Scale bars = 200 nm **E.** Post-DGUC nanoparticle tracking analysis identified particles with a size distribution consistent with the TEM imaging in both marginal zone (MZ) and plaque zone (PZ)**.** Graphs shows mean ± SEM. **F.** Proteomic data pre-processing and per-sample violin plots for EVs derived from carotid tissue plaque zone and marginal zone (*n*=46). Following valid value filtering, data was median normalized, log2 transformed, and imputed from a Gaussian distribution.

**Supplemental Figure 7. EV-miRNA transcriptomics data post-alignment quality metrics.** **A.** Alignment breakdown chart features each sample as a column. The percentage of reads with different alignment outcomes (unique paired, non-unique, unaligned) is quantified by the y-axis and visualized by colored stacked columns. **B.** Coverage plot showing average read depth for each sample. **C.** Total reads plot showing total reads for each sample. **D.** Average base quality per read plot displays the proportion of reads with a given Phred quality score wherein all base quantities within the read are averaged. Lines represent individual samples. **E.** Average base quality per position plot displays the average Phred score for each position in the reads. Lines represent individual samples. **F.** Donor uncorrected principal component analysis (PCA) showing EV-miRNA (top panel) and EV-protein (bottom panel) profiles of EVs enriched from carotid plaque zones (PZ, purple) versus marginal zones (MZ, pink) (miRNA *n*=20 pairs; protein *n*=23 pairs).

**Supplemental Figure 8. TISSUES Database identifies known cell and tissues types driving atherosclerotic vascular disease. A-B.** Dot plot of top (by strength metric, FDR<0.05) cell type and tissue annotations derived from inputting all EV-proteins **(A)** and all differentially enriched EV-proteins in marginal **(B,** top panel**)** and plaque zones **(B,** bottom panel**)**. **C.** Proteins used to inform cell annotations (SMC, foam cell, and B-lymphoblastoid cell) in the TISSUES database. Colored nodes represent query proteins and edges represent protein-protein interactions (curated and experimentally validated) with line thickness indicating the strength of data support.

**Supplemental Figure 9. Publicly available carotid scRNA-sequencing data identifies over 17,000 single cells that can be annotated to distinct cell types based on gene signatures. A-C.** Uniform manifold approximation and projection (UMAP) of 17,000 plaque- and marginal-region derived single cells labelled by sample **(A)**, region **(B)**, and cell-type **(C)**. **D-F.** Dot plot of marker gene expression for T cells **(D)**, macrophages and dendritic cells **(E)**, and vascular smooth muscle cells (VSMCs; **F**). Dot size and color darkness indicate the percentage of gene-expressing cells, and average expression, respectively. **G)** UMAP of markers *MS4A1*, *MZB1*, *TPSAB1*, *VWF* for identifying B-cell lymphocytes, plasma cells, mast cells, and endothelial cells (ECs), respectively.

**Supplemental Figure 10. Global atherosclerotic plaque EV communication patterns. A.** Normalized (DeSeq2 median normalization) counts of EV-miRNAs by identity in plaque and marginal zones. **B-D:** Integration with publicly available carotid plaque scRNA-seq dataset. **B.** Stacked bar plot visualizing enriched signaling pathways by overall information flow from global EV-proteome to plaque (purple) and marginal (pink) single cells. Pathways are colored by their dominance within the plaque or marginal regions. **C.** Heatmap depicting EV-cell signaling strength employed by all EV-proteins to atherosclerotic single cells in plaque (right panel) and marginal (left panel) regions. **D.** Chord diagrams representing global EV ligand to EC (left), macrophage (center), and SMC (right) ligand-receptor pairs. Arrows represent EV-receptor to ligands interactions. Individual cell populations annotated in legend. Bar graphs show mean ± SEM. Statistical significance assessed by two-tailed paired t-test (A) with Benjamini-Hochberg false discovery rate correction.

**Supplemental Figure 11. Site specific EV communication strategies in plaque and marginal zones. A-B.** Heatmap depicting differential marginal and plaque EV-cell signaling strength stratified by tissue site; derived from layering differentially enriched EV proteins in marginal **(A)** and plaque **(B)** zones to atherosclerotic single cells in plaque and marginal regions, respectively. Strength of EV-derived signaling molecule and cell type shown on right and top bar graph, respectively. **C.** Chord diagrams representing differentially expressed EV ligand to macrophage ligand-receptor pairs. Arrows represent EV-ligand interactions. Individual cell populations annotated in legend. Marginal and plaque zones represented in left and right panels, respectively.

**Supplemental Figure 12. Further characterization of EV release, contents, and pathway analysis in asymptomatic and symptomatic plaques. A.** Hematoxylin and eosin (H&E) and Movat pentachrome stain of asymptomatic (Patient 3 and 4) and symptomatic (Patient 5) of carotid plaque specimens. Large plaque with significant inflammation, but without a necrotic core (Patient 3). Large, calcified plaque with calcified/necrotic center with intraplaque hemorrhage (Patient 4). H&E staining with intraplaque hemorrhage indicated by green arrowhead and Movat pentachrome staining with intraplaque hemorrhage indicated by cyan arrowhead. Large, calcified plaque with calcified/necrotic center with intraplaque hemorrhage (Patient 5). H&E staining with intraplaque hemorrhage indicated by green arrowhead and Movat pentachrome staining with intraplaque hemorrhage indicated by cyan arrowhead. Scale bar (500 μm) as shown. **B.** Differences plot from the Wilcoxon test comparing asymptomatic and symptomatic plaque versus marginal EV concentrations (Figure 5C-D). **C-D.** Nanoparticle tracking analysis (NTA) of nanoparticle concentration between asymptomatic versus symptomatic cohorts in marginal **(C)** and plaque **(D)** zones. **E.** Protein-protein interaction network with all differentially enriched EV-proteins in asymptomatic and symptomatic plaque versus marginal zones. Represents full STRING network wherein edges indicate both functional and physical protein interactions with a medium confidence interaction score (0.400). **F.** Bar graph of all significant KEGG pathways (adjusted p-value<0.05) derived from shared asymptomatic and symptomatic plaque-derived EV-proteins (adjusted p-value<0.05). Cancer-, and infection- associated pathways were excluded from analysis.

**Supplemental Figure 13. Recipient cell repertoire in asymptomatic versus symptomatic (vulnerable) plaques. A-B.** Box plot of predicted EV-miRNA-mRNA target signature enrichment among cells of the atherosclerotic plaque, informed by integrating differentially detected EV-miRNA - mRNA targets in asymptomatic **(A)** and symptomatic **(B)** plaques with scRNA-seq of carotid atherosclerotic plaques.

**Supplemental Figure 14. Symptom (asymptomatic versus symptomatic) and site-specific (plaque versus marginal) differences in EV-ligand-receptor interactions. A.** Chord diagrams representing ligand-receptor pairs between differentially enriched EV-proteins (ligands) to ECs (receptor) in symptomatic marginal zones. Red arrows represent EV-ligand to receptor interactions. Individual cell populations annotated in legend. **B-D.** Chord diagrams representing ligand-receptor pairs between differentially enriched EV-proteins (ligands) to SMCs (receptor) in asymptomatic plaque zone **(B)**, symptomatic marginal zone **(C)** and symptomatic plaque zone **(D)**. Purple arrows represent EV-ligand to receptor interactions. Individual cell populations annotated in legend. **E-G.** Chord diagrams representing ligand-receptor pairs between differentially enriched EV-proteins (ligands) to macrophages (receptor) in asymptomatic plaque zone **(E)**, symptomatic marginal zone **(F)** and symptomatic plaque zone **(G)**. Yellow arrows represent EV-ligand to receptor interactions. Individual cell populations annotated in legend.

**Supplemental Figure 15. Approach to externally validate cell communication predictions in dataset from the Biobank of Karolinska Endarterectomy (BiKE).** A. CD63 protein expression in symptomatic (S) vs asymptomatic (AS) patients in both central (plaque zone) and distal (marginal zone) regions. B. NAMPT (visfatin) protein normalized to CD63 level by zone in S versus AS patients in both central (plaque zone) and distal (marginal zone) regions. C. VEGFA protein normalized to CD63 level by zone in S versus AS patients in both central (plaque zone) and distal (marginal zone) regions. Protein levels are expressed in arbitrary units (AU). CD63 zone values were determined based on CD63 protein levels in each of the central AS (0.86 AU set to 1.1), distal AS (0.79 AU set to 1), central S (1.20 AU set to 1.5) and distal S (0.96 AU set to 1.2). NAMPT and VEGFA protein levels were then multiplied by the corresponding CD63 zone values for each group to account for CD63 expression in these zones. Statistical comparison was performed using a two-tailed t-test when comparing AS vs S and a paired t-test when comparing distal vs central zones as they were collected from the same patient. P-value threshold for significance was set to 0.05 and results are presented as mean ± standard deviation (SD). There were n=9 patients in the AS group and n=9 patients in the S group.

**Supplemental Figure 16. Bioinformatic and network analysis to delineate distinct and shared functions of EV-miRNAs and EV-proteins. A.** Bubble plots depicting top Gene Ontology pathways (by fold enrichment) for EV-miRNA (left panel), EV-proteins (middle panel), and network analysis (BIONIC network integration algorithm, 512 embedded features, Louvain-clustered modules) to integrate whole EV-secretome (miRNA and proteins; right panel). **B.** Bubble plots depicting top Kyoto Encyclopedia of Genes and Genome pathways (by fold enrichment) for EV-miRNA (left panel), EV-proteins (middle panel), and network analysis (BIONIC network integration algorithm, 512 embedded features, Louvain-clustered modules) to integrate whole EV-vesiculome (miRNA and proteins; right panel). Bubbles colored by q-value and sized by fold enrichment.

**Supplemental Figure 17. Overrepresentation analysis delineates distinct and shared pathways being modulated by plaque EV-miRNA and EV-proteome with symptom status-based distinctions in pathway identity and strength. A-B.** Overrepresentation analysis with Gene Ontology (GO) pathways using differentially expressed predicted EV-miRNA-mRNA targets **(A)** and EV-proteins **(B)** in asymptomatic and symptomatic groups. **C-D.** Overrepresentation analysis with Kyoto Encyclopedia of Genes and Genomes pathways using differentially expressed predicted EV-miRNA-mRNA targets **(C)** and EV-proteins **(D)** in asymptomatic and symptomatic groups.

**Supplemental Figure 18. Overrepresentation analysis finds endothelial and angiogenesis-based pathways as contributors to plaque vulnerability. A-C.** Subset analysis of differentially expressed EV-miRNAs **(A,B)** and EV-proteins **(C)** querying specific endothelial and angiogenesis based KEGG Pathways. Pairwise comparison of mean fold enrichment of EC-specific pathways in symptomatic versus asymptomatic plaques **(B)**. **D.** Subset analysis of differentially expressed EV-proteins querying specific endothelial and angiogenesis-based Gene Ontology Biological Process (GO-BP) Pathways. **E-F.** Subset analysis of differentially expressed EV-vesiculome (miRNA and proteins; Per-module) querying specific endothelial and angiogenesis-based Gene Ontology Biological Process (GO-BP; **E**) and Kyoto Encyclopedia of Genes and Genomes pathways (KEGG; **F**).

**Supplemental Figure 19. Functional validation of increased angiogenesis with plaque-derived vesicles employing a newly optimized low-input sprouting assay. A-B.** Pairwise comparison of mean fold enrichment of endothelial cell (EC)-specific pathways in EV-protein based Gene Ontology (GO; **A**) and Kyoto Encyclopedia of Genes and Genomes (KEGG**; B**) pathways in symptomatic versus asymptomatic plaques. Individual tests labelled above graph. **C-D.** Pairwise comparison of mean fold enrichment of EC-specific pathways in EV-vesiculome (miRNA and protein) based Gene Ontology (GO; **C**) and Kyoto Encyclopedia of Genes and Genomes (KEGG**; D**) pathways in symptomatic versus asymptomatic plaques. Individual tests and p-values labelled above graph. **E.** Schematic of sprouting angiogenesis assay depicting EV-enrichment from plaque and marginal zones of symptomatic patients, treatment of immortalized human umbilical endothelial cell (HUVEC) derived spheroids, followed by staining and quantification. **F.** Representative confocal microscopy images of HUVEC-spheroid angiogenesis after addition of EVs (2-4X$10^{10}$ EVs; 24 h), vascular endothelial growth factor (VEGF; positive control), plaque zone (PZ) elute (PZ EV control), marginal zone (MZ) elute (MZ EV control), assay control (media only). Scale bar represents 100 µm. **G.** Quantification of average sprout length (n=188-331 sprouts from 7–10 spheroids from 3 independent experiments). **H-J.** Quantification of sprout number **(H)**, sprout length **(I)**, and average sprout length **(J)** with addition of VEGF (50 ng/ml, positive control; *n*=6-8). Bar graphs show the mean±SEM. Violin plots show median ± interquartile range. Statistical significance assessed by 1-way ANOVA with

Sidak multiple comparisons test **(**adjusted p-values shown, G). P-values meeting significance (<0.05) shown in bolded text **(G)**.
